# Retrotransposons facilitate the tissue-specific horizontal transfer of circulating tumor DNA between human cells

**DOI:** 10.1101/2022.08.10.501131

**Authors:** Munevver Cinar, Lourdes Martinez-Medina, Pavan K. Puvvula, Arsen Arakelyan, Badri N. Vardarajan, Neil Anthony, Ganji P. Nagaraju, Dongkyoo Park, Lei Feng, Faith Sheff, Marina Mosunjac, Debra Saxe, Steven Flygare, Olatunji B. Alese, Jonathan Kaufman, Sagar Lonial, Juan Sarmiento, Izidore S. Lossos, Paula M. Vertino, Jose A. Lopez, Bassel El-Rayes, Leon Bernal-Mizrachi

**Affiliations:** Department of Hematology and Medical Oncology, Winship Cancer Institute of Emory University, Atlanta; Kodikaz therapeutic solutions, LCC, New York, NY; Bioinformatics group, Institute of Molecular Biology NAS RA, Yerevan, Armenia; Departments of Integrated Cellular Imaging Core, Winship Cancer Institute of Emory University, Atlanta, GA; Division of hematology and oncology O’Neal Comprehensive Cancer Center, University of Alabama at Birmingham, Birmingham, AL; Pathology and Laboratory Medicine, Winship Cancer Institute of Emory University, Atlanta, GA; Department of Computational Biology/ Genetics, The University of Utah, Salt Lake City, UT; Department of Surgery, Winship Cancer Institute of Emory University, Atlanta, GA; Department of Medicine, Division of Hematology-Oncology and Molecular and Cellular Pharmacology, Sylvester Comprehensive Cancer Center, University of Miami, Miami, FL; Department of Biomedical Genetics and the Wilmot Cancer Institute, University of Rochester Medical Center, Rochester, NY; Bloodworks Northwest Research Institute, Division of Hematology, University of Washington School of Medicine, Seattle, WA

**Author notes:** **Corresponding Author:** Leon Bernal-Mizrachi, MD, Winship Cancer Institute, Emory University 1365 Clifton Road, Building C, 3rd floor Atlanta, GA 30322.

## Abstract

A variety of organisms have been shown to have altered physiology or developed pathology due to gene transfer, but mammals have never been shown to do so. Here, we show that circulating tumor DNA (ct) can promote cell-specific horizontal gene transfer (HGT) between human cancer cells and explain the mechanisms behind this phenomenon. Once ctDNA enters the host cell, it migrates to the nucleus and integrates into the cell’s genome, thereby transferring its genetic information. We determine that retrotransposons of the ERVL, SINE, and LINE families are necessary for cell targeting and the integration of ctDNA into host DNA. Using chemically synthesized retrotransposons, we found that AluSp and MER11C reproduced multiple myeloma’s (MM) ctDNA’s cell targeting and integration into MM cells. We also discovered that ctDNA might, as a result of HGT, influence the treatment response of multiple myeloma and pancreatic cancer models. Overall, this is the first study to show that retrotransposon-directed HGT can promote genetic material transfer in cancer. There is, however, a broader impact of our findings than just cancer since cell-free DNA has also been found in physiological and other pathological conditions as well. Furthermore, with the discovery of transposons-mediated tissue-specific targeting, a new avenue for the delivery of genes and therapies will emerge.

## Introduction

The transfer of genes between cells is known to play an important physiological and pathological role in many organisms. In prokaryotes, horizontal gene transfer (HGT) produces changes more impactful in the genome evolution than the branched trajectory^1^. Acquisition of novel traits through HGT can provide prokaryotes with survival and evolutionary advantages against environmental stressors ^2, 3^. HGT in bacteria occurs by multiple molecular mechanisms. Among them, conjugation and transformation take place through mobile genetic elements such as transposons^2, 4–7^.

The exchange of genetic material via transposons is a ubiquitous method for HGT in some eukaryotes such as insects and plants. From the initial discovery of transposon-mediated genetic exchange between rice and millet plants ^8^, much evidence of transposon-mediated HGT has been detected in plants, particularly parasitic plants. Acquisition of the host plant’s genetic material allows parasitic plants to evolve more rapidly to adapt to new and changing environments. In fact, on many occasions, the exact moment of transposon-mediated HGT defines the branch point in the evolution of different versions of the invasive plants ^9^. Similar events of transposon-mediated HGT have been observed in drosophila.

In contrast to the examples above, the evidence in humans for HGT and any potential mechanism involved are less well established. Under physiological conditions, a few studies have shown the relocation of non-gene-coding regions between human cells ^10, 11^. In immunology, early data suggest that the exchange of cell-free DNA from a T cell can elicit the synthesis of antibodies by B cells. In pathological conditions such as cancer, the evidence is even more rare. The discovery that tumor-derived cell-free DNA can contain genetic alterations relevant to tumorigenesis ^12^ led researchers over the last decade to evaluate the possibility that circulating tumor DNA (ctDNA) serves as a vehicle for genetic exchange between tumor cells ^13^. Their results suggest that HGT can occur and alter tumor phenotype by transferring oncogenic genes or reshaping the tumor microenvironment ^14–16^. However, definitive evidence of ctDNA-mediated HGT and the mechanism involved in HGT has been lacking until now.

In this study, we found that tissue-specific retrotransposons mediate the process whereby ctDNA target and transmits genetic material to cells that resemble their cell of origin. These discoveries lay the framework for a new field of study on the role of ctDNA in cell-cell communication and its ramifications in fields such as embryogenesis, cell evolution, cancer, immunology, and therapeutic gene transfer.

## Results

### ctDNA incorporation into tumor cells resembling its cell of origin

To determine whether ctDNA is capable of targeting cancer cell lines, we isolated ctDNA from the plasma of various patients with multiple myeloma (MM), metastatic pancreatic cancer (PC), and colon cancer (CC). First, we verified that nucleic acids extracted from plasma were genomic DNA by showing that DNase I digestion abolished the DNA signal on agarose gel but RNase A or proteinase K did not (Supplemental Figure 1A). Next, we confirmed that the DNA found in the circulation of these patients reflects the cancer patients’ tumor genome. Comparisons of exon and whole-genome sequencing from primary tumors and the ctDNA of PC and MM patients revealed that 100% of the ctDNA mutations correspond to the tumor mutational landscape, but roughly 20% of the tumor mutations cannot be identified by ctDNA sequencing. These results confirm that ctDNA was produced by tumor cells (Supplemental Figure 1B). Based on these findings, we set out to evaluate the possibility that ctDNA targets cancer cells and is capable of translocating into the nucleus. To do so, we introduced rhodamine-labeled ctDNA from patients with MM [n=4], pancreatic [PC, n=3], colon [CC, n=3], and lung cancer [LC, n=1] into the culture media of cell lines that matched the ctDNA’s tumor tissue of origin. The ctDNA localized in the nucleus of all of the corresponding tumor cell types (Figure 1A, Supplemental Video 1A and B) at different levels compared to the control signal of cells alone. We validated this observation using flow cytometry; more than half of multiple myeloma cells (MM1s:52-54%) and almost all pancreatic cancer (MIA: 99%) and colon cancer (HCT116:99%) cells were in contact with ctDNA (Figure 1B). However, after removing plasma membrane-bound ctDNA with trypsin, only 20-30% of the MM1 cells and almost all MIA (99%) and HCT116 (99%) cells remained positive for Cy5 labeled ctDNA. These startling results demonstrate the efficiency of ctDNA in transferring genetic material between tumor cells.

**Figure 1.**
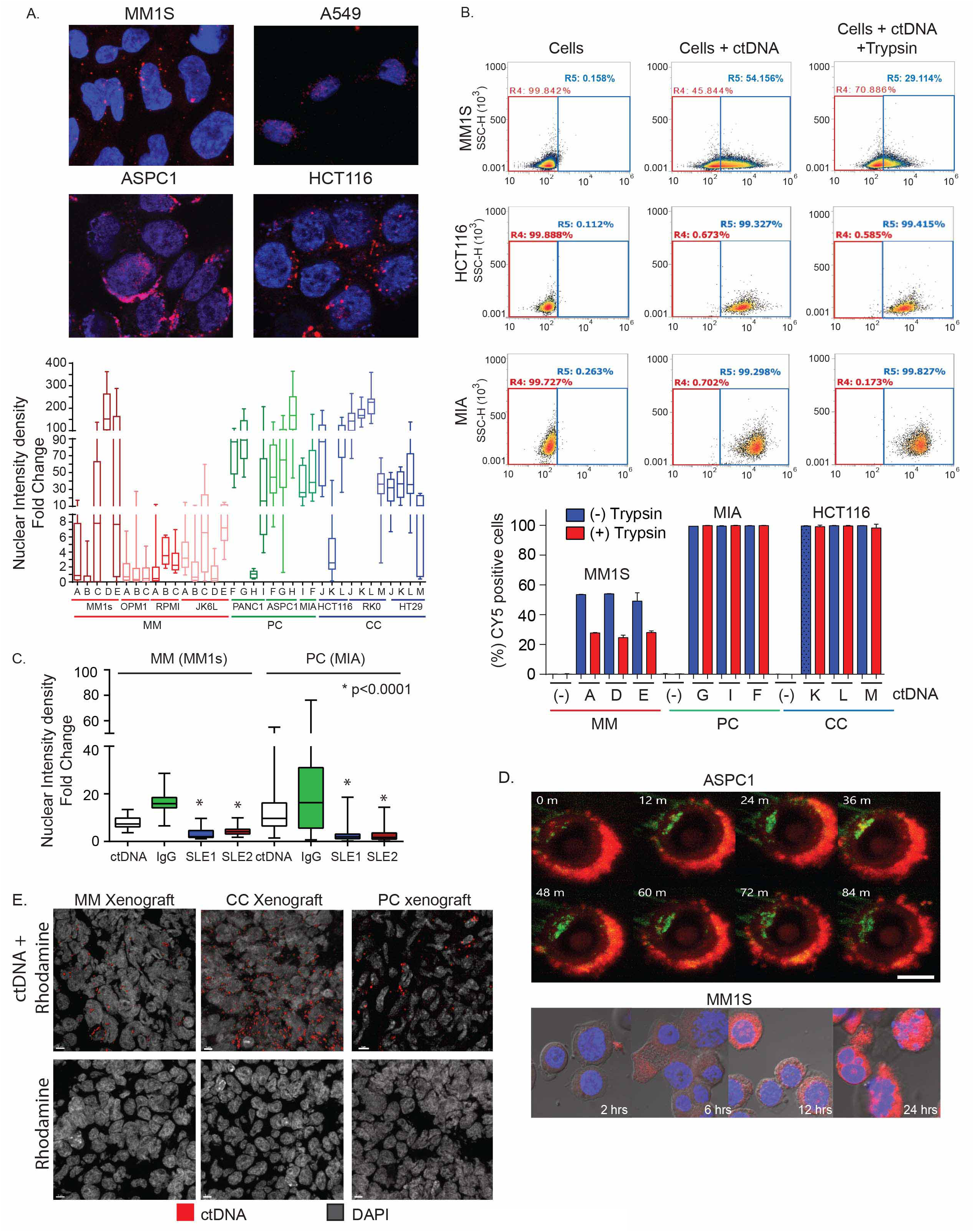
ctDNA incorporation into tumor cells. A. Index images showing the nuclear localization of rhodamine-labeled ctDNA (red) in multiple myeloma (MM1s) and lung (A549), pancreatic (ASPC1), and colon (HCT116) cancer cell lines. The lower panel shows the box and whiskers plot summarizing fold change nuclear density derived from the comparison of cell lines cultured with ctDNA derived from patients with multiple myeloma (n=5), pancreatic (n=3), or colon (n=3) cancer and the background nuclear intensity of untreated control cells. Letters on the x-axis refer to individual patient samples. B. Flow cytometry assay demonstrated a high percentage of cells with ctDNA incorporation. One million myeloma (MM1s), pancreatic (MIA), and colon (HCT116) cancer cells were cultured with CY5-labelled ctDNA (1μg/mL) for 24 hours. To determine ctDNA internalization, cells were then treated for 30 minutes with trypsin (100 uL of 0.25% Trypsin) to remove any ctDNA bound to membrane proteins. (Lower panel) Box plot summarizes triplicate experiments’ data of each cell line cultured with multiple ctDNA samples derived from patient samples. Letters on the x-axis refer to individual patient samples. C. Anti-double stranded DNA antibodies (Anti-dsDNA) reduce ctDNA nuclear localization in multiple myeloma (MM1s) and pancreatic cancer (MIA) cells. Nuclear intensity density fold change of rhodamine-labeled ctDNA in cells after cultured with PBS (control), IgG (control), and anti-dsDNA antibodies from SLE patients 1 (SLE1) and 2 (SLE2). Images were taken 24 hours after coculturing antibody-ctDNA with cells. The nuclear signal was measured using similar methods as described in Figure 1A. D. Time course of cytoplasmic and nuclear localization of rhodamine-ctDNA from patients with pancreatic cancer and multiple myeloma in ASPC-1 (upper) MM1s (bottom) cells, respectively. In ASPC-1 cells, the membrane was labeled with GFP (green), and ctDNA was labeled with rhodamine (red). In MM1s cells time course, the nucleus was labeled with DAPI (blue) and ctDNA with rhodamine (red) E. Tumor localization of rhodamine-ctDNA and rhodamine alone (control) 48 hours after tail injection (representative images from triplicate experiments). Images in all samples were taken with an open red channel for rhodamine detection. MM: Multiple myeloma, PC: Pancreatic cancer, and CC: Colon cancer.

To validate that ctDNA can enter cells, we used monoclonal antibodies against double-stranded DNA (dsDNA) extracted from patients with systemic lupus erythematosus (SLE) that bind to specific DNA sequences ^17^, which we hypothesized would disrupt the interaction of double-stranded ctDNA with a potential cell membrane receptor. We cultured ctDNA isolated from MM and PC patients overnight with two different anti-dsDNA antibodies. Subsequently, the ctDNA-antibody mixtures were exposed to cells for 24 hours. Anti-dsDNA antibodies significantly decreased rhodamine-labeled ctDNA signals in the nucleus (ctDNA nuclear intensity: 3.23-4.2 MM and 2.06-2.8 PC) compared to the IgG isotype control (16.5MM ctDNA and 25.6 PC, p0.0001, Figure 1C) or PBS (7.9 MM and 14 PC). To corroborate the capacity of ctDNA to transport and integrate non-orthologous genetic material into target cells, we ligated a linearized CMV-Green Fluorescent Protein (GFP) vector into the middle of a ctDNA fragment of a myeloma patient and added the product to MM1s cells in culture. We detected a GFP signal in cells cultured with the ctDNA-containing CMV-GFP construct, whereas cells cultured with CMV-GFP alone did not show a GFP signal (Supplemental Figure 1C). These data provide further evidence that ctDNA can mediate HGT between cancer cells.

Next, we investigated the time required for ctDNA to enter the nucleus in multiple myeloma and pancreatic cancer cell lines. In the pancreatic cancer cell line ASPC-1, ctDNA from a PC patient targeted the cell membrane within seconds, internalized into the cytoplasm within minutes, and localized into the nucleus within 10 minutes (Figure 1D and Supplemental Video 2A-C). In contrast, ctDNA from a multiple myeloma patient reached the cell membrane of MM1s cells within 2 hours, internalized into the cytoplasm within 6 hours, and reached the nucleus within 8 hours, with a maximum nuclear localization at 24 hours (Figure 1D).

### ctDNA preferentially migrates to tumor cells in mice

To test *in vivo* the hypothesis that ctDNA targets tumor cells, we developed xenograft models using PC (MIA), MM (MM1s), and CC (HCT-116) cell lines. In a pilot study using a PC xenograft, we injected rhodamine-labeled PC ctDNA or PBS (control) into the tail of the mice. Results determined that the maximum tumor localization of rhodamine-labeled ctDNA was detected at 48 hours post-tail vein injection (Supplemental Figure 1D). We then tested the tumor localization using ctDNA from different cancer patients. Rhodamine-labeled ctDNA isolated from patients with multiple myeloma, pancreatic cancer, and colon cancer (n=3 per tumor type) were injected into the tail-vein of mice bearing the corresponding tumor xenografts (n=3 per tumor type). Mice bearing the same type of tumors were injected with rhodamine alone (n=2) as a control. Tumors and organs (liver, spleen, lung, kidney, colon, and pancreas) were harvested 48 hours after injection, and frozen sections were prepared. Confocal microscopy using a red channel identified a high rhodamine signal in the tumors of mice injected with rhodamine-labeled ctDNA compared to control mice (Figure 1E). No immunofluorescence signal was detected in any organs examined, suggesting that ctDNA preferentially accumulates in tumor cells (Supplemental Figure 1E).

### ctDNA is predominantly incorporated into cells from the same cell of origin

The observation that ctDNA preferentially targets tumor cells raises the possibility that ctDNA has a selective tropism for cells similar to those from which the ctDNA originated. We tested this possibility by coculturing ctDNA with tumor cell lines from tissues distinct from the patient ctDNA tissue of origin. When ctDNAs derived from PC or CC patients were cultured with MM cell lines, the ctDNA clustered on the periphery of the cell membrane and failed to internalize. We observed a similar phenomenon in experiments in which PC and CC cell lines were cocultured with ctDNA extracted from mismatched-tumor-type patient plasma. On the other hand, ctDNA’s nuclear internalization was significantly increased when ctDNA was cocultured with cells matching the ctDNA’s tumor type (Figure s 2A and B). We validated this unexpected finding by simultaneously adding ctDNA from patients that matched or did not match the tumor type of MM (MM1s and JK6L), PC (PANC1 and ASPC-1), and CC (HT29 and HCT-116) cell lines and measuring the nuclear localization of the ctDNA. The ctDNA only accumulated in the cell nucleus if it originated from a patient whose tumor matched the cell line’s tissue of origin (Figure 2C and D). Otherwise, the ctDNA remained at the cell exterior. Finally, 3D reconstruction of the images allowed us to identify that in the infrequent events in which mismatched ctDNA migrated into the nucleus, it colocalized with matching ctDNA (Supplemental Video 3A and B).

**Figure 2.**
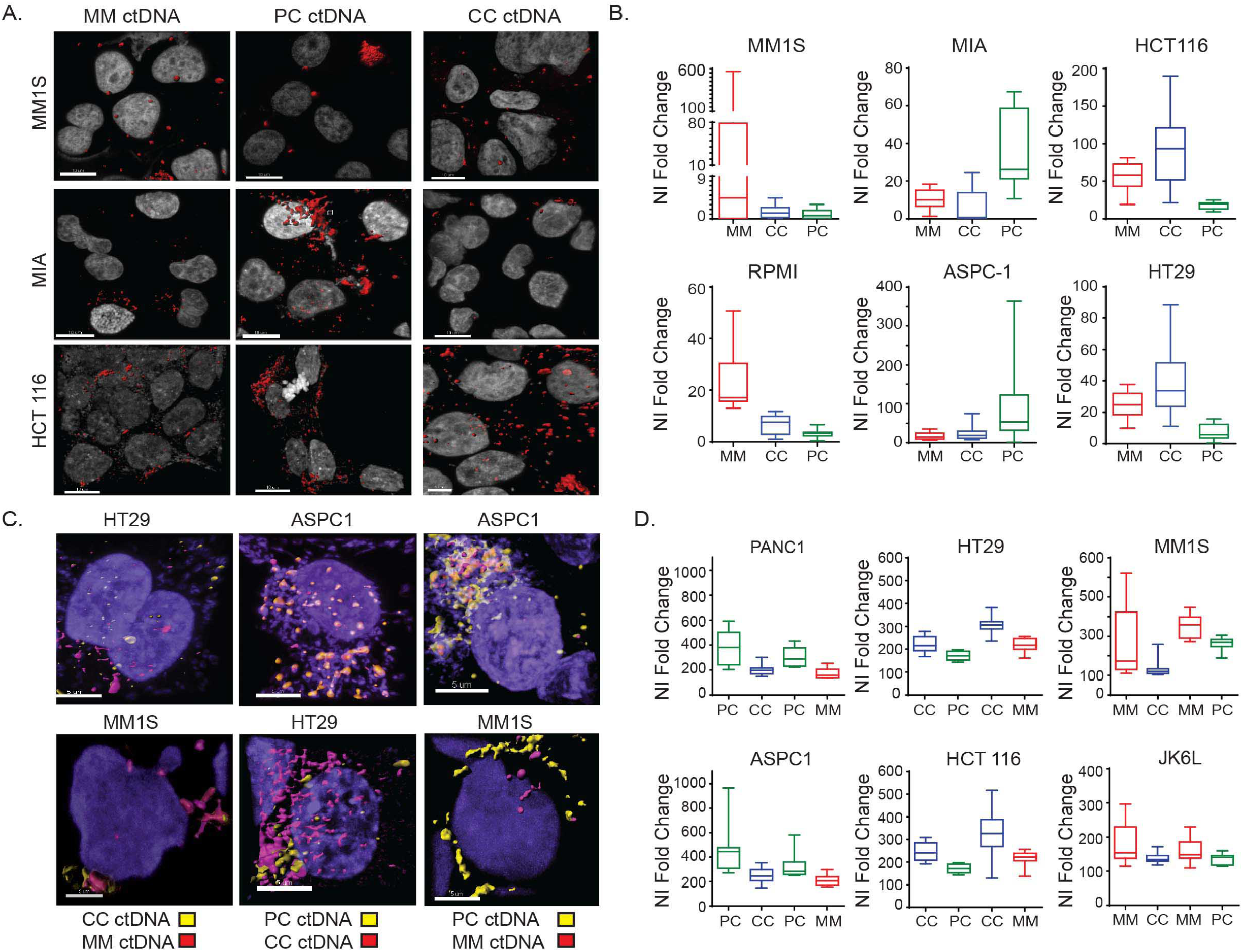
ctDNA cell-specific targeting. A. Index images of 10 experiments and B. fold change of ctDNA nuclear density measurements in cell lines cultured with ctDNA matching or not matching the patient’s cancer type. B. whiskers plot summarizing fold change of nuclear density measurements of cells culture with ctDNA derived from cancer patients matching or not the tumor type of the cell line (n=10 experiments) C. Index images and (D) fold change of nuclear density measurements of simultaneous cocultured of tumor matched and unmatched ctDNA and cell lines (n=10 experiments). MM: multiple myeloma, CC: colon cancer, and PC: pancreatic cancer. MM cells: MM1S, RPMI, JK6L, PC cells: ASPC-1, PANC1, MIA, and CC cells: HT29 and HCT 116. Error bars in the box and whiskers plot identified the standard deviation of triplicate experiments.

We then studied whether ctDNA displayed cell-specific tropism in vivo in 2 xenograft models (MM and PC). Rhodamine-MM and CY5-PC-labeled ctDNA were simultaneously injected into the tail veins of each animal (n=2). Microscopy of the tumors demonstrated that rhodamine-ctDNA from MM patients accumulated more in MM xenografts than in PC xenografts. In contrast, CY5-labeled PC ctDNA accumulated in the PC xenograft but not in MM xenografts (Supplemental Figure 1E). These data demonstrate that *in vivo* as *in vitro*, ctDNA selectively targets cancer cell types similar to its cell of origin.

### Chromosomal integration of ctDNA

We carried out metaphase chromosomal spreads of MM, PC, and CC cell lines after they were cocultured with ctDNA matching the tumor cell type (N=3 per tumor type) to test if ctDNA fragments are capable of integrating into the cell genome. Data showed rhodamine-ctDNA bands incorporated in multiple chromatids (Figure 3A and Supplemental Figure 2A). The number of chromatids with integrated ctDNA appears to vary depending on the cell line (Figure 3B).

**Figure 3.**
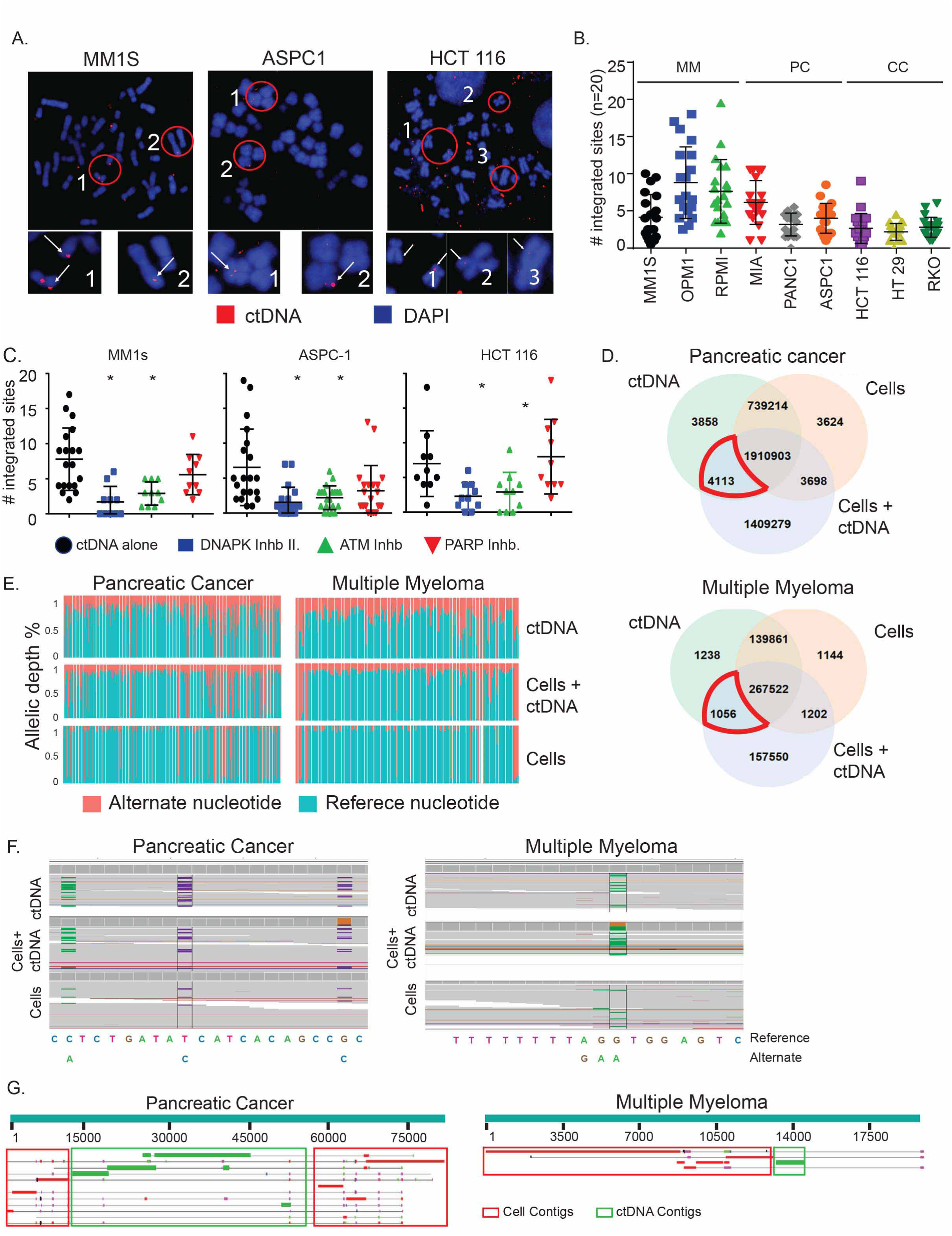
Integration of ctDNA is mediated by non-homologous end joining (NHEJ) repair and integrases. A. Immunofluorescence index images of ctDNA (red) integration into chromatids (blue) in MM (MM1S), PC (ASPC-1), and CC (HCT116) cell lines. Circles define zoomed regions of interest. White arrows identify an area of ctDNA integration. B. Scatter plot displaying the number of chromatids with ctDNA integrations in different MM (MM1s, RPMI, OPM1), PC (MIA, PANC1, ASPC-1), and CC (HCT 116, HT29, RKO) cell lines. Error bars indicate standard deviations of triplicate experiments (n=20 metaphases per experiment). Color and symbol shapes signify different cell types. C. Incorporation of rhodamine-ctDNA fragments into chromosomes of multiple myeloma (MM1s), pancreatic cancer (ASPC-1), and colon cancer (HCT 116) cell lines after treatment with NHEJ (DNAPKcs and ATM), and PARP (n=10). Cells were pretreated for 2 hours with inhibitors of DNA-PK (inhibitor II), ATM (KU-55933), and PARP (NU1025) before the addition of rhodamine-ctDNA to the cell culture. The number of integrated sites was measured in 10-20 metaphases per condition. D. Gain of nucleotide variants in cells cocultured with ctDNA. Comparative SNV analysis between cell genome, ctDNA, and coculture. Venn diagram displays exclusive and shared SNVs between each experimental condition. The area marked in red highlights SNVs commonly shared between ctDNA and coculture condition ctDNA/cell. E. Stacked bar diagram demonstrating the changes in allele depth of the variant (red) and reference (Blue) allele in multiple myeloma ctDNA, cell line genome, and coculture condition. Cells under coculture conditions have more depth in the varian allele in several locations compared to the control cell genome. F. Figure s demonstrating two index IVG variant calls images and their allele frequency in multiple myeloma and pancreatic cancer experimental conditions. Green horizontal bars (Adenine) and blue (guanine) are, in these cases, the alternate nucleotide, and gray bars are the reference nucleotide. G. Blast images demonstrate the transition point of insertion between cell genome contigs (red boxes) and ctDNA contigs (green boxes). Results were obtained after comparing the contigs carrying insertions in the coculture condition with the reference cell genome. Coculture contigs carrying a ctDNA insertion were identified using NucDiff analysis. MM: Multiple myeloma, PC: Pancreatic cancer, and CC: Colon Cancer.

The capacity of ctDNA to integrate into the genome was further defined using DNA repair inhibitors. Myeloma (MM1s), pancreas (ASPC-1), and colon (HCT 116) cancer cells were treated for 2 hours with the inhibitors of DNA damage response (i.e., ATM inhibitor (KU-55933), DNA-dependent protein kinase catalytic subunit inhibitor (DNA-PK inhibitor II), and an inhibitor of alternative join repair, such as poly (ADP-ribose) polymerase (PARP, NU1025)]. Following treatment with these inhibitors, ctDNA was added to the culture medium, and chromosome spreads were performed 24 hours later. Inhibition by the ATM and DNA-PKcs inhibitors significantly reduced the integration of ctDNA into the genome compared to untreated cells. In contrast, PARP inhibition had little effect on ctDNA integration in most cells (Figure 3C and Supplemental Figure 2B). These results could not be explained by changes in the viability of the cells after treatment with the different inhibitors (Supplemental Figure 2C). Overall, these data further demonstrate the capacity of ctDNA to integrate into the host cell genome and suggest cellular pathways that mediate ctDNA 1integration/ligation.

These results prompted us to analyze the ctDNAs sequences integrated into the cell genome. We performed a comparison analysis of whole-genome sequences from 2 cell lines, MM1s (multiple myeloma) and MIA (pancreatic cancer) cell lines, ctDNA from the patients used to generate the data in Figure 3B, and cell lines after being cocultured for 24 hours with the isolated ctDNA. First, we evaluated whether single nucleotide variants (SNVs) unique to the ctDNA could be identified in the coculture condition. Nucleotide variants were identified by comparing the sequence from each experimental condition after alignment to the reference human genome (Hg38). Multiple SNVs were identified in common between coculture conditions and ctDNA that were not present in the genome of the untreated control cells (here called “SNVs of interest,” Figure 3D). Further detailing of these SNVs of interest demonstrated that cells cocultured with ctDNA showed skewing of the variant allele frequency towards that of the variant ctDNA allele compared to cells alone (Figure 3E). Further examples of SNVs exhibiting ctDNA ’skewing’ in the variant allele frequency (VAF) after coculture with ctDNA are shown in Figure 3F and Supplemental Figure 3A.

Next, we used the sequence data to reconstruct genomic contigs for each experimental condition by de novo assembly. Comparison analysis of the contig sequences between the cell line, ctDNA, and coculture condition indicated that ctDNA fragments were integrated into the cell genome under coculture conditions. Blast analysis further validated the gain of ctDNA fragments in coculture conditions and identified the transition points of ctDNA insertion (Figure 3G and Supplemental Figure 3B). Thus, these orthogonal analytical approaches confirm the introduction of ctDNA fragments into the cell genome under coculture conditions. Analysis of incorporated ctDNA fragments and genome-insertion sites identified from both tumor models indicated that most (∼67%) of the inserted ctDNA fragments originated from chromosomes 3 and 7. Moreover, 80% of the ctDNA fragments targeted cellular chromosomal regions near the genomic location from which ctDNA originated (Supplemental Spreadsheet 1). The remaining 20% were inserted into a chromosomal location distinct from their site of origin. These findings are consistent with the higher genetic recombination rate between homologous regions ^18, 19^.

Next, we explored whether inserting the ctDNA fragments enriched any particular pathway. Due to the small number of insertions, the gene ontology analysis of the tissue-specific inserted segments failed to attain statistical significance (Supplemental Figure 4). Gene ontology analysis of the tissue-specific inserted fragments failed to reach significance due to the low number of insertions utilized (Supplemental Figure 4). However, several trends of pathways enriched by MM and PC inserted fragments. The phosphatidylinositol metabolic process, microtubule and calmodulin binding, double-stranded RNA binding, and mitochondrial protein-containing complex were among the enriched pathways in MM. The pathways for cell junction assembly, cell morphogenesis regulation, cell cortex, histone deacetylase complex, and protein tyrosine phosphatase activity, among others, tended to be enriched in PC inserted pieces (Supplemental Figure 4).

### Retrotransposons are located at the 5ʹ and 3ʹ ends of the inserted ctDNA fragments

Transposable elements (TE) play an essential role in HGT in prokaryotes and a few eukaryotes ^20, 21^. Hence, we designed experiments to identify the presence of transposons in ctDNA fragments. To map ctDNA genomic junctions more precisely, we ligated a PACBIO probe to multiple myeloma and pancreatic cancer ctDNA samples to label their 5ʹ and 3ʹ ends. We then performed whole-genome sequencing using the Illumina platform. De novo assembly was performed on the sequencing output. ctDNA contigs containing the PACBIO adaptor were then matched to those contigs containing inserted ctDNA sequences identified in the coculture experiments. RepeatMaster^22^ detected and classified the transposable element on ctDNA sequences at the insertion points (<100 nucleotides from transition point between cell and ctDNA) ^22^. This analysis demonstrated that the ctDNA fragments integrated into the cell genome contained more transposable elements than non-integrated ctDNA fragments (Supplemental Table 1). These findings suggest that retrotransposons mediate horizontal gene transfer in cancer cells, as they do in prokaryotes and plants.

Next, we searched for transposons that preferentially target cells of matching tumor types. Hence, we compared the list of transposons identified at the ctDNA insertion point in matching and mismatching coculture conditions (i.e., MM ctDNA and MM cells or PC cells) and selected transposons that were uniquely inserted in the matching conditions. Interestingly, class I transposons, ERV-Ls, LTRs, SINEs, and LINEs, comprised the majority of transposable elements at the transition points in pancreatic cancer or multiple myeloma. Within matching PC ctDNA, AluSx, MIRc, and MTL1J were some of the most common retrotransposons subfamilies located at insertion points. In multiple myeloma, AluSp, MER11, MER11C, AluJb, and L2a, among other subfamilies of retrotransposons, were identified at the MM ctDNA insertion sites (Figure 4A and Supplemental Table 2).

**Figure 4.**
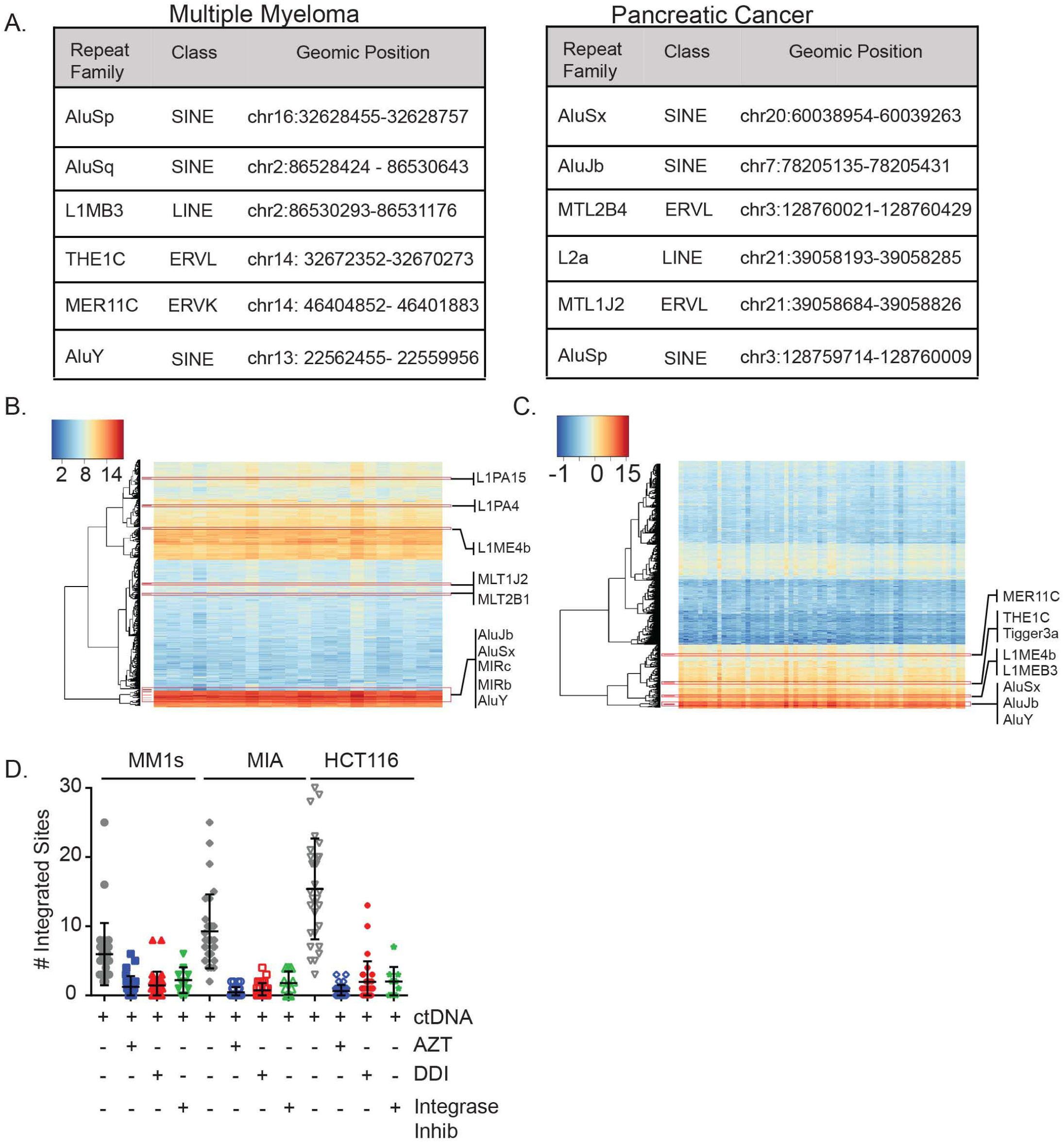
Transposon-mediated horizontal gene transfer of ctDNA in cancer cells. A. Tables summarizing the list of tissue-specific transposons identified at the ctDNA insertion sites. B. Expression levels of selected TE in PC and MM tumor samples. D. Scatter plot displaying the number of chromatid with rhodamine-ctDNA integration of MM (MM1s), PC (ASPC-1), and CC (HCT 116) cell lines (n=30) after treatment with two reverse transcriptase inhibitors (AZT and DDI) and an integrase inhibitor (raltegrabir). Cells were pretreated, and analysis was performed using similar protocols as in Figure 3A. MM: Multiple myeloma, PC: pancreatic cancer, and CC: Colon cancer.

Activation of short interspersed nuclear elements (SINEs) and long interspersed nuclear elements (LINE) has been observed in cancer, leaving open the possibility of retrotransposition. We, therefore, evaluated the expression of transposable elements in RNA from cancer cells from 60 MM and 23 PC patients. To this end, batch normalized raw data was used to measure the expression variability (methods section). The analysis confirmed several tissue-specific retrotransposons to be highly expressed across pancreatic cancer or multiple myeloma. These findings demonstrate that the retrotransposons identified at insertion sites were being transcribed in these cancer types (Figure s 4B and C).

To evaluate the role of retrotransposons in horizontal gene transfer of ctDNA, we analyzed the impact of inhibiting reverse transcription or integrases on ctDNA integration. Therefore, before adding the ctDNA to the culture media, we treated the multiple myeloma (MM1s), pancreatic cancer (MIA), and colon cancer (HCT116) cell lines with the reverse-transcriptase inhibitors zidovudine (AZT) or didanosine (DDI), as well as with the integrase inhibitor raltegravir ^23^. When compared to the no-treatment control, the use of the reverse transcriptase and integrase inhibitors significantly decreased ctDNA chromatid integration (Figure 4D and Supplemental Figure 2A), giving proof that retrotransposons are crucial for ctDNA integration into the host cell’s genome.

### Chemically synthetized retrotransposons target cells of similar tumor origin

To determine if the retrotransposons identified above can transfer genetic material between cancer cells, we chemically synthesized several MM retrotransposons containing point mutations unique to the tumor type and tested their HGT capacity. To this end, we selected among the list of retrotransposons those that can be synthetized via the gBlock gene fragment method (AluSp, AluSg2, MER11C, THE1A, and AluSx). Before any cell line experiment, we evaluated whether they were able to resist degradation by DNAses present in complete culture media. A synthetic sequence with similar length and GC content that does not encode a retrotransposon and a PC retrotransposon (AluSx1) were used as controls. Agarose gel electrophoresis demonstrated that all fragments remain intact under culture conditions (Supplemental Figure 5A). Following these results, we labeled these retrotransposons with CY5 and evaluated their capacity to target and internalize in MM cell lines. First, we evaluated the kinetics of retrotransposon capture by flow cytometry. Our results showed that cell capture of CY5-AluSp increases over time more efficiently than the capture of the control sequence (Supplemental Figure 5B). Subsequently, we performed dose titration experiments to evaluate cell capture and internalization. We considered the DNA internalized if it was not removed from the cells with trypsin treatment. Cell capture and internalization CY5-AluSp were dose-dependent and more efficient than the capture of the control sequence (Supplemental Figure 5C). Based on these findings, we compared the efficiency of cell capture of all transposons in two MM cell lines at 4 hours of culture. CY5-AluSp and CY5-MER11C were captured more efficiently than the other MM transposons, the control sequence, or PC transposons (Figure 5A and Supplemental Figure 5D).

**Figure 5.**
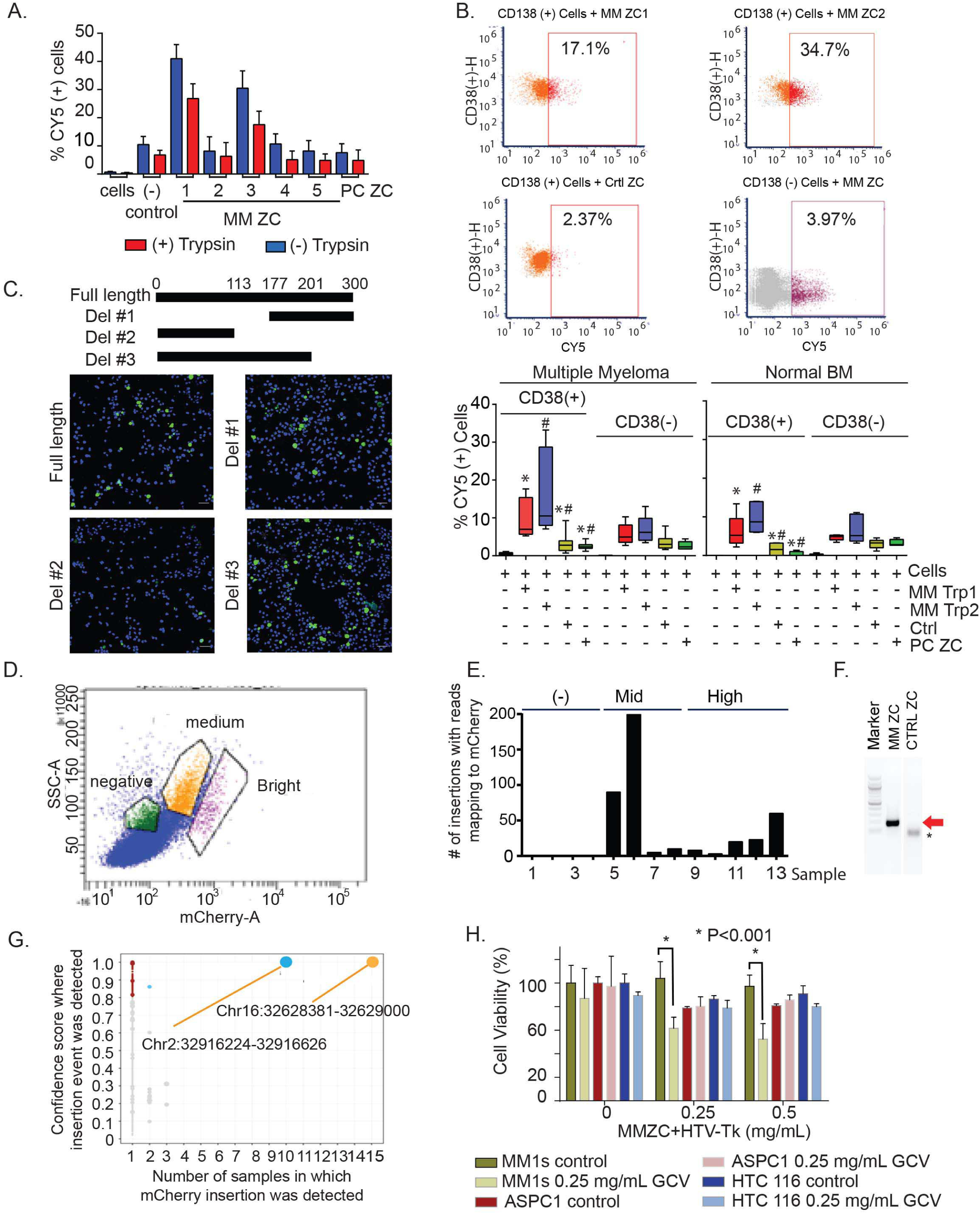
MM transposons mediated horizontal gene transfer to cancer cells. A. MM1s cell capture an internalization of different multiple myeloma retrotransposons and controls after 4 hours of culture. B. Index flow cytometry images displaying the cell capture of AluSp, MER11, PC transposon (AluSx) and control sequence in plasma cells and non-plasma cells derived from bone marrows of patients with newly diagnosed or treated multiple myeloma patients (n=12) and patients with various cancers other than multiple myeloma (n=5). The lower part of the image display box plots summarizing the number of CY5 positive cells measured after 14 hours of culture of twelve multiple myeloma and five non-myeloma bone marrows. C. Effect in MM1s cell internalization of Cy5 labeled AluSp and its deletion mutants. Images were captured after culturing cells with the retrotranspsons for 8 hours. D. flow cytometry image displaying different degrees of mCherry expression in cells cultured for 24 hours with AluSp-CMV-mCherry cassette. E. Bar chart displaying the summary of the number of insertions identified in mCherry expressing (+) and non-expressing (-) cells. F. Validation of the mCherry integration by PCR from chromatin extracted from cells treated with AluSp-CMV-mCherry cassette or control-CMV-mCherry cassette. *nonspecific band. G. Graph displays the confidence of detection of CMV-mCherry insertions identified versus frequency of each specific site of insertion. H. Cell survival of 3 dierent cell lines cocultured with TE-HSV-Tk-GFP for 24 hours prior to adding ganciclovir (GCV). Apoptosis was measured at 96 hours after GVC addition. MM: multiple myeloma, CC: colon cancer and PC: pancreatic cancer, TE-CMV-GFP: transposon element joining to CMV-GFP, HSV-TK: herpes simplex virus thymidine kinase. Error bars in box and whiskers plot identi-ed the standard deviation of triplicate experiments.

The cell targeting of AluSp and MER11C was then measured in bone marrow samples from patients with multiple myeloma or without multiple myeloma but with other bone marrow disorders. After 14 hours of culture with marrows derived from myeloma patients, we observed more CY5-positive plasma cells (CD 38 [+]) in marrows cultured with CY5-AluSp or -MER11C than with control or PC sequences (Figure 5B). In contrast, AluSp or MER11C did not exhibit a difference in CD38 (-) cell capture compared to control or PC sequences. As further evidence of their ability to specifically target malignant plasma cells, we found that AluSp and MER11C-CY5 capture was more effective in CD38 (+) cells than in CD38 (-) cells (AluSp CD38 [+]: 9.9% 1.6% vs CD38 [-]: 5.2% 0.8% P=0.03 and MER11C: CD38 [+]: 16.50% 3.8 vs CD38 [-]: 7% 1.2% P=0. However, there was no distinction between CD38 (+) and CD38 (-) cells in control or PC sequence cell capture. Experiments with the bone marrow of non-myeloma patients demonstrated a similar tendency of greater CD38 (+) cell capture by AluSp and MER11C compared to controls or PC sequences, with the trend disappearing in CD38 (-) cells. Lastly, when we compare the uptake of malignant CD38 (+) cells to non-malignant CD38 (+) cells, we see a non-significant increase in the AluSp and MER11C uptake of malignant CD38(+) cells.

To test whether the retrotransposon sequence contains a cell recognition signal, we generated several deletion mutants of the AluSp sequence and evaluated their impact on cell capture. As shown in Figure 5C, deleting the 3’ end of AluSp reduced MM cell capture, suggesting that the last 230 based pairs contain an MM cell recognition sequence. Multiple sequence alignments (MSA) of AluSp and all MM-specific transposons revealed several conserved nucleotides and two GC and AT-rich regions towards the 3’ end region (Supplemental Figures 5E and F). Having determined that AluSp was able to target MM cells specifically, we evaluated whether it could deliver other genetic material into cells. For this, we ligated AluSp sequences to both ends of a linearized CMV-mCherry cassette. These AluSp-mCherry ligates were then added to the culture media of MM1s cells for 24 hours, after which mCherry expression was evaluated microscopically. More AluSp-mCherry–treated cells expressed mCherry than cells incubated with the linearized vector or the circular CMV-mCherry vector (Supplemental Figure 5G). Flow cytometry also detected mCherry expression in MM1s cells (Figure 5D). To determine whether the mCherry cassette integrated into the MM1s genome, we sequenced the genomes of single cells expressing different levels of mCherry. mCherry insertions were identified by recognizing sequences with one read aligned to the cell genome while the mate aligned to the CMV-mCherry sequence. This analysis detected various mCherry insertions in cells with mid or high expression levels, while no insertions were identified in cells without mCherry expression (Figure 5E). In contrast, when cells were cultured with control-mCherry sequence, only a few were found with high levels of mCherry insertion and expression. mCherry integration was confirmed by mCherry PCR of cells cultured with AluSp-mCherry vector but not control-mCherry cultured cells (Figure 5F). The AluSp-CMV-mCherry vector integrated high confidence into two specific regions in the genome (Chr2:32916224-32916626 and Ch16:32628381-32629000), which are enriched for simple repeats and AluSp, respectively (Figure 5G).

After demonstrating that AluSp can deliver a gene, we investigated whether this property can be exploited therapeutically. We ligated a herpes simplex virus thymidine kinase (HSV-Tk) killer gene between AluSp sequences (AluSp-HSV-Tk-GFP) and tested this vector’s ability to kill MM, PC, and CC cell lines. Cells were cultured for 24 hours with the AluSp-HSV-Tk-GFP and then cultured for 96 hours in ganciclovir (GCV). At that point, we measured the effect on cell numbers of the biologically active byproduct of GCV produced by the action of HSV-Tk. As shown in Figure 5H, GCV markedly reduced the viability of MM cells previously treated with MM-AluSp-HSV-Tk. In contrast, AluSp-HSV-Tk/GCV treatment did not affect the viability of PC or CC cell lines. These results demonstrated that the synthetic AluSp could transfer and integrate genetic cargo into specific cells (Figure 5H).

### ctDNA alters the drug response of MM or PC cell lines

Having demonstrated horizontal gene transfer in cancer cells, we explored potential consequences of transfer on the phenotype of target cells, including on the target cell’s response to drugs. We therefore cultured Bortezomib-sensitive (BS) MM cell lines (MM1s and OPM1) for 24h with DNase-treated or non-treated serum from bortezomib-resistant (BR) patients or control serum from non-cancer patients. Subsequently, increasing doses of bortezomib were added to culture media, and cell survival was measured 24 hours later. Compared to control plasma, plasma from patients resistant to bortezomib increased the bortezomib resistance of MM1s and OPM1 (Figure 6A). This effect of BR plasma was eliminated by pretreating with DNase (Figure 6A). When BR MM cell lines (JK6L and RPMI) were cultured with a plasma of patients sensitive to bortezomib, the cells exhibited a significant restoration of bortezomib sensitivity compared to when the cell lines were cultured with control plasma. DNAse treatment of BS plasma abolished this effect to levels similar to control plasma (Figure 6B). Similar experiments were performed with PC cell lines (MIA Paca-2 [MIA] and PANC1) using DNase-treated gemcitabine (GEM)-resistant (GR) and control plasma. Both cell lines became resistant to GEM when cultured with GR plasma but not with control plasma. DNase-treatment of the GR plasma abrogated the effect (Figure 6C). In contrast, DNase treatment of control plasma resulted in increased resistance of PC cells to gemcitabine (Figure 6C, lower graphs).

**Figure 6.**
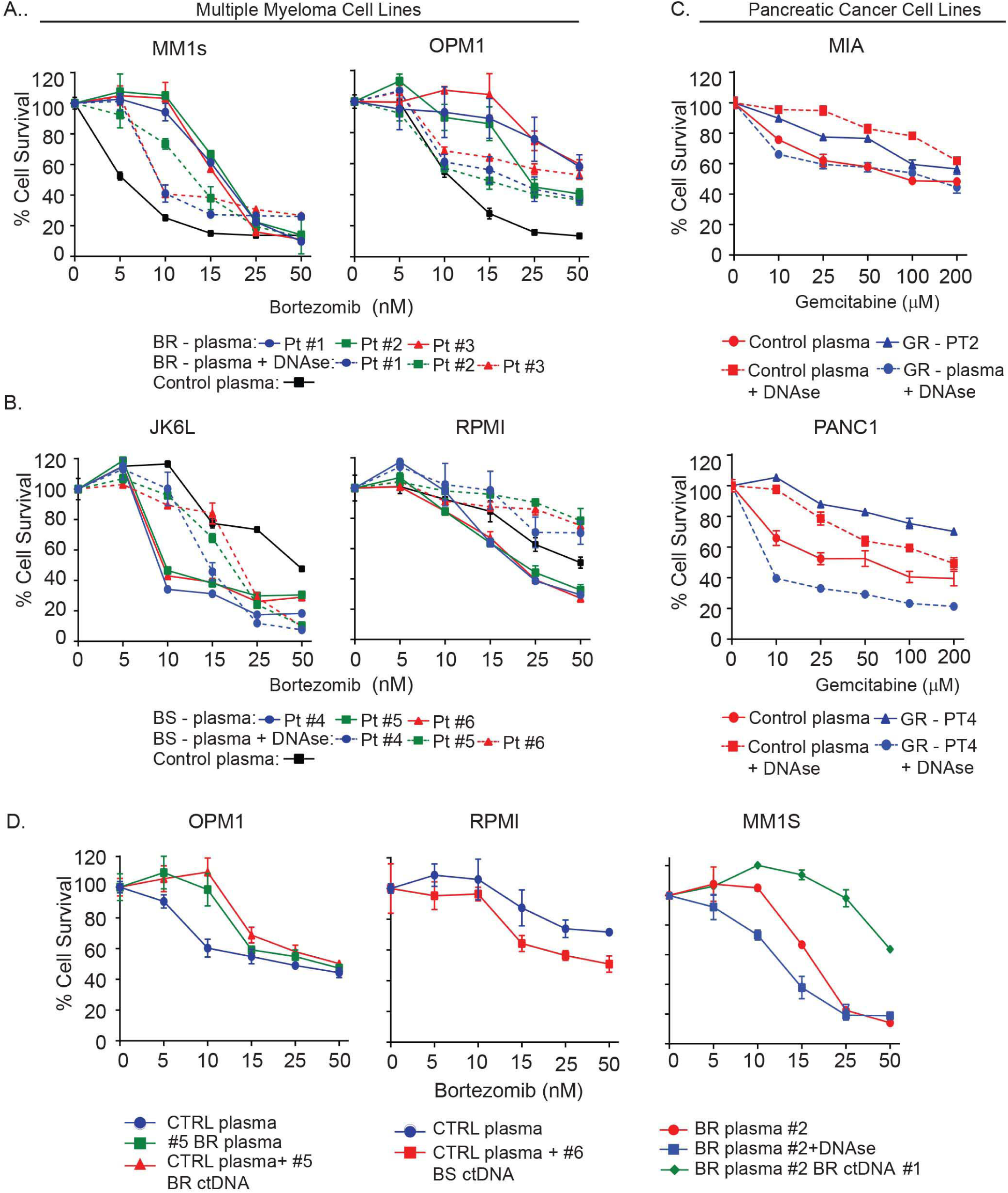
ctDNA-mediated transmission alters the sensitivity of the cells to bortezomib or gemcitabine. A. Cell viability assay measuring sensitivity to bortezomib in OPM1 and MM1s cells cultured with control plasma derived from non-cancer patient or DNAse I treated or non-treated plasma from patients that failed to respond to bortezomib. B. Cell viability assay measuring sensitivity to bortezomib in RPMI and JK6L cells cultured with control or DNase I-treated or non-treated plasma from a patient that responded to bortezomib-based treatment. C. Cell viability assay measuring sensitivity to gemcitabine in pancreatic cancer cell lines (MIA and ASPC-1) cultured with plasma from patients resistant to gemcitabine, similar plasma pretreated with DNAse I or a control non-cancer patient. For the corresponding DNAse I-treated samples, plasma was treated with DNase I for 10 minutes. D. (Left), Comparison of cell viability response to bortezomib in OPM1 cells cultured with plasma from a patient that failed to respond to bortezomib treatment, the combination of control plasma with ctDNA derived from the same patient resistant to bortezomib and control plasma alone (non-cancer patient). (Middle). Viability response of RPMI to control plasma or control plasma with added ctDNA obtained from a patient that achieved a complete response to bortezomib. (Right) Cell viability assessment after bortezomib treatment of MM1s cells cultured with plasma from a bortezomib resistant patient (BR#2) alone or after treatment with DNAse I or coculture with ctDNA from a different bortezomib-resistant (BR#1) patient. MM: multiple myeloma and PC: pancreatic cancer. BR: Bortezomib resistant, BS: Bortezomib sensitive, and GR: Gemcitabine resistant. Error bars indicate the standard deviation of triplicate experiments.

To further characterize the contribution of ctDNA transferring drug response, we added cell-free ctDNA from bortezomib-resistant or -sensitive patients to media containing plasma of a patient without cancer or BR. The plasma was then incubated with MM1 or OPM1 cells. Both cell types lost sensitivity to bortezomib (Figure 6D). Resistance to bortezomib was increased even further when the cells were incubated with the plasma of a BR patient to which BR patient ctDNA had been added (Figure 6D). In contrast, adding the ctDNA of a BS patient to RPMI cells cultured with control plasma from a non-cancer patient resulted in increased sensitivity to bortezomib (Figure 6D). These findings suggest that ctDNA transmits genes that confer drug sensitivity or resistance from one cell to another.

## Discussion

Horizontal transmission of TEs plays a significant role in the evolution of prokaryotes and some eukaryotes. However, the role of HGT in humans has not been clearly described. We have discovered that naked ctDNA can serve as a vehicle for transferring genes between cancer cells. Beyond defining the fundamentals of ctDNA HGT, we discovered a previously unknown feature of ctDNA: its preference for cells that are similar to those from which it originated. The identification of retrotransposons at ctDNA insertion sites, the reduction of ctDNA insertions by inhibitors of retrotransposition, and the results showing that synthetically-generated retrotransposons can deliver payloads to target cells support the notion that retrotransposons have an essential role in mediating HGT in cancer. Furthermore, our experiments using deletion mutants of the retrotransposon delineated that these elements contain the address that specifies the delivery of ctDNA to a specific target cell. Importantly, these results are laying the groundwork for research into the use of a synthetic Transposon-mediated Gene Transfer (TGet) to precisely transfer “cargo” to particular cells.

There is limited data available in humans to suggest that some cells transfer information through the release of naked double-stranded DNA^11, 24^. Two groups have explored the role of ctDNA as a messenger of genetic material in cancer *in vivo* models over the last decade ^11, 14, 15^. Garcia-Olmo and colleagues suggested that plasma from colon cancer patients is capable of promoting tumor development in a murine embryo fibroblast cell line (NIH-3T3). More importantly, this team showed that plasma from CC patients transmitted oncogenes, including K-Ras, into NIH-3T3 cells ^15, 25^. These findings were later supported by Trejo-Becerril and colleagues, who observed the transfer of the human K-Ras oncogene from SW480 xenograft to chemically-induced colon cancer tissue in rats ^14^. These results align with our studies demonstrating the transfer of genetic material from a patient’s plasma to cancer cell lines. We demonstrated that ctDNA integrates into host cells using whole-genome sequencing, GFP expression in cells cocultured with a ctDNA-CMV-GFP cassette, and mCherry integration in cells treated with ctDNA-derived retrotransposons. Thus, we provide solid evidence that ctDNA functions as a vehicle for gene transfer and is consistent with our hypothesis that ctDNA can play a role in altering the genetic architecture of the tumor mass. More importantly, our observations demonstrating the tissue-specific tropism of ctDNA provide a potential mechanistic explanation for the studies of Trejo-Becerril ^14^, which showed the transfer of mutated human K-RAS from colon cancer cell lines grafted in mice carrying dimethylhydrazine-induced colonic tumors. However, our data falls short of identifying the mechanisms by which ctDNA binds cells and is internalized. Hence, further work elucidating the cellular mechanism for ctDNA recognition will be essential for developing inhibitors to prevent the transmission of genetic material between cancer cells as a means of spreading drug resistance.

The mechanism by which cell-free DNA transfers information between tumor cells has remained elusive ^14, 15^. The evidence that HGT in bacteria, flies, and plants is primarily mediated by specific mobile elements ^26^ suggested the possibility that a similar mechanism occurs between cancer cells. We uncovered multiple lines of evidence for the central role of retrotransposons in ctDNA HGT: 1) genomic analysis identified the presence of retrotransposons at insertion sites and their expression in tumor samples; 2) both integrase and reverse-transcriptase inhibitors reduced ctDNA integration into host cells; 3) synthetic MM retrotransposons were captured by both MM cell lines and plasma cells derived from patients, and 4) MM transposons were able to deliver and integrate mCherry and HSV-TK genes into MM cell lines. Nevertheless, it remains unclear why transposons can perform tissue-specific HGT. The process of retrotransposition is known to be reactivated in cancer cells ^27–30 47^. Thus, it is conceivable that upon arrival at the host cell, a particular transposon carried by ctDNA co-opts the cellular retrotransposition machinery to integrate into the cancer cell genome. Hence, their tissue-specific integration may be determined by the retrotransposon’s sequence and the host’s expression of the retrotransposition machinery ^28^.

Cell-free DNA is incorporated into immune complexes in SLE, serving as an antigen recognition signal ^31^. Specifically, a DNA sequence within the immune complex is what immune cells and antibodies recognize ^32, 33^. In an analogous fashion, our work suggests that a particular sequence within the retrotransposon serves as a recognition signal for target cancer cells. Indeed, we demonstrated that pretreatment of ctDNA with SLE antibodies reduces the number of cells capturing labeled ctDNA in MM and PC cell lines. Furthermore, we demonstrated that a synthetic Alu sequence targets and delivers cargo into tumor cells and that deletion of the 3ʹ end of the Alu disrupts cancer cell targeting, highlighting that certain retrotransposons contain a cell-targeting signal. However, not every ctDNA fragment is captured by the target cell, suggesting that the recognition sequence is not present in each ctDNA fragment. We proved this by demonstrating that different retrotransposons identified at insertion sites displayed variable cell targeting efficiency in MM. These findings together highlight the presence of specific sequence signals that define the tropism for cell targeting, here called “zip code sequence.”

Our experiments showing that pretreating cells with trypsin prevent the cellular uptake of ctDNA and that trypsin treatment after culturing the cells with ctDNA reduces the number of ctDNA-positive cells strongly suggest the presence of a plasma membrane protein receptor for the cell-targeting sequence. This putative receptor remains unidentified. Hence, this work generates new questions to be addressed, including identifying consensus sequences for cell targeting, determining which genetic and phenotypic characteristics of the target cell regulate recognition by ctDNA, and identifying the membrane protein receptor(s) for ctDNA.

The high efficiency by which 2 of the MM transposon sequences target plasma cells while largely sparing other bone marrow cells opens the possibility of exploiting these transposons as therapeutic vectors. Similar to our demonstration of specific delivery of HSV-Tk into MM cells, these transposons could also deliver drugs, radioisotopes, nanoparticles, or killer genes.

Given the genomically instability of cancer cells, we were surprised by the frequency with which the ctDNA fragments were inserted into homologous chromosomal locations. It is known that accurate replication and repair of DNA ensure the proper functioning of a cell. Mechanisms such as homologous recombination play a vital role in this process, particularly between regions with identical sequences ^34^. Therefore, ctDNAs would be expected to highjack this machinery to integrate in genomic regions corresponding to those from which they originated. Hence, the sequence of the ctDNA can, to a substantial degree, define the directionality of insertion.

The relevance of HGT in cancer is unknown. Our data demonstrating that ctDNA derived from patients changes the drug response phenotype of a cell line is somewhat surprising. The possibility of transmitting cancer through blood transfusion is a controversial subject of investigation. In 2007 a large study from Scandinavia (n=888,843) of cancer-free transfusion recipients demonstrated an increased rate of cancer development in years after transfusion compared to the expected incidence rate ^35, 36^, suggesting the possibility that certain blood units contain substances that promote the development of occult cancers. Similarly, other studies have demonstrated an increase in cancer incidence after transfusion; however, the specificity of particular cancer remains a matter of discussion ^37^. Although some studies indicate that transfusion has immunosuppressive effects and the presence of cytokines that can cause carcinogenesis ^38^, it is likely that ctDNA in some transfused units could potentially promote cancer development.

In summary, we report horizontal gene transfer between human cells capable of altering the phenotype of recipient cells. The mechanism involves the secretion of the transposon and accompanying genomic DNA from donor cells into the extracellular space from where it eventually reaches the blood. From the blood, the ctDNA can reach the target cell, which is usually a cell of the same tissue origin as the donor cell. On the target cell, a membrane receptor (likely a protein) recognizes a specific address sequence in the transposon region of the ctDNA and internalizes the ctDNA. The ctDNA travels to the nucleus and becomes integrated into the genome, usually in a region homologous to the site of ctDNA origin, in a mechanism involving reverse transcriptase and integrases. This is the first definitive report of horizontal gene transfer between human cells, and the work raises at least as many questions as it answers. Yet to be determined are the mechanism of ctDNA secretion, the nature of the specific receptor on the target cell, the mechanism of internalization, the cGAS-cGAMP-STING host response ^39^, and other mechanistic aspects. Our findings also raise the possibility that genetic information can be transferred between individuals, through blood transfusion, for example. Finally, the ability to specifically target certain tumor cells potentially adds a powerful new weapon to treat cancer and other diseases.

## Methods

### Clinical specimens and sample preparation

We obtained retrospective plasma samples from 10 multiple myeloma (MM), ten pancreatic cancer (PC), three colon cancer (CC), and two lung cancer (LC) patients from samples stored in the Tissue and Acquisition Bank of the Winship Cancer Institute of Emory University. All patients provided written informed consent approving the use of their samples under Institutional Review Board approval. The ten patients with newly diagnosed MM were treated with bortezomib-containing regimens (among them were five responders and five non-responders). Of the ten PC patients, seven with advanced staged cancer were treated with gemcitabine (at the time of obtaining the plasma sample, two patients were in partial response, and five had progressive disease), and three early-stage patients had undergone surgical resection. Response in MM was determined using the International Uniform Response Criteria for Multiple Myeloma, and PC patients were evaluated using the Response evaluation criteria in solid tumors (RECIST) criteria. Plasma was isolated after centrifuging blood samples at 1500 rpm for 10 minutes. The plasma supernatant was collected for storage. Plasma from non-cancer patients was purchased from Innovative Research, MI.

Bone marrow samples for the coculture of transposons were obtained from fifteen MM patients at various statuses of the disease (four newly diagnosed, three patients post-bone marrow transplant, and eight patients at relapse) and five patients without myeloma (2 patients newly diagnosed with myelodysplastic syndrome, two patients post-treatment of acute myeloid leukemia and one patient with diffuse large b cell lymphoma without marrow involvement)

### Cell lines and reagents

The following cell lines were grown in Roswell Park Memorial Institute (RPMI) 1640 medium: Four multiple myelomas (OPM1, RPMI, JK6L, and MM1S); one PC (ASPC-1); and one LC (A549). Colon cancer cell lines (HCT-116and HT-29) were cultured in McCoy media. Pancreatic cancer cell lines (PANC1 and MIA Paca-2 [MIA]) were cultured in DMEM media, and CC cell line RKO was cultured in EMEM. All culture media were supplemented with 10% fetal bovine plasma, 1% L-glutamine, one mM sodium pyruvate, and 50 μg/ml penicillin-streptomycin. In MIA cells, 2.5% horse serum was also added to the culture medium.

Bortezomib, gemcitabine, ganciclovir, and DNaseI were purchased from Sigma-Aldrich. RNase A, 4, 6-diamidino-2-phenylindole (DAPI), CellLight Plasma Membrane-GFP, Bacman 2.0, and Platinum Polymerase High Fidelity PCR Kit were purchased from Thermofisher-Scientific. DNA-PK inhibitor II and Raltegravir were purchased from Santa Cruz Biotechnology, KU-55933 (ATM kinase inhibitor) from Selleckchem, and NU1025 (PARP inhibitor VI) from Calbiochem. Quick Ligation Kit was purchased from New England Biolabs. Proteinase K, QIAamp MinElute ccfDNA Kits protocol, and QIAmp DNA Blood Mini Preb Kit were purchased from Qiagen. Cell TiterBlue Cell Viability Assay Kit was purchased from Promega. Label IT Cx-Rhodamine and Cy5 were purchased from Mirus Bio. pLenti-III HSVtk Lentivirus vector was purchased from Applied Biological Materials (abm).

### Cell viability

Cell viability assays were performed in 96 well, black, clear-bottom microplates. For MM cell line studies, 3×10^4^ cells were cultured for 24 hours with media containing 10% human plasma of a bortezomib resistant or -sensitive patient or a control non-cancer patient. Cells were then cultured for 24 hours with titrating concentrations of bortezomib (doses: 0, 5, 10, 15, 50 nM, Sigma). For PC cell line viability studies, 1×10^4^ cells were incubated for 24 hours with media containing plasma of a gemcitabine-resistant patient or non-cancer patient. Subsequently, titrating concentrations of gemcitabine (doses: 0, 10, 25, 50, 100, 200 µM, Sigma-Aldrich, MO) were added to the culture media, and cells were then incubated for 96 hours. CellTiter-Blue Cell Viability Assay was used to evaluate cell viability according to the manufacturer’s instructions (Promega). Cell viability was measured by a fluorescent protocol in a microplate reader. Experiments were performed in triplicate of 3 independent studies.

### ctDNA extraction and immunofluorescence labeling

Isolation of ctDNA was performed following the QIAamp MinElute ccfDNA Kits protocol (Qiagen, Cambridge, MA). The ctDNA was fluorescently labeled with either Cx-rhodamine or CY5 using the Label IT^®^ Nucleic Acid Labeling kit (Mirus Bio LLC, WI).

### Image acquisition

One x 10^6^ cells cultured in 1 ml of culture media were incubated with rhodamine- or CY5-labeled DNA at different time points. The methodology used to perform immunofluorescence in suspension cell culture was conducted as previously described ^40, 41^. Adherent cells (2.5 x 10^5^ cells) were grown on a coverslip prior to processing. After cells were attached to slides, slides were washed with PBS twice, and the cells were then counterstained with 4, 6-diamidino-2-phenylindole (DAPI) for nuclear detection (ThermoFisher MA). For live-cell imaging, the plasma membrane was labeled following the cellLight Plasma Membrane-green fluorescent protein (GFP), Bacman 2.0 protocol (ThermoFisher MA). The presented images are from triplicate experiments. Images were acquired using a Leica SP8 LIGHTING confocal microscope housed in the Cell Imaging and Microscopy Shared Resource, Winship Cancer Institute of Emory University.

Lattice light-sheet microscopy was used to obtain live images and movie acquisition presented in Supplemental Videos 1-3. Images were acquired using a 3i v1 Lattice Light Sheet microscope in sample scanning mode, with Δs of 0.8 µm and 71 steps, and a 20 µm x-dither scan of the lattice pattern created with a 0.550 outer NA / 0.500 inner NA annuli. Volume data were collected using a Hamamatsu ORCA-Flash 4.0 v2 via a Semrock FF01-446/523/600/677 blocking filter for both 488 nm and 560 nm laser channels (5% and 10% power, respectively) every 3 mins for 1 -2 hours. Raw data was deskewed using 3i SlideBook 6 software to create correctly orientation volumetric data. The 3D visualization, surfaces, and movies were created in Bitplane Imaris 9. Isosurface settings were user selected for each dataset to represent signal boundaries efficiently.

### Nuclear localization quantification and image analysis

For quantification of the nuclear localization of rhodamine- or Cy5-ctDNA, we obtained 10 images per sample in fields with a minimum of 10 cells. Volumetric data sets were acquired using a Leica SP8 confocal microscope. All data were acquired with the same x, y, and z sampling and with the same xy zoom. Z-stack total heights were varied to encapsulate the thickness of the randomly selected field of view. Data were analyzed using Fiji ^24^, ilastik ^25^, and Matlab computational software (MathWorks Inc, MA). A Fiji macro was used to convert raw.lif files as required; ilastik machine learning models were trained and then applied to classify specific nuclear morphologies; and Matlab was used to process resulting probability maps and quantify rhodamine- or Cy5-DNA signal within the nuclei. Further information about image processing and quantification of the ctDNA nuclear localization is available at GitHub repository (https://github.com/nranthony/nuc_ctDNA_process). The nuclear intensity fold change value was calculated by measuring the nuclear intensity produced by the rhodamine or Cy5 labeled ctDNA over the background intensity of the nuclear signal in control cells.

### Chromosome spreads and ctDNA banding identification

Rhodamine-labeled ctDNA from patients with MM (n=3), PC (n=3), and CC (n=3) was added to culture media. Briefly, 1 x 10^6^ cells cultured in 1.5 mL of medium were incubated with 1 μg/mL of rhodamine-ctDNA. At 24 hours, the cells were transferred to a 15 ml tube and incubated in 10mL of medium containing 15 µL Colcemid (10μg/mL) at 37°C for 20 minutes before harvesting. After centrifugation and media removal, cells were resuspended with 10 mL of pre-warmed 0.075M KCl and incubated at 37°C for 20 minutes. Two mL of fixative (3:1 methanol: acetic acid) were added and incubated for 10 minutes before centrifugation and aspiration. Samples were then resuspended in 10mL of fixative and incubated at room temperature for 10 minutes, followed by 2 additional washes with a fixative. Slides were prepared in a Thermotron chamber where temperature and humidity were controlled for optimum metaphase spreading. Serial micro-pipetting was performed, 3 µL at a time, until at least 25 cells were visible per field at 20x magnification. After drying slides at room temperature for 1 hour, nuclei were stained with DAPI plus antifade reagent, and coverslips were applied to slides. Cytogenetic technicians performed the readout. Ten to twenty metaphase nuclei were counted per experiment, with touching and overlapping cells excluded. The number of chromosomes with integrated rhodamine bands was counted.

### Assessment of ctDNA integration with non-homologous end-joining repair, the alternative pathway, and transposase inhibitors

To investigate the mechanisms involved in ctDNA integration into the chromosomes of MM1s, ASPC-1, and HTC116 cells, 1 ×10^6^ cells were treated with inhibitors of the non-homologous end-joining (NHEJ) repair system. The inhibitors were ataxia-telangiectasia mutated (ATM) inhibitor KU-55933 (10μM, Santa Cruz Biotechnology, TX) and the DNA-dependent protein kinase, catalytic subunit (DNA-PKCS) inhibitor I (30 μM, Sigma-Aldrich, MO). In addition, we used inhibitors for alternative NHEJ repair pathways, also known as microhomology-end joining, such as the poly ADP ribose polymerase (PARP) inhibitor NU1025 (200 μM, Sigma-Aldrich, MO) and the transposase inhibitor raltegravir (100nM, Sigma-Aldrich, MO). After 2 hours post-drug treatment with inhibitors, rhodamine-labeled ctDNA was added to the culture media and incubated for an additional 24 hours. Cell growth was then arrested, and chromosome spreads were performed as noted above. The number of rhodamine-ctDNA integration sites for each cell was determined by counting a minimum of 10 cells in metaphase. The ctDNA band integration identification and integration counts was performed by personnel from the cytogenetic laboratory of Emory University.

### Xenograft experiments

Mice were housed in a clean facility with an ambient temperature of 65-75°F, 40-60% humidity, and 12 light/12 dark cycles. All experiments included male and female animals. The protocols followed were approved by the Emory University Institutional Animal Care and Use Committee and compliant with ethical regulations for studies involving laboratory animals.

We performed pilot and validation xenograft studies to evaluate the accumulation of ctDNA in tumors in mice. For the pilot time-course study, three mice bearing pancreatic cancer MIA cell xenografts were generated by injecting 1×10^6^ cells in the dorsum of J:NU (007850) outbred nude mice. After tumors reached a volume of 0.5 cm, mice were assigned to a specific experimental arm: 2 mice underwent tail vein injection with rhodamine-labeled ctDNA, and a third mouse received a tail-vein injection with PBS as a control. Tumors from tail vein ctDNA-injected mice were harvested 24 and 48 hours post-injection. For mice in the control group, the tumor was harvested 48 hours post-injection. Based on the results of these experiments, we selected a harvest time point of 48 hours post-injection. For the validation study, xenograft models were developed using human-derived PC (MIA), MM (MM1s), and CC (HCT-116) cell lines. Following the pilot study protocol, three mice per tumor xenograft were then dosed to assess tumor localization of labeled ctDNA. At harvest time, tumors and selected organs (liver, lung, large bowel, pancreas, and spleen) underwent frozen section dissection. Each slide was fixed with paraformaldehyde 4% and stained with DAPI before mounting the coverslip.

### Whole-genome sequencing and Whole exon sequencing

ctDNA was extracted from ten MM and ten PC patients using the methods described above. DNA from CD138(+) cells and from cell lines used in *in vitro* experiments (MM1s and MIA) was extracted using the Blood & Cell Culture DNA Mini Kit (Qiagen, MD). DNA from fresh frozen paraffin-embedded pancreatic tumors was obtained using QIAamp DNA FFPE Tissue Kit Print (Qiagen, MD).

After extraction, ctDNA was ligated to the PACBIO adaptor (GCGCTCTGTGTGCT) using the ABM DNA Library Prep Kit for Illumina Sequencing (Applied Biological Materials Inc. Canada). PACBio-labeled ctDNA and regular ctDNA were subjected to standard methods for library preparation and sequencing using Illumina and Agilent protocols, respectively. Applied Biological Materials Inc. prepared the libraries and performed whole-exon and -genome sequencing. The average target coverage was 50X.

### Nucleotide variance concordance between tumor and ctDNA

#### Quality Control and Alignment To Reference Genome

The raw sequence data was subjected to quality control checks using FastQC (https://www.bioinformatics.babraham.ac.uk/projects/fastqc/). Samples that failed the QC checks were trimmed using BBDuk (https://jgi.doe.gov/data-and-tools/bbtools/bb-tools-user-guide/bbduk-guide/) in the adapter trimming mode for paired reads. The sequence reads were then aligned to the human genome GRCh38 assembly (https://www.ncbi.nlm.nih.gov/assembly/GCF_000001405.39) using the Falcon Accelerated Genomics Pipeline (https://github.com/falconcomputing/falcon-genome). This is an accelerated version of the GATK Best Practices Pipeline. Beginning with paired-end FASTQ sequence files, the first step mapped the sequences to the reference. The resulting mapped BAM file was sorted and duplicates marked. We ran the GATK 4.1.3 best practice somatic mutation pipeline with base recalibration, with read orientation filtering to account for damage seen when using FFPE samples and (https://www.intel.com/content/dam/www/programmable/us/en/pdf/literature/wp/wp-accelerating-genomics-open1-fpgas.pdf, https://www.intel.com/content/www/us/en/healthcare-it/solutions/genomicscode-gatk.html and https://pdfs.semanticscholar.org/e85d/4f927d91e9f25b7c20de71f91c78250771bb.pdf).

#### Variant Calling And Annotation

Variant calling was done using VarDict (https://academic.oup.com/nar/article/44/11/e108/2468301), a novel and versatile variant caller for next-generation sequencing in cancer research. VarDict was chosen based on the recommendations from Sandmann et al ^42^. We used an allele frequency threshold of 0.01. Variants for 6 of the samples (for which a control samples were not available) were called in single sample mode (https://github.com/AstraZeneca-NGS/VarDict). For the samples where controls were available, paired mode was run in order to distinguish between somatic and germline variants. The called variants were annotated using SnpEff (http://snpeff.sourceforge.net/SnpEff.html), which is a variant annotation and effect prediction tool. We used SnpEff’s pre-built GRCh38.86 database for the annotations.

#### Analysis for concordance of ctDNA and primary tumor sample

Single nucleotide variant concordance between ctDNA and corresponding tumor samples was analyzed using bcftools isec (http://samtools.github.io/bcftools/bcftools.html) to obtain concordant variants.

### De Novo Assembly and identification of ctDNA tissue specific sequences

To identify insertion in the coculture conditions, we used two different approaches for assembling the genome. In approach 1, reads from ctDNA (5 MM and 5 PC, cell lines, and cells cocultured with matched and mismatched ctDNA samples were assembled using ABySS ^43^ de novo assembler. Before assembly, ct-DNA samples were 10x depth normalized with bbnorm [5]. The best k-mer size for the assembly was predicted and selected with the KmerGenie tool ^44^, (https://jgi.doe.gov/data-and-tools/software-tools/bbtools/). In addition, ct-DNA samples used in cell line cultured experiments (772 and P201812-2) were assembled without any read depth normalization.

ctDNA contig-level assembly sequences from MM and PC samples were used for cluster analysis with the aim of identifying tissue-specific sequences. cd-hit-est-2d was used to select contigs specific for MM and PC ^45^ with 95% of identity as a threshold. cd-hit-est-2d compared to sequence datasets (data base 1 [db1]=MM and data based 2 [db2]=PC) and reports sequences that are not similar in db2 as well as sequences that are similar between db1 and db2. Since we were interested in MM- and PC-specific contigs, we performed cd-hit-est-2d twice: first, MM contig assemblies were assigned as db2 (for MM-specific contigs), and then PC contig assemblies were assigned as db2 (for PC-specific contigs).

In approach 2 whole-genome sequences (WGS) from ctDNA (5 MM and 5 PC, cell lines, and cells cocultured with matched and mismatched ctDNA samples were used to assemble genomes using SPAdes 3.12.0 ^46^. Prior to de novo assembly, ct-DNA samples were 10x depth normalized with bbnorm (https://jgi.doe.gov/data-and-tools/software-tools/bbtools/). The best k-mer size for the assembly was predicted with the KmerGenie tool ^44^. In addition, ct-DNA samples used in cell line cultured experiments were assembled without any read depth normalization. After this step human mitochondrial DNA was removed using 2 strategies. First, sequence data was aligned to human genome build 38 with the Burrows-Wheeler aligner ^47^ and mitochondrial specific reads were removed. Second, sequence reads were assigned to human mitochondrial genome using Centrifuge (https://ccb.jhu.edu/software/centrifuge/manual.shtml) and a custom database.

### Detection tissue specific insertions

Approach 1. The first method aligned all cell line samples to the human reference genome(hg38) with bwa-mem software (v 0.7.17) ^48^. After the alignment step de novo insertions detection was performed with the Pamir tool ^49^. Pamir uses “one-end anchor” reads (i.e. one end is mapped while the other is unmapped around the breakpoint location) and orphan reads (read pairs where none of the ends can be mapped to the reference) to characterize the novel sequence contents and their insertion breakpoints. Algorithm steps include de novo assembly, re-alignment, and clustering on mentioned reads to generate contigs for putative novel insertions. Aligned BAM files for cell culture sequences along with the hg38 genome reference were supplied to Pamir. The tool outputs a VCF file with the sequence, location, and length of identified novel insertions.

To select cancer type-specific inserts, full-length contigs were converted to BLAST databases. Then, insertions identified in the match-, mismatch- and no-culture samples for corresponding cell line samples (MM1S or MIA) were blasted against the corresponding sample full-length assembly contig database (MM ctDNA for MM1S cells and PC ctDNA for MIA cells). An insert was considered cancer-specific if: 1) it was present in the matching coculture sample but not in mismatch coculture and no-culture samples; 2) it was aligned to the corresponding ct-DNA sample database with an identity at least 70% (to maximize hits for further processing).

To define if the selected unique contigs are present in all samples, we have created a BLAST database for all samples and blasted the unique contigs against each database. The myeloma database consisted of contigs from five MM patients; for pancreatic cancer the database consisted of ten PC patients. Unique contigs from each cancer type were blasted against the corresponding database. Then, we selected the contigs, with alignment length >= 650bp and BLAST identity ≥90%. As a result, we have defined the unique contig sets, which were present in all samples belonging to one cancer type (MM1S - 14 contigs and MIA - 13 contigs). In the next step, we have aligned contigs in all samples with the corresponding insertion, using MAFFT [10], and constructed the consensus sequences.

Approach 2. For the second approach, we used NucDiff ^50^ to identify subsequence repeats, deletions, or novel insertions by comparing WGS assembled de novo. NucDiff uses NUCmer, delta-filter, and show-snps programs from MUMmer ^51^ for comparing closely related sequences. For both MM and PC, we made the following comparisons: a) MM ctDNA cocultured with MM Cells with the MM cells’ sequence; b) MM ctDNA sequence from the same patient used for the coculture experiment with the MM cells’ sequence; c) MM ctDNA cocultured with PC cells; d) MM ctDNA with the PC cells sequence; and e) PC ctDNA coculture with PC cells with the PC cells sequence, PC ctDNA sequence from the same patient used for the coculture experiment with the PC cells, and PC ctDNA coculture with MM cells. For each comparison, we set the parameter “minimum length of a maximal exact match” to 250 (-l) and the minimum cluster length to 200 (- c) to optimize the running time and detect large structural differences between the genomes.

The “struct.gff” and “snp.gff” output files from NucDiff was parsed to identify events that represent transfer of DNA from the ctDNA sample to the cell in the coculture experiment. For each event, we required a) minimum length of >100 base pairs, and b) that the structural event identified with reference to a specific cell contig should be present in both the coculture and ctDNA with an exact match of the insertion sequence with the ctDNA assembly. Inserted sequences from the qualifying structural events in were aligned to the human genome build 37 using Bowtie2 ^52^ to identify genomic co-ordinates of the ctDNA sequence that inserts into the cell. Genes that harbor inserted ctDNA sequences were annotated using genomic co-ordinates from build 37 of the human genome. Enrichment of gene-ontology pathways terms were tested in the identified genes using the R clusterProfiler (https://bioconductor.org/packages/release/bioc/html/clusterProfiler.html).

### Identification of transposable elements and its nucleotide variants

To determine the locations of transposable-like regions in the contigs, sequences were analyzed, and transposable elements (TEs) were identified and classified using RepeatMasker version 4.1.0^22^ and . The Dfam database (release 3.1 ^53^) of repetitive DNA families was used as a reference for identifying repeats in ctDNA contig sequences that were part of qualifying structural events as described above. For each repeat sequence identified by RepeatMasker, we computed the overall frequency of the specific repeat (e.g., for AluSp or L1) and their parent class (e.g., SINE).

Once the TE elements were identified, we aligned hit repeats/TE sequences, unique contigs for all samples, and insertion sequence with MAFFT software version 7. Mutations in TE sequences were identified by comparison of nucleotides at each position in the multiple sequence alignment with the “Biostrings” R package (version 4.1). We considered a position to contain mutation if the substitution was present in contigs of all samples. In the case of ambiguous nucleotides in contigs introduced by short read alignment (putative heterozygosity in the sample), non-matched nucleotide was considered as a mutation if it was present in all samples. Finally, based on the identified mutations, we have constructed the mutated transposon sequences for each unique contigs (for MM1S and MIA datasets).

### Transposon linearized vector

A polynucleotide comprising sequences corresponding to the transposon that contained mutations shared by all the MM samples was generated by Integrated DNA Technologies, Inc (IDT). The sequence of these oligos are described in the supplemental methods section. Similar methods were use to generate deletion mutants from AluSp.

### Statistical analysis

Two side student-T test was use as statistical analysis method for evaluating the difference between ctDNA nuclear localization among cell, the number of base gain in match and mismatch coculture sequencing experiments, and the number of ctDNA integrations in chromatids under the different experimental conditions. Statistical analysis for transposon enrichment was performed using Chi-square test as noted above.

## Data Availability

Sequencing data used for producing Figure 3C-D, 4B-C and Supplemental Figure 1B2 and 5A-D is available under Figshare portal: https://figshare.com/account/home#/projects/87485.

For image processing and quantification of the ctDNA nuclear localization see GitHub repository for the algorithms: https://github.com/nranthony/nuc_ctDNA_process.

## Acknowledgments

We thank Dr. Ravi Majeti for their critical review and comments. LBM was supported by Georgia Research Alliance venture development award. ISL is supported by grant 1R01CA233945 from the National Cancer Institute, the Dwoskin, and Anthony Rizzo Families Foundations and Jaime Erin Follicular Lymphoma Research Consortium. Research reported in this publication was also supported in part by the Winship Shared Resource of Winship Cancer Institute of Emory University and National Institutes of Health (NIH)/National Cancer Institute under award number 2P30CA138292-04. The content is solely the responsibility of the authors and does not necessarily reflect the official views of the National Institute of Health. Also Imaging Core was supported in part by PHS Grant UL1TR000454 from the Clinical and Translational Science Award Program, National Institutes of Health, National Center for Advancing Translational Sciences.

## Author contributions

CM: Conceived, planned and carried out the experiments, performed data analysis, and participated in manuscript writing. B.N.V: developed the design, theory and performed the computation analysis, also participated in manuscript writing. G.P.N, D.P: Conceived and planned the *in vivo* experiments. F.S: Performed and analyzed all chromosomal-related experiments. S.F: Performed Computational analysis. M.M, O.B.A, D.S: Contributed to sample preparation and data analysis; I.S.L: contributed to the design of some experiments, interpretation of the results, and writing the manuscript. J,K, S.L, J.S: Contributed to sample acquisition, data interpretation, and writing the manuscript. B.E: Contributed to the design of some experiments, sample acquisition, data interpretation and writing the manuscript. L.B-M: Conceived the original idea of the manuscript, planned the experiments, contributed on the sample acquisition and approach for data analysis, and interpretation of the results, and participated in manuscript writing. All authors discussed the results and contributed to the final manuscript.

## Supplemental Figures

**Supplemental Figure 1.** A. Agarose gel containing ctDNAs from Multiple myeloma, pancreatic cancer, and colon cancer ctDNA without or with treatment with Rnase A, DNase I, and proteinase K. B. Pie chart showing the single nucleotide variants (SNV) shared between tumor from patients and their corresponding ctDNA (orange) and SNV present only in the tumor sample (blue). C. GFP expression in MM1s cells cocultured with linearized Cytomegalovirus-green fluorescent protein: CMV-GFP, ctDNA bound to CMV-GFP, and ctDNA-CMV-GFP. D. ctDNA localization in tumor xenograft and multiple organs in all ctDNA tail-injected mice. Confocal microscopy of the xenograft PC tumors harvested from mice after tail injection with rhodamine-PC ctDNA. Tumors were harvested at 24 and 48 hours post-injection. E. Images of different organs harvested from xenograft-mice tail injected with rhodamine-ctDNA (MM, CC, and PC) 48 hours after tail injection (n=3 per tumor xenograft). Images were taken under the red channel to identify rhodamine fluorescence. MM: multiple myeloma, CC: colon cancer, PC: pancreatic cancer, and GFP: Green fluorescent protein.

**Supplemental Figure 2.** ctDNA integration into the cell genome. A-B. Visualization images displaying other examples of the gain of a ctDNA integration into chromatids (red) of CC (HT29 and RKO) and PC (MIA and PANC1) cell lines. Circles define zoomed regions of interest. White arrows identify an area of ctDNA integration. B. Examples of the metaphase images of various cancer cell lines (MM1s, ASPC-1, and HT116) treated with ATM, DNAPKcs, PARP, and transposase inhibitors used to generate Figure 3C. Circles define zoomed regions of interest. White arrows identify an area of ctDNA integration. C. Cell viability assays of MM1, ASPC1, and HTC116 cells after 24 hours treatment with 100 nM of ATM inhibitor (KU-55933), 30 mM of DNAPKcs Inhibitor II, 200 mM of PARP inhibitor (NU1025) or 100nM of raltegratvir. PC: Pancreatic cancer and CC: Colon cancer. MM1s: Myeloma cell line.

**Supplemental Figure 3.** A. Visualization images displaying other examples of the gain of a ctDNA SNV in the coculture experiment. B. Blast images demonstrate the transition point of a ctDNA insertion event in multiple myeloma or pancreatic cancer coculture when compared to the cell line and ctDNA contigs. Gren boxes reflect cell genome contigs, and red boxes reflect ctDNA contigs. Coculture Contigs carrying and insertion were identified using NucDiff analysis.

**Supplemental Figure 4.** Gene ontology analysis demonstrates the processes and pathways enriched in the matching coculture conditions. Gene ontology processes include biological, molecular function, and cellular functions.

**Supplemental Figure 5.** A. PCR of the Control- or all transposons performed from DNA extracted after culturing these constructs with complete media for 4 hours. B. Time course of Cy5-AluSp and -control sequence treated MM1s cells (1 μg/mL). CY5(+) cells were detected by flow cytometry. C. Dose titration experiments of Cy5-AluSp and -control sequence treated MM1s cells. Before flow cytometry, half of the samples were treated with trypsin to identify how much DNA was internalized. D. flow cytometric screening of all transposons and controls in U266 cells after 4 hours in culture. E. Graphical display of the adenine (A)-thymine (T) and guanine (G)-cytosine (C) enrich regions or both. F. Consensus sequences from all MM retrotransposons were obtained after multiple alignments using EMBOSS Cons software. G. Microscopy images of MM1s cells cultured with AluSp-CMV-mCherry, CMV-mCherry linear vector or cells transfected with CMV-mCherry circular vector. Images were captured after 24 hours of coculture with DNA.

## Supplemental Table and Video Legends

**Supplemental Table 1.** Table summarizing the distribution of the contigs containing transposons and the fraction of transposons observed at 5’ or 3’ end in inserted *vs*. non inserted ctDNA fragments.

**Supplemental Table 2.** Table summarizing the number of contigs containing a specific class and type of TE in all inserted and non-inserted contigs in MM and PC coculture experiments. Statistical analysis was performed using the Chi-square test. Significance was determined on the basis of a p-value of <0.05. MM: multiple myeloma, PC: pancreatic cancer.

**Supplemental Spreadsheet 1.** Summary of the chromosomal location of origin and insertion and their frequency in the MM (A) and PC (B) coculture experiment. CHR: chromosome. MM: multiple myeloma, PC: pancreatic cancer

**Supplemental Video 1A and B.** Rotational images of 3D reconstruction of cellular and nuclear capture of rhodamine-labeled PC ctDNA in ASPC1 cells (B) and MM ctDNA in MM1s (A). Images show cellular localization of ctDNA. Membrane identified by bright field (gray color) and ctDNA (yellow color).

**Supplemental Video 2. A.** Slide image of ASPC1 demonstrating the capturing and nuclear localization of rhodamine-labeled PC ctDNA. **B.** Different slices of Z-stack images ASPC1 cells demonstrating ctDNA capturing in the cell membrane and invagination of cell membrane for internalization of ctDNA. **C.** 3D video reconstruction of B. Cell membrane was labeled using CellLight Plasma Membrane GFP kit (TermoFisher Scientific, MA)

**Supplemental Video 3 A and B**. 3D reconstruction of colocalization of match (rhodamine – Red) and unmatched (CY5 – green) ctDNA coculture with MM1s cells.

