## Supplementary figures and images for "Retrotransposons facilitate the tissue-specific horizontal transfer of circulating tumor DNA between human cells"

### Supplemental Figure 1

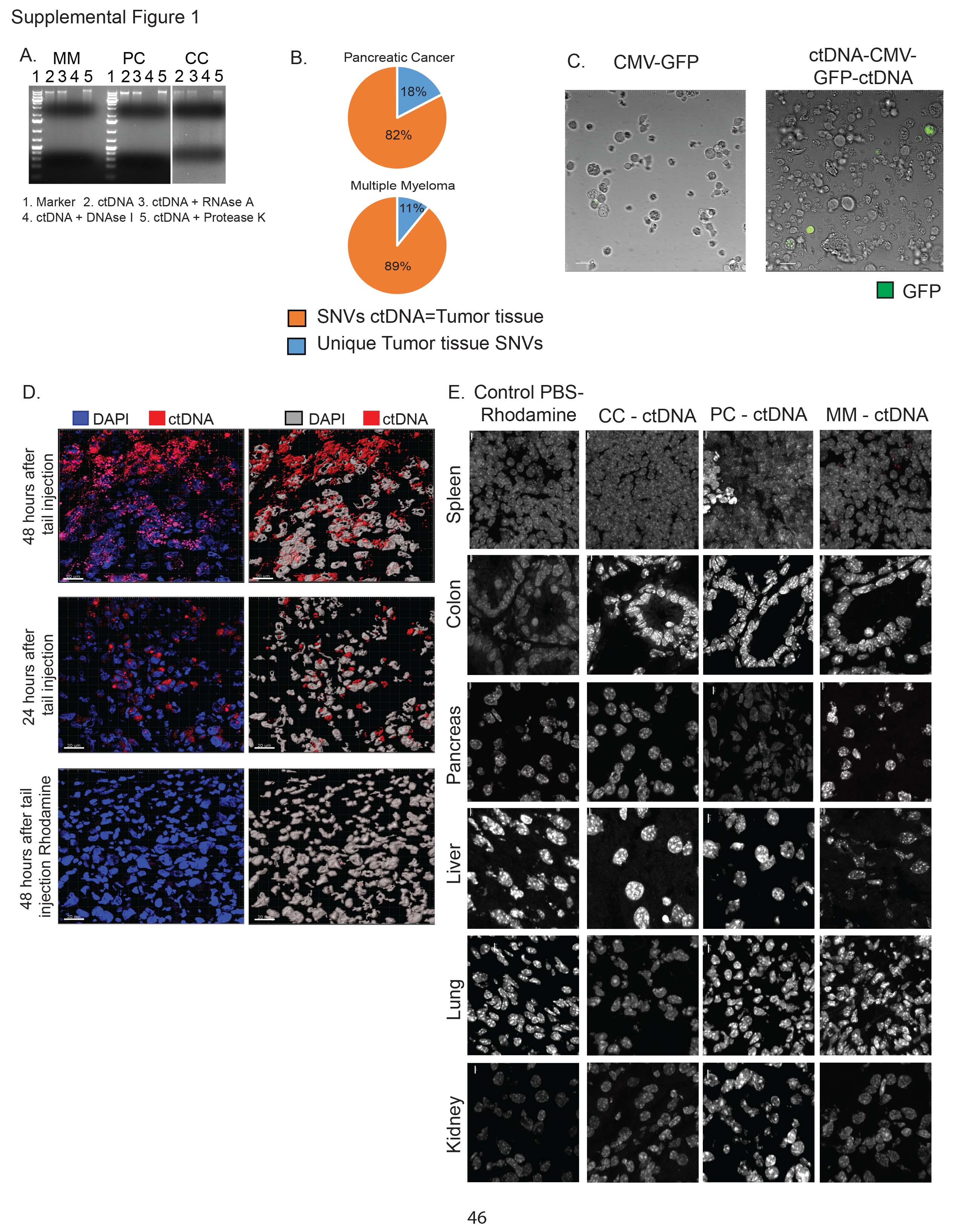

### Supplemental Figure 2

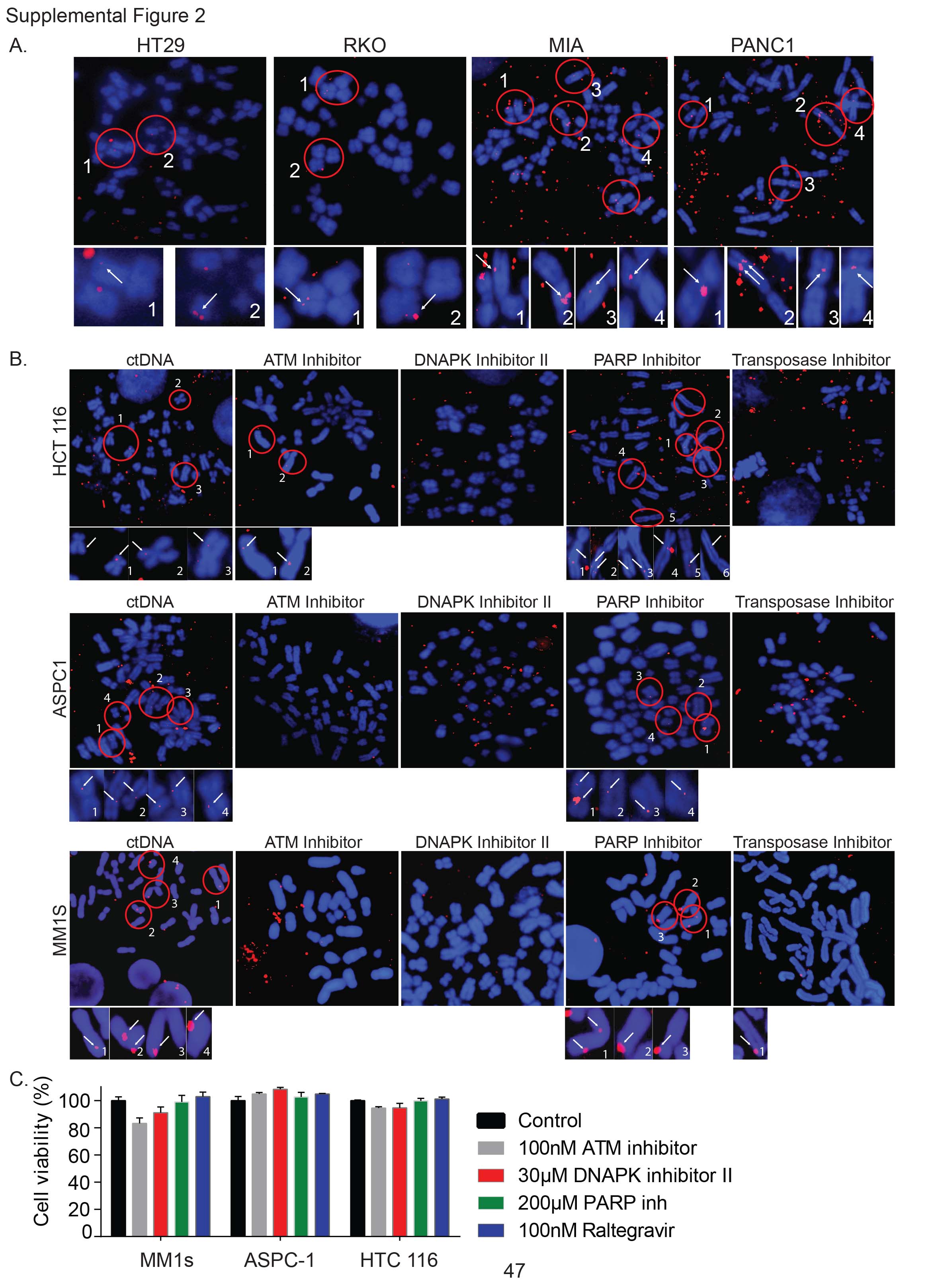

### Supplemental Figure 3

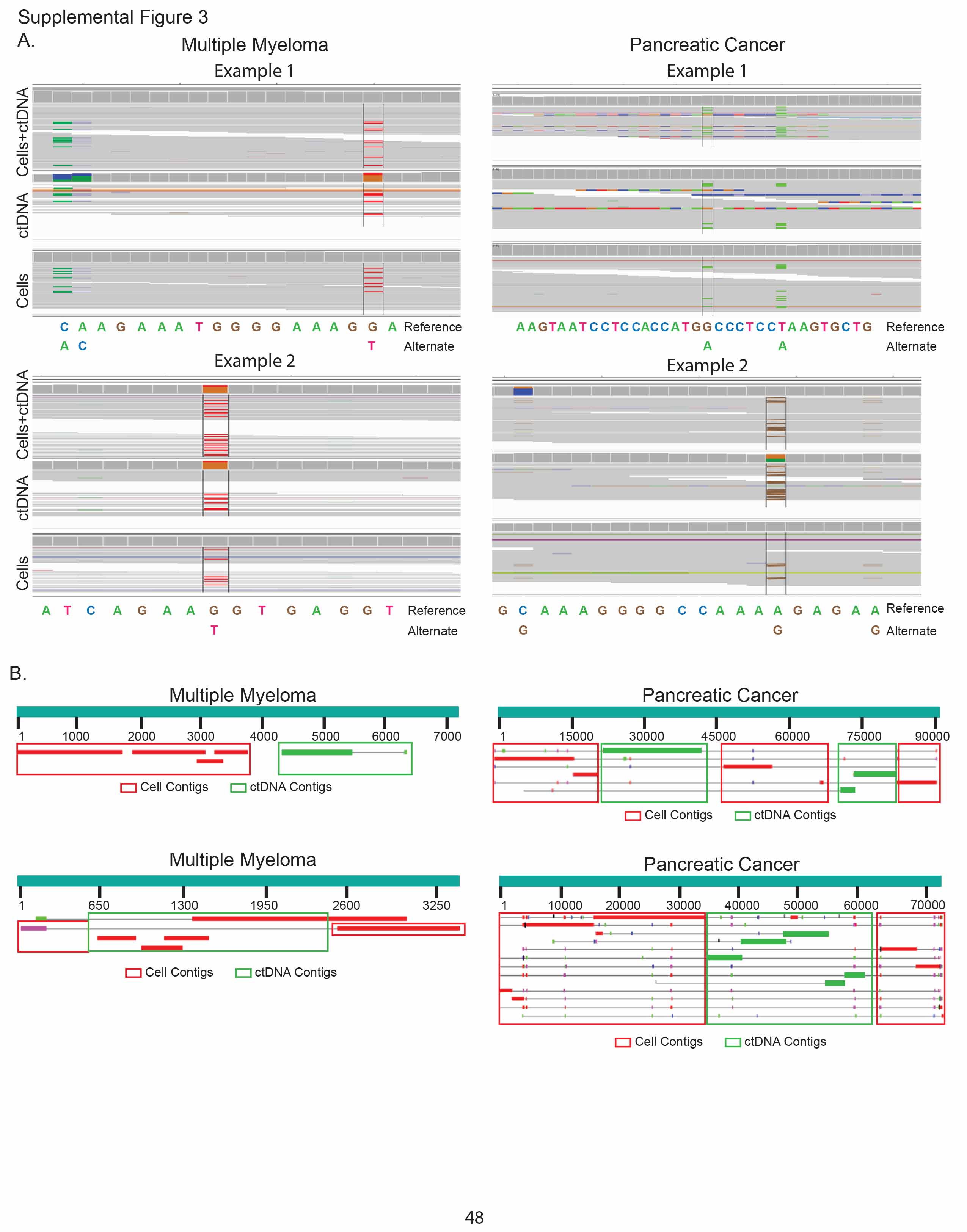

### Supplemental Figure 4

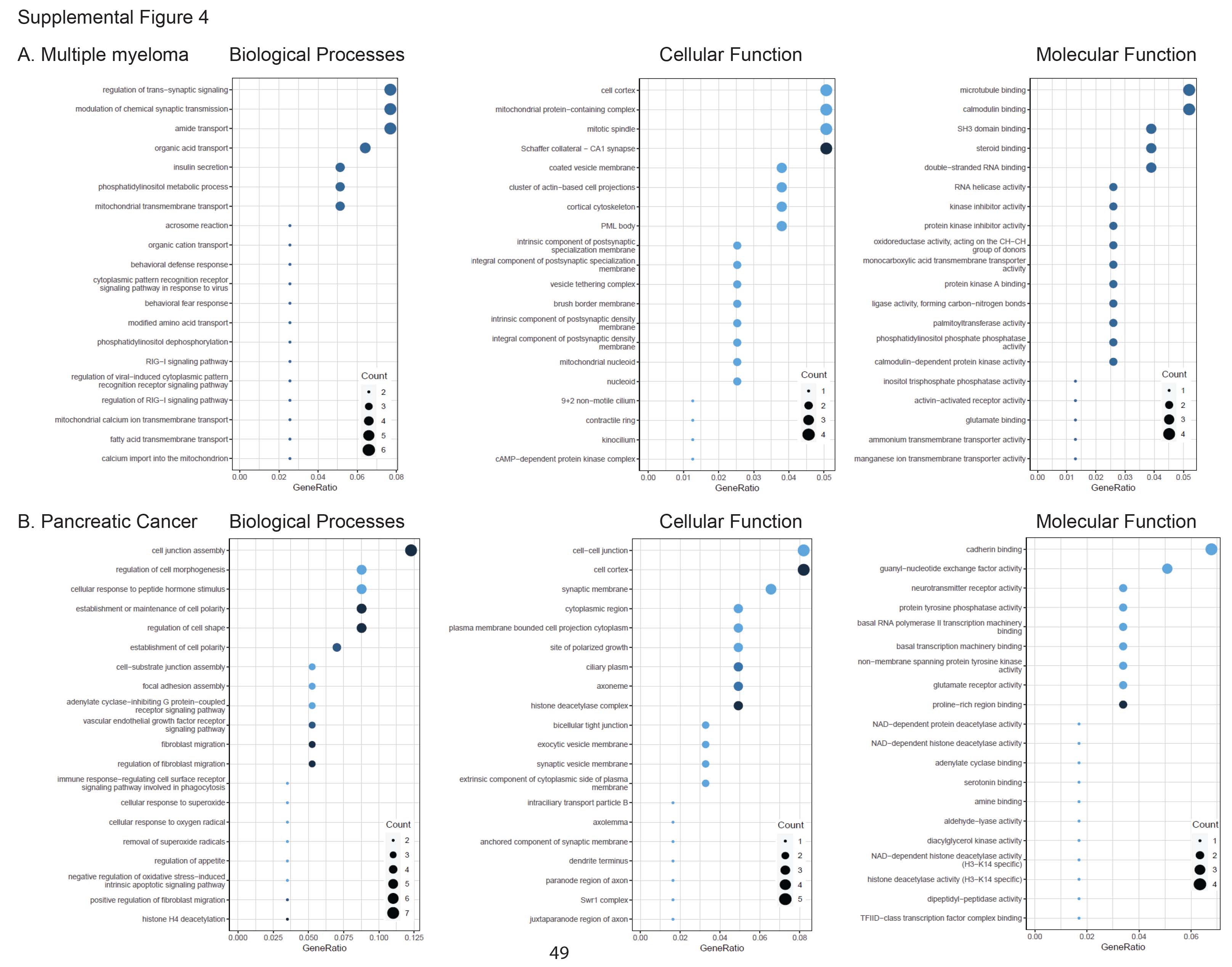

### Supplemental Figure 5

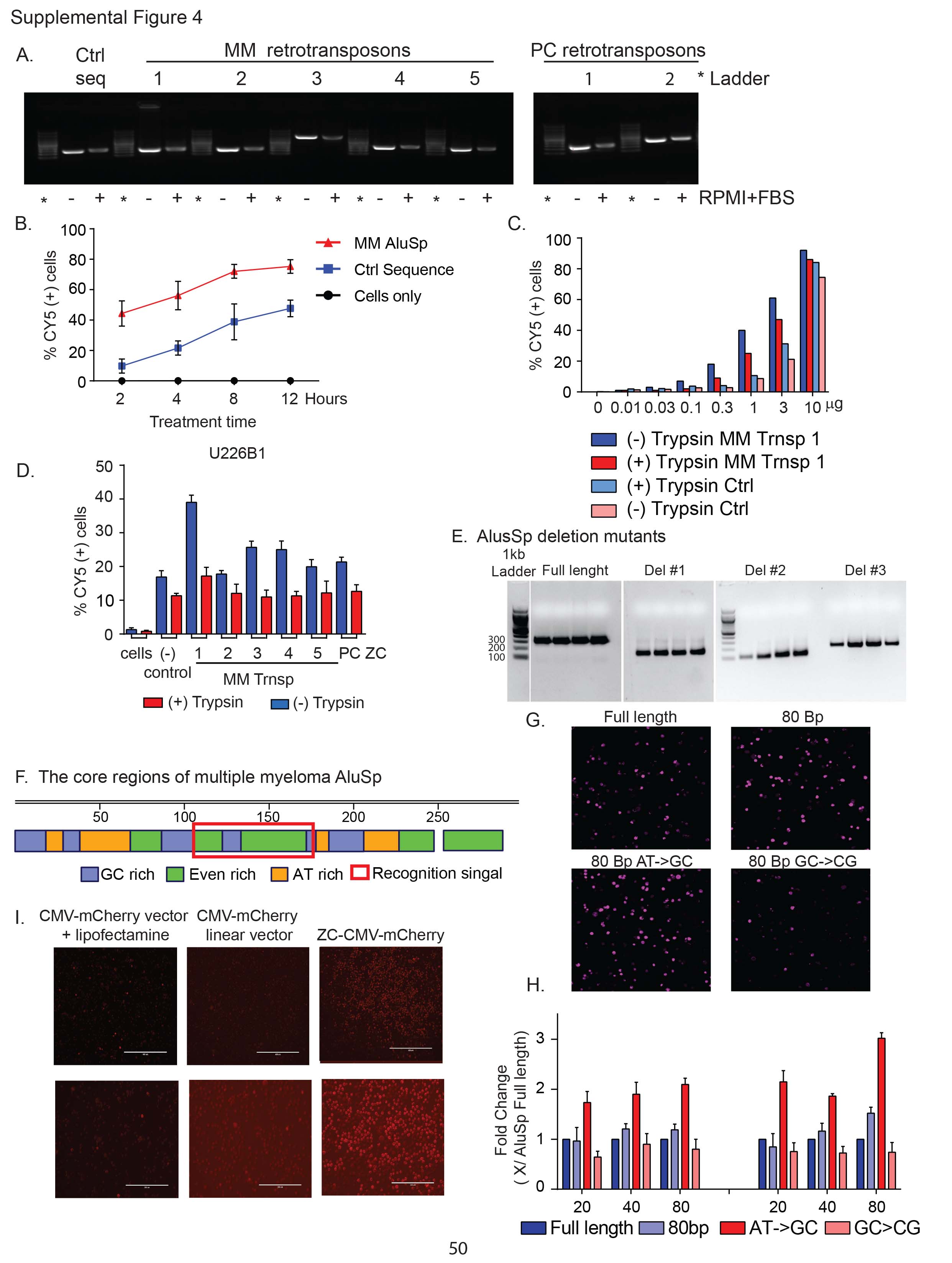

### Supplemental Video 1-2

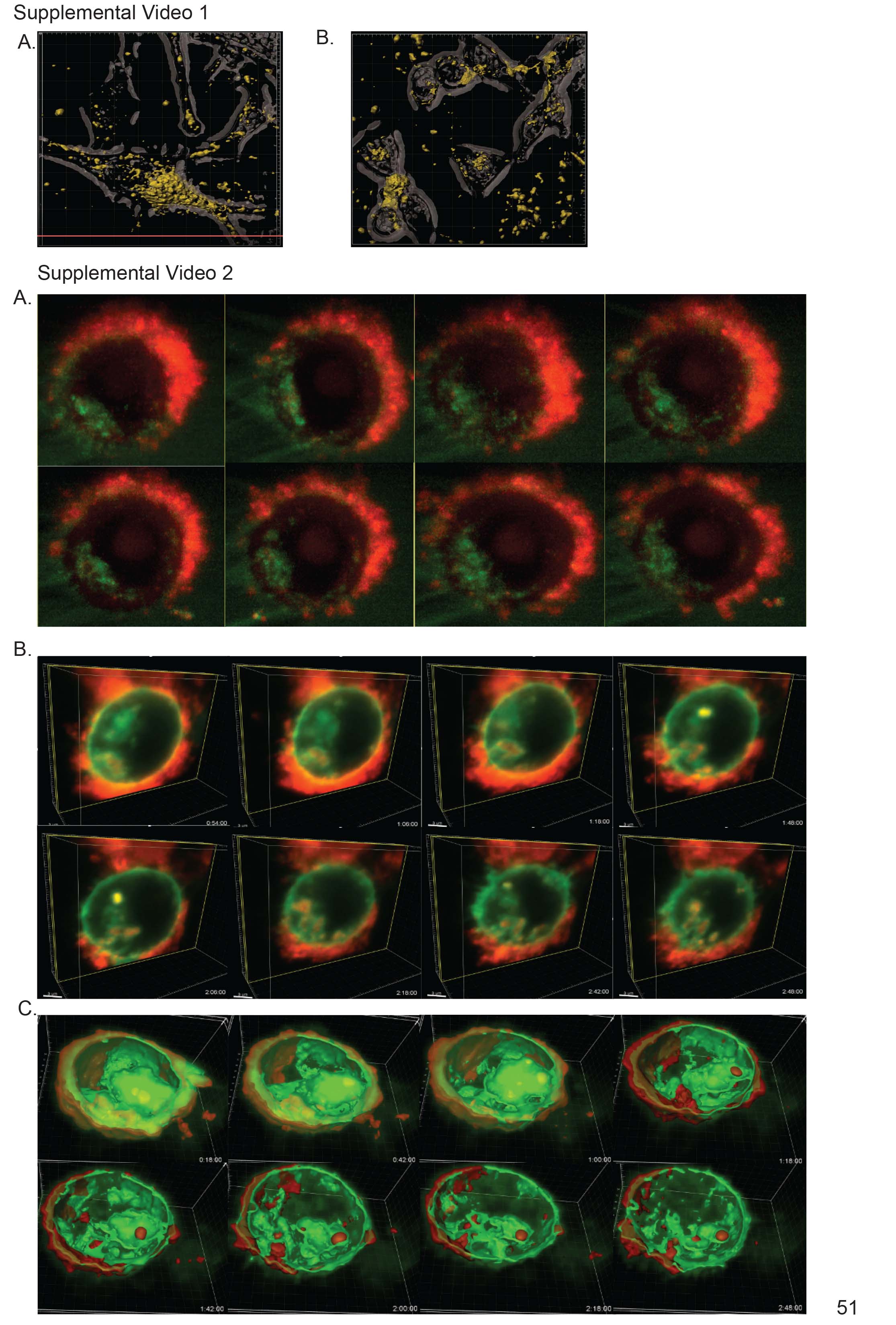

### Supplemental Video 3

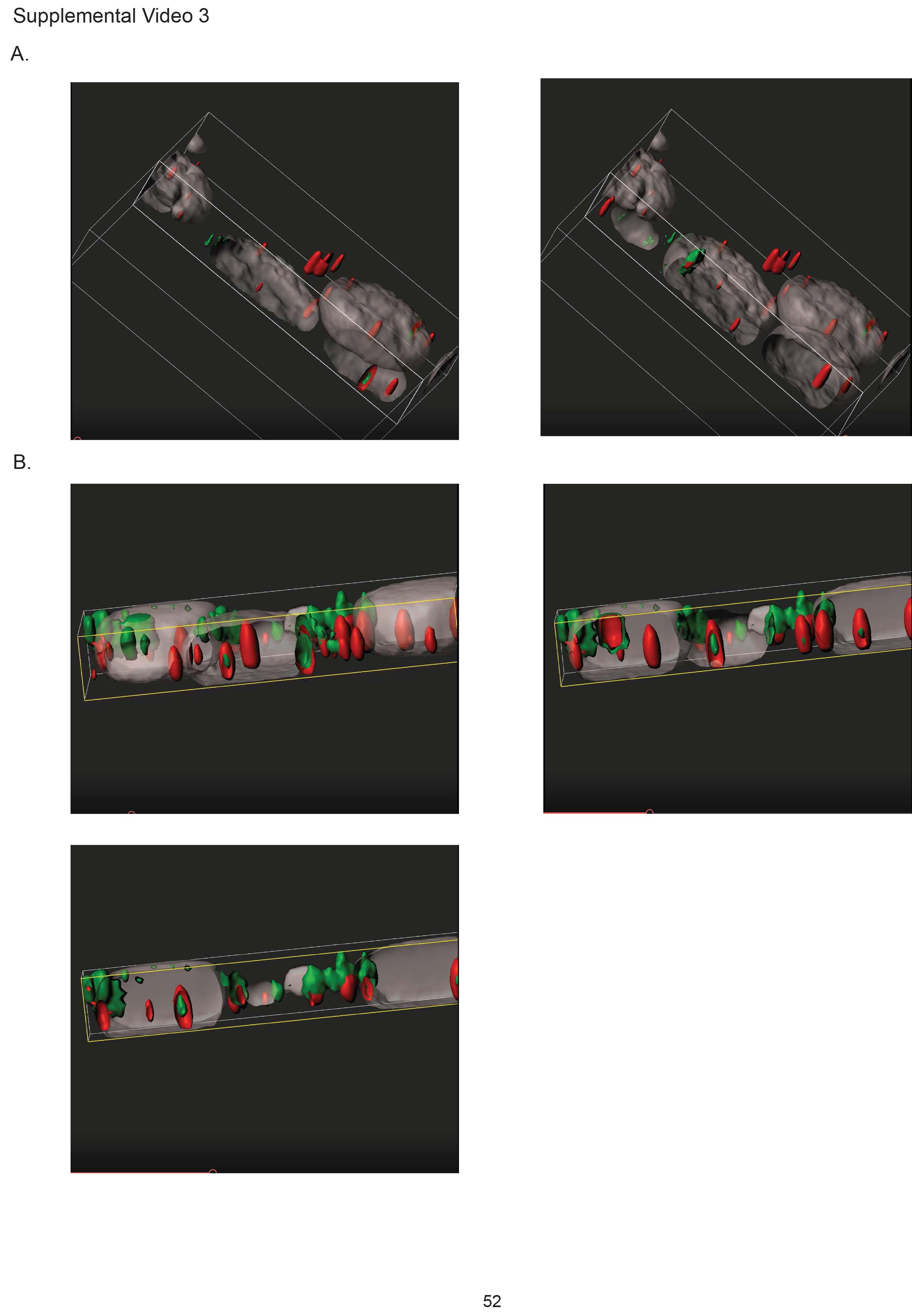
