## Supplemental Table 1A for "Retrotransposons facilitate the tissue-specific horizontal transfer of circulating tumor DNA between human cells"

**Supplemental Table 2A: Summary of the chromosomal location of origin and insertion and their frequency in the Multiple Myeloma co-culture experiment. CHR: chromosome**

| CHR.FROM | CHR.TO | No_Of_Variants | % | Same_chr |
|---|---|---|---|---|
| 3 | 3 | 1606 | 8.397825 | 1 |
| 4 | 4 | 1476 | 7.718051 | 1 |
| 2 | 2 | 1405 | 7.346789 | 1 |
| 7 | 7 | 1226 | 6.410793 | 1 |
| 1 | 1 | 1197 | 6.259151 | 1 |
| 5 | 5 | 1086 | 5.678728 | 1 |
| 6 | 6 | 902 | 4.716586 | 1 |
| 11 | 11 | 769 | 4.021125 | 1 |
| 12 | 12 | 766 | 4.005438 | 1 |
| 10 | 10 | 755 | 3.947919 | 1 |
| 8 | 8 | 636 | 3.325664 | 1 |
| 9 | 9 | 555 | 2.902113 | 1 |
| 15 | 15 | 481 | 2.515164 | 1 |
| 18 | 18 | 476 | 2.489019 | 1 |
| 17 | 17 | 421 | 2.201422 | 1 |
| X | X | 418 | 2.185735 | 1 |
| 20 | 20 | 345 | 1.804016 | 1 |
| 16 | 16 | 313 | 1.636687 | 1 |
| 13 | 13 | 272 | 1.422297 | 1 |
| 19 | 19 | 220 | 1.150387 | 1 |
| 21 | 21 | 197 | 1.030119 | 1 |
| 22 | 22 | 163 | 0.852332 | 1 |
| 14 | 14 | 79 | 0.413093 | 1 |
| 2 | 3 | 34 | 0.177787 | 0 |
| 2 | 4 | 34 | 0.177787 | 0 |
| 4 | 3 | 31 | 0.1621 | 0 |
| 7 | 2 | 30 | 0.156871 | 0 |
| 1 | 4 | 28 | 0.146413 | 0 |
| 1 | 3 | 27 | 0.141184 | 0 |
| 3 | 7 | 27 | 0.141184 | 0 |
| 3 | 2 | 26 | 0.135955 | 0 |
| 5 | 3 | 26 | 0.135955 | 0 |
| 1 | 7 | 25 | 0.130726 | 0 |
| 2 | 7 | 25 | 0.130726 | 0 |
| 5 | 7 | 25 | 0.130726 | 0 |
| 7 | 4 | 25 | 0.130726 | 0 |
| 12 | 4 | 25 | 0.130726 | 0 |
| 2 | 1 | 24 | 0.125497 | 0 |
| 5 | 4 | 24 | 0.125497 | 0 |
| 3 | 5 | 23 | 0.120268 | 0 |
| 4 | 5 | 23 | 0.120268 | 0 |
| 3 | 4 | 22 | 0.115039 | 0 |
| 4 | 7 | 22 | 0.115039 | 0 |
| 5 | 1 | 22 | 0.115039 | 0 |

| | | | | |
|---|---|---|---|---|
| 6 | 5 | 21 | 0.10981 | 0 |
| 7 | 3 | 21 | 0.10981 | 0 |
| 11 | 4 | 21 | 0.10981 | 0 |
| X | 1 | 21 | 0.10981 | 0 |
| 1 | 10 | 20 | 0.104581 | 0 |
| 2 | 6 | 20 | 0.104581 | 0 |
| 10 | 4 | 20 | 0.104581 | 0 |
| 17 | 7 | 20 | 0.104581 | 0 |
| 4 | 1 | 19 | 0.099352 | 0 |
| 8 | 4 | 19 | 0.099352 | 0 |
| 2 | 10 | 18 | 0.094123 | 0 |
| 3 | 8 | 18 | 0.094123 | 0 |
| 8 | 1 | 18 | 0.094123 | 0 |
| 9 | 3 | 18 | 0.094123 | 0 |
| 12 | 5 | 18 | 0.094123 | 0 |
| 15 | 2 | 18 | 0.094123 | 0 |
| 1 | 2 | 17 | 0.088894 | 0 |
| 1 | 18 | 17 | 0.088894 | 0 |
| 4 | 2 | 17 | 0.088894 | 0 |
| 4 | 6 | 17 | 0.088894 | 0 |
| 6 | 1 | 17 | 0.088894 | 0 |
| 11 | 7 | 17 | 0.088894 | 0 |
| 12 | 3 | 17 | 0.088894 | 0 |
| 17 | 3 | 17 | 0.088894 | 0 |
| 1 | 5 | 16 | 0.083665 | 0 |
| 2 | 5 | 16 | 0.083665 | 0 |
| 2 | 8 | 16 | 0.083665 | 0 |
| 3 | 6 | 16 | 0.083665 | 0 |
| 8 | 2 | 16 | 0.083665 | 0 |
| 8 | 3 | 16 | 0.083665 | 0 |
| 12 | 7 | 16 | 0.083665 | 0 |
| 15 | 3 | 16 | 0.083665 | 0 |
| 1 | 6 | 15 | 0.078435 | 0 |
| 3 | 11 | 15 | 0.078435 | 0 |
| 4 | 10 | 15 | 0.078435 | 0 |
| 5 | 2 | 15 | 0.078435 | 0 |
| 6 | 3 | 15 | 0.078435 | 0 |
| 6 | 4 | 15 | 0.078435 | 0 |
| 7 | 5 | 15 | 0.078435 | 0 |
| 9 | 4 | 15 | 0.078435 | 0 |
| 9 | 7 | 15 | 0.078435 | 0 |
| 10 | 2 | 15 | 0.078435 | 0 |
| 10 | 7 | 15 | 0.078435 | 0 |
| X | 7 | 15 | 0.078435 | 0 |
| 1 | 12 | 14 | 0.073206 | 0 |
| 2 | 11 | 14 | 0.073206 | 0 |
| 3 | 1 | 14 | 0.073206 | 0 |

| | | | | |
|---|---|---|---|---|
| 3 | 10 | 14 | 0.073206 | 0 |
| 3 | 12 | 14 | 0.073206 | 0 |
| 5 | 6 | 14 | 0.073206 | 0 |
| 5 | 8 | 14 | 0.073206 | 0 |
| 8 | 5 | 14 | 0.073206 | 0 |
| 11 | 1 | 14 | 0.073206 | 0 |
| 11 | 3 | 14 | 0.073206 | 0 |
| 11 | 6 | 14 | 0.073206 | 0 |
| 18 | 7 | 14 | 0.073206 | 0 |
| 18 | 12 | 14 | 0.073206 | 0 |
| 1 | 8 | 13 | 0.067977 | 0 |
| 3 | X | 13 | 0.067977 | 0 |
| 4 | 8 | 13 | 0.067977 | 0 |
| 6 | 7 | 13 | 0.067977 | 0 |
| 7 | 1 | 13 | 0.067977 | 0 |
| 8 | 7 | 13 | 0.067977 | 0 |
| 9 | 1 | 13 | 0.067977 | 0 |
| 10 | 3 | 13 | 0.067977 | 0 |
| 12 | 1 | 13 | 0.067977 | 0 |
| 15 | 4 | 13 | 0.067977 | 0 |
| 5 | 10 | 12 | 0.062748 | 0 |
| 10 | 5 | 12 | 0.062748 | 0 |
| 12 | 6 | 12 | 0.062748 | 0 |
| 12 | 17 | 12 | 0.062748 | 0 |
| 14 | 7 | 12 | 0.062748 | 0 |
| 15 | 7 | 12 | 0.062748 | 0 |
| X | 12 | 12 | 0.062748 | 0 |
| 2 | 18 | 11 | 0.057519 | 0 |
| 3 | 9 | 11 | 0.057519 | 0 |
| 4 | 12 | 11 | 0.057519 | 0 |
| 5 | X | 11 | 0.057519 | 0 |
| 6 | 15 | 11 | 0.057519 | 0 |
| 15 | 1 | 11 | 0.057519 | 0 |
| 17 | 4 | 11 | 0.057519 | 0 |
| 18 | 1 | 11 | 0.057519 | 0 |
| 18 | 10 | 11 | 0.057519 | 0 |
| 20 | 5 | 11 | 0.057519 | 0 |
| X | 3 | 11 | 0.057519 | 0 |
| 1 | 11 | 10 | 0.05229 | 0 |
| 1 | 15 | 10 | 0.05229 | 0 |
| 2 | 12 | 10 | 0.05229 | 0 |
| 2 | 20 | 10 | 0.05229 | 0 |
| 4 | 15 | 10 | 0.05229 | 0 |
| 5 | 18 | 10 | 0.05229 | 0 |
| 7 | 10 | 10 | 0.05229 | 0 |
| 10 | 1 | 10 | 0.05229 | 0 |
| 11 | 2 | 10 | 0.05229 | 0 |

| | | | | |
|---|---|---|---|---|
| 12 | 2 | 10 | 0.05229 | 0 |
| 13 | 7 | 10 | 0.05229 | 0 |
| 16 | 7 | 10 | 0.05229 | 0 |
| 18 | 4 | 10 | 0.05229 | 0 |
| 19 | 1 | 10 | 0.05229 | 0 |
| 19 | 7 | 10 | 0.05229 | 0 |
| 20 | 10 | 10 | 0.05229 | 0 |
| 21 | 4 | 10 | 0.05229 | 0 |
| X | 2 | 10 | 0.05229 | 0 |
| X | 20 | 10 | 0.05229 | 0 |
| 1 | 17 | 9 | 0.047061 | 0 |
| 1 | X | 9 | 0.047061 | 0 |
| 4 | 9 | 9 | 0.047061 | 0 |
| 4 | X | 9 | 0.047061 | 0 |
| 5 | 11 | 9 | 0.047061 | 0 |
| 5 | 13 | 9 | 0.047061 | 0 |
| 6 | 2 | 9 | 0.047061 | 0 |
| 7 | 6 | 9 | 0.047061 | 0 |
| 7 | 11 | 9 | 0.047061 | 0 |
| 8 | 6 | 9 | 0.047061 | 0 |
| 8 | 12 | 9 | 0.047061 | 0 |
| 9 | 2 | 9 | 0.047061 | 0 |
| 12 | 10 | 9 | 0.047061 | 0 |
| 12 | 11 | 9 | 0.047061 | 0 |
| 12 | 13 | 9 | 0.047061 | 0 |
| 13 | 3 | 9 | 0.047061 | 0 |
| 13 | 4 | 9 | 0.047061 | 0 |
| 17 | 12 | 9 | 0.047061 | 0 |
| 20 | 1 | 9 | 0.047061 | 0 |
| 20 | 2 | 9 | 0.047061 | 0 |
| 20 | 7 | 9 | 0.047061 | 0 |
| 2 | 9 | 8 | 0.041832 | 0 |
| 3 | 18 | 8 | 0.041832 | 0 |
| 5 | 9 | 8 | 0.041832 | 0 |
| 6 | 9 | 8 | 0.041832 | 0 |
| 8 | 15 | 8 | 0.041832 | 0 |
| 9 | 6 | 8 | 0.041832 | 0 |
| 9 | 8 | 8 | 0.041832 | 0 |
| 11 | 5 | 8 | 0.041832 | 0 |
| 13 | 1 | 8 | 0.041832 | 0 |
| 13 | 15 | 8 | 0.041832 | 0 |
| 14 | 3 | 8 | 0.041832 | 0 |
| 15 | 5 | 8 | 0.041832 | 0 |
| 15 | 10 | 8 | 0.041832 | 0 |
| 16 | 1 | 8 | 0.041832 | 0 |
| 16 | 2 | 8 | 0.041832 | 0 |
| 18 | 3 | 8 | 0.041832 | 0 |

| | | | | |
|---|---|---|---|---|
| 18 | 5 | 8 | 0.041832 | 0 |
| 18 | 6 | 8 | 0.041832 | 0 |
| 20 | 3 | 8 | 0.041832 | 0 |
| 2 | 15 | 7 | 0.036603 | 0 |
| 2 | 21 | 7 | 0.036603 | 0 |
| 3 | 13 | 7 | 0.036603 | 0 |
| 4 | 17 | 7 | 0.036603 | 0 |
| 4 | 19 | 7 | 0.036603 | 0 |
| 5 | 12 | 7 | 0.036603 | 0 |
| 5 | 15 | 7 | 0.036603 | 0 |
| 5 | 16 | 7 | 0.036603 | 0 |
| 6 | 21 | 7 | 0.036603 | 0 |
| 10 | 6 | 7 | 0.036603 | 0 |
| 11 | 10 | 7 | 0.036603 | 0 |
| 11 | X | 7 | 0.036603 | 0 |
| 13 | 2 | 7 | 0.036603 | 0 |
| 14 | 5 | 7 | 0.036603 | 0 |
| 14 | 6 | 7 | 0.036603 | 0 |
| 16 | 4 | 7 | 0.036603 | 0 |
| 19 | 2 | 7 | 0.036603 | 0 |
| 19 | 4 | 7 | 0.036603 | 0 |
| X | 4 | 7 | 0.036603 | 0 |
| X | 10 | 7 | 0.036603 | 0 |
| 1 | 16 | 6 | 0.031374 | 0 |
| 2 | 13 | 6 | 0.031374 | 0 |
| 2 | 22 | 6 | 0.031374 | 0 |
| 3 | 20 | 6 | 0.031374 | 0 |
| 5 | 20 | 6 | 0.031374 | 0 |
| 6 | 10 | 6 | 0.031374 | 0 |
| 8 | 10 | 6 | 0.031374 | 0 |
| 9 | 5 | 6 | 0.031374 | 0 |
| 9 | 10 | 6 | 0.031374 | 0 |
| 9 | 12 | 6 | 0.031374 | 0 |
| 10 | 8 | 6 | 0.031374 | 0 |
| 11 | 8 | 6 | 0.031374 | 0 |
| 11 | 9 | 6 | 0.031374 | 0 |
| 11 | 18 | 6 | 0.031374 | 0 |
| 11 | 21 | 6 | 0.031374 | 0 |
| 12 | 8 | 6 | 0.031374 | 0 |
| 12 | 18 | 6 | 0.031374 | 0 |
| 13 | 9 | 6 | 0.031374 | 0 |
| 15 | 6 | 6 | 0.031374 | 0 |
| 15 | 17 | 6 | 0.031374 | 0 |
| 15 | X | 6 | 0.031374 | 0 |
| 17 | 1 | 6 | 0.031374 | 0 |
| 17 | 2 | 6 | 0.031374 | 0 |
| 17 | 8 | 6 | 0.031374 | 0 |

| | | | | |
|---|---|---|---|---|
| 18 | 2 | 6 | 0.031374 | 0 |
| 20 | 9 | 6 | 0.031374 | 0 |
| X | 5 | 6 | 0.031374 | 0 |
| X | 6 | 6 | 0.031374 | 0 |
| X | 11 | 6 | 0.031374 | 0 |
| 1 | 13 | 5 | 0.026145 | 0 |
| 1 | 20 | 5 | 0.026145 | 0 |
| 1 | 21 | 5 | 0.026145 | 0 |
| 1 | 22 | 5 | 0.026145 | 0 |
| 2 | 14 | 5 | 0.026145 | 0 |
| 2 | 17 | 5 | 0.026145 | 0 |
| 3 | 15 | 5 | 0.026145 | 0 |
| 3 | 21 | 5 | 0.026145 | 0 |
| 4 | 11 | 5 | 0.026145 | 0 |
| 4 | 16 | 5 | 0.026145 | 0 |
| 5 | 17 | 5 | 0.026145 | 0 |
| 6 | 12 | 5 | 0.026145 | 0 |
| 6 | 17 | 5 | 0.026145 | 0 |
| 7 | 9 | 5 | 0.026145 | 0 |
| 7 | 15 | 5 | 0.026145 | 0 |
| 7 | X | 5 | 0.026145 | 0 |
| 8 | 11 | 5 | 0.026145 | 0 |
| 8 | 16 | 5 | 0.026145 | 0 |
| 8 | 17 | 5 | 0.026145 | 0 |
| 9 | 16 | 5 | 0.026145 | 0 |
| 10 | 11 | 5 | 0.026145 | 0 |
| 10 | 12 | 5 | 0.026145 | 0 |
| 10 | 13 | 5 | 0.026145 | 0 |
| 10 | X | 5 | 0.026145 | 0 |
| 11 | 12 | 5 | 0.026145 | 0 |
| 11 | 13 | 5 | 0.026145 | 0 |
| 11 | 15 | 5 | 0.026145 | 0 |
| 13 | 6 | 5 | 0.026145 | 0 |
| 14 | 1 | 5 | 0.026145 | 0 |
| 16 | 15 | 5 | 0.026145 | 0 |
| 17 | 15 | 5 | 0.026145 | 0 |
| 17 | 20 | 5 | 0.026145 | 0 |
| 19 | 9 | 5 | 0.026145 | 0 |
| 20 | 4 | 5 | 0.026145 | 0 |
| 21 | 2 | 5 | 0.026145 | 0 |
| 21 | 6 | 5 | 0.026145 | 0 |
| 22 | 3 | 5 | 0.026145 | 0 |
| X | 16 | 5 | 0.026145 | 0 |
| 4 | 13 | 4 | 0.020916 | 0 |
| 4 | 21 | 4 | 0.020916 | 0 |
| 6 | 8 | 4 | 0.020916 | 0 |
| 6 | 13 | 4 | 0.020916 | 0 |

| | | | | |
|---|---|---|---|---|
| 6 | 16 | 4 | 0.020916 | 0 |
| 6 | 18 | 4 | 0.020916 | 0 |
| 6 | 20 | 4 | 0.020916 | 0 |
| 7 | 8 | 4 | 0.020916 | 0 |
| 7 | 19 | 4 | 0.020916 | 0 |
| 8 | 9 | 4 | 0.020916 | 0 |
| 8 | 14 | 4 | 0.020916 | 0 |
| 8 | X | 4 | 0.020916 | 0 |
| 9 | 11 | 4 | 0.020916 | 0 |
| 9 | X | 4 | 0.020916 | 0 |
| 10 | 16 | 4 | 0.020916 | 0 |
| 10 | 20 | 4 | 0.020916 | 0 |
| 11 | 22 | 4 | 0.020916 | 0 |
| 12 | X | 4 | 0.020916 | 0 |
| 13 | 8 | 4 | 0.020916 | 0 |
| 13 | 12 | 4 | 0.020916 | 0 |
| 13 | 18 | 4 | 0.020916 | 0 |
| 13 | 21 | 4 | 0.020916 | 0 |
| 13 | X | 4 | 0.020916 | 0 |
| 15 | 8 | 4 | 0.020916 | 0 |
| 15 | 16 | 4 | 0.020916 | 0 |
| 15 | 18 | 4 | 0.020916 | 0 |
| 15 | 19 | 4 | 0.020916 | 0 |
| 16 | 5 | 4 | 0.020916 | 0 |
| 16 | 6 | 4 | 0.020916 | 0 |
| 16 | 11 | 4 | 0.020916 | 0 |
| 16 | 12 | 4 | 0.020916 | 0 |
| 16 | 18 | 4 | 0.020916 | 0 |
| 17 | 5 | 4 | 0.020916 | 0 |
| 17 | 6 | 4 | 0.020916 | 0 |
| 17 | 11 | 4 | 0.020916 | 0 |
| 17 | 18 | 4 | 0.020916 | 0 |
| 18 | 8 | 4 | 0.020916 | 0 |
| 18 | 17 | 4 | 0.020916 | 0 |
| 21 | 3 | 4 | 0.020916 | 0 |
| 21 | 7 | 4 | 0.020916 | 0 |
| 21 | 10 | 4 | 0.020916 | 0 |
| 21 | 11 | 4 | 0.020916 | 0 |
| 22 | 2 | 4 | 0.020916 | 0 |
| 22 | 11 | 4 | 0.020916 | 0 |
| X | 8 | 4 | 0.020916 | 0 |
| X | 13 | 4 | 0.020916 | 0 |
| X | 15 | 4 | 0.020916 | 0 |
| X | 18 | 4 | 0.020916 | 0 |
| 1 | 9 | 3 | 0.015687 | 0 |
| 2 | X | 3 | 0.015687 | 0 |
| 3 | 17 | 3 | 0.015687 | 0 |

| | | | | |
|---|---|---|---|---|
| 3 | 22 | 3 | 0.015687 | 0 |
| 4 | 14 | 3 | 0.015687 | 0 |
| 4 | 18 | 3 | 0.015687 | 0 |
| 4 | 20 | 3 | 0.015687 | 0 |
| 5 | 21 | 3 | 0.015687 | 0 |
| 6 | 11 | 3 | 0.015687 | 0 |
| 7 | 17 | 3 | 0.015687 | 0 |
| 7 | 18 | 3 | 0.015687 | 0 |
| 7 | 21 | 3 | 0.015687 | 0 |
| 8 | 18 | 3 | 0.015687 | 0 |
| 8 | 20 | 3 | 0.015687 | 0 |
| 8 | 21 | 3 | 0.015687 | 0 |
| 9 | 13 | 3 | 0.015687 | 0 |
| 9 | 15 | 3 | 0.015687 | 0 |
| 9 | 18 | 3 | 0.015687 | 0 |
| 10 | 9 | 3 | 0.015687 | 0 |
| 10 | 15 | 3 | 0.015687 | 0 |
| 11 | 19 | 3 | 0.015687 | 0 |
| 12 | 20 | 3 | 0.015687 | 0 |
| 13 | 10 | 3 | 0.015687 | 0 |
| 13 | 11 | 3 | 0.015687 | 0 |
| 14 | 2 | 3 | 0.015687 | 0 |
| 14 | 4 | 3 | 0.015687 | 0 |
| 14 | 8 | 3 | 0.015687 | 0 |
| 14 | 9 | 3 | 0.015687 | 0 |
| 14 | 12 | 3 | 0.015687 | 0 |
| 14 | 15 | 3 | 0.015687 | 0 |
| 15 | 11 | 3 | 0.015687 | 0 |
| 15 | 20 | 3 | 0.015687 | 0 |
| 16 | 3 | 3 | 0.015687 | 0 |
| 16 | 17 | 3 | 0.015687 | 0 |
| 16 | 19 | 3 | 0.015687 | 0 |
| 16 | X | 3 | 0.015687 | 0 |
| 17 | 10 | 3 | 0.015687 | 0 |
| 18 | 11 | 3 | 0.015687 | 0 |
| 18 | 15 | 3 | 0.015687 | 0 |
| 18 | 20 | 3 | 0.015687 | 0 |
| 18 | 21 | 3 | 0.015687 | 0 |
| 18 | X | 3 | 0.015687 | 0 |
| 19 | 3 | 3 | 0.015687 | 0 |
| 19 | 5 | 3 | 0.015687 | 0 |
| 19 | 11 | 3 | 0.015687 | 0 |
| 19 | 16 | 3 | 0.015687 | 0 |
| 19 | 17 | 3 | 0.015687 | 0 |
| 20 | 6 | 3 | 0.015687 | 0 |
| 20 | 8 | 3 | 0.015687 | 0 |
| 20 | 11 | 3 | 0.015687 | 0 |

| | | | | |
|---|---|---|---|---|
| 20 | 17 | 3 | 0.015687 | 0 |
| 21 | 5 | 3 | 0.015687 | 0 |
| 22 | 1 | 3 | 0.015687 | 0 |
| 22 | 5 | 3 | 0.015687 | 0 |
| X | 9 | 3 | 0.015687 | 0 |
| 1 | 14 | 2 | 0.010458 | 0 |
| 2 | 19 | 2 | 0.010458 | 0 |
| 3 | 16 | 2 | 0.010458 | 0 |
| 3 | 19 | 2 | 0.010458 | 0 |
| 4 | 22 | 2 | 0.010458 | 0 |
| 5 | 14 | 2 | 0.010458 | 0 |
| 5 | 19 | 2 | 0.010458 | 0 |
| 6 | 14 | 2 | 0.010458 | 0 |
| 6 | X | 2 | 0.010458 | 0 |
| 7 | 20 | 2 | 0.010458 | 0 |
| 7 | 22 | 2 | 0.010458 | 0 |
| 8 | 13 | 2 | 0.010458 | 0 |
| 8 | 22 | 2 | 0.010458 | 0 |
| 9 | 20 | 2 | 0.010458 | 0 |
| 9 | 21 | 2 | 0.010458 | 0 |
| 10 | 14 | 2 | 0.010458 | 0 |
| 10 | 17 | 2 | 0.010458 | 0 |
| 10 | 19 | 2 | 0.010458 | 0 |
| 11 | 16 | 2 | 0.010458 | 0 |
| 11 | 17 | 2 | 0.010458 | 0 |
| 11 | 20 | 2 | 0.010458 | 0 |
| 12 | 15 | 2 | 0.010458 | 0 |
| 13 | 5 | 2 | 0.010458 | 0 |
| 13 | 20 | 2 | 0.010458 | 0 |
| 14 | 10 | 2 | 0.010458 | 0 |
| 14 | 13 | 2 | 0.010458 | 0 |
| 14 | 18 | 2 | 0.010458 | 0 |
| 14 | X | 2 | 0.010458 | 0 |
| 15 | 9 | 2 | 0.010458 | 0 |
| 15 | 12 | 2 | 0.010458 | 0 |
| 15 | 13 | 2 | 0.010458 | 0 |
| 15 | 21 | 2 | 0.010458 | 0 |
| 16 | 10 | 2 | 0.010458 | 0 |
| 17 | 9 | 2 | 0.010458 | 0 |
| 17 | 19 | 2 | 0.010458 | 0 |
| 17 | 21 | 2 | 0.010458 | 0 |
| 19 | 12 | 2 | 0.010458 | 0 |
| 19 | 22 | 2 | 0.010458 | 0 |
| 20 | 18 | 2 | 0.010458 | 0 |
| 21 | 1 | 2 | 0.010458 | 0 |
| 21 | 12 | 2 | 0.010458 | 0 |
| 22 | 4 | 2 | 0.010458 | 0 |

| | | | | |
|---|---|---|---|---|
| 22 | 6 | 2 | 0.010458 | 0 |
| 22 | 10 | 2 | 0.010458 | 0 |
| 22 | 15 | 2 | 0.010458 | 0 |
| 22 | 18 | 2 | 0.010458 | 0 |
| X | 19 | 2 | 0.010458 | 0 |
| 1 | 19 | 1 | 0.005229 | 0 |
| 5 | 22 | 1 | 0.005229 | 0 |
| 6 | 19 | 1 | 0.005229 | 0 |
| 6 | 22 | 1 | 0.005229 | 0 |
| 7 | 12 | 1 | 0.005229 | 0 |
| 7 | 13 | 1 | 0.005229 | 0 |
| 9 | 14 | 1 | 0.005229 | 0 |
| 9 | 17 | 1 | 0.005229 | 0 |
| 9 | 19 | 1 | 0.005229 | 0 |
| 10 | 18 | 1 | 0.005229 | 0 |
| 11 | 14 | 1 | 0.005229 | 0 |
| 12 | 16 | 1 | 0.005229 | 0 |
| 12 | 21 | 1 | 0.005229 | 0 |
| 12 | 22 | 1 | 0.005229 | 0 |
| 13 | 16 | 1 | 0.005229 | 0 |
| 13 | 17 | 1 | 0.005229 | 0 |
| 13 | 19 | 1 | 0.005229 | 0 |
| 13 | 22 | 1 | 0.005229 | 0 |
| 14 | 11 | 1 | 0.005229 | 0 |
| 14 | 20 | 1 | 0.005229 | 0 |
| 14 | 21 | 1 | 0.005229 | 0 |
| 15 | 14 | 1 | 0.005229 | 0 |
| 15 | 22 | 1 | 0.005229 | 0 |
| 16 | 9 | 1 | 0.005229 | 0 |
| 16 | 20 | 1 | 0.005229 | 0 |
| 17 | 13 | 1 | 0.005229 | 0 |
| 17 | 16 | 1 | 0.005229 | 0 |
| 18 | 9 | 1 | 0.005229 | 0 |
| 18 | 16 | 1 | 0.005229 | 0 |
| 19 | 6 | 1 | 0.005229 | 0 |
| 19 | 8 | 1 | 0.005229 | 0 |
| 19 | 10 | 1 | 0.005229 | 0 |
| 19 | 15 | 1 | 0.005229 | 0 |
| 20 | 12 | 1 | 0.005229 | 0 |
| 20 | 13 | 1 | 0.005229 | 0 |
| 20 | 15 | 1 | 0.005229 | 0 |
| 20 | 19 | 1 | 0.005229 | 0 |
| 20 | 22 | 1 | 0.005229 | 0 |
| 21 | 8 | 1 | 0.005229 | 0 |
| 21 | 9 | 1 | 0.005229 | 0 |
| 21 | 15 | 1 | 0.005229 | 0 |
| 21 | 17 | 1 | 0.005229 | 0 |

| | | | | |
|---|---|---|---|---|
| 21 | 18 | 1 | 0.005229 | 0 |
| 21 | 19 | 1 | 0.005229 | 0 |
| 22 | 7 | 1 | 0.005229 | 0 |
| 22 | 8 | 1 | 0.005229 | 0 |
| 22 | 12 | 1 | 0.005229 | 0 |
| 22 | 13 | 1 | 0.005229 | 0 |
| 22 | 16 | 1 | 0.005229 | 0 |
| 22 | 20 | 1 | 0.005229 | 0 |
| 22 | X | 1 | 0.005229 | 0 |
| X | 14 | 1 | 0.005229 | 0 |
| X | 17 | 1 | 0.005229 | 0 |
| X | 21 | 1 | 0.005229 | 0 |
| X | 22 | 1 | 0.005229 | 0 |
| 1 | Y | 0 | 0 | 0 |
| 2 | 16 | 0 | 0 | 0 |
| 2 | Y | 0 | 0 | 0 |
| 3 | 14 | 0 | 0 | 0 |
| 3 | Y | 0 | 0 | 0 |
| 4 | Y | 0 | 0 | 0 |
| 5 | Y | 0 | 0 | 0 |
| 6 | Y | 0 | 0 | 0 |
| 7 | 14 | 0 | 0 | 0 |
| 7 | 16 | 0 | 0 | 0 |
| 7 | Y | 0 | 0 | 0 |
| 8 | 19 | 0 | 0 | 0 |
| 8 | Y | 0 | 0 | 0 |
| 9 | 22 | 0 | 0 | 0 |
| 9 | Y | 0 | 0 | 0 |
| 10 | 21 | 0 | 0 | 0 |
| 10 | 22 | 0 | 0 | 0 |
| 10 | Y | 0 | 0 | 0 |
| 11 | Y | 0 | 0 | 0 |
| 12 | 9 | 0 | 0 | 0 |
| 12 | 14 | 0 | 0 | 0 |
| 12 | 19 | 0 | 0 | 0 |
| 12 | Y | 0 | 0 | 0 |
| 13 | 14 | 0 | 0 | 0 |
| 13 | Y | 0 | 0 | 0 |
| 14 | 16 | 0 | 0 | 0 |
| 14 | 17 | 0 | 0 | 0 |
| 14 | 19 | 0 | 0 | 0 |
| 14 | 22 | 0 | 0 | 0 |
| 14 | Y | 0 | 0 | 0 |
| 15 | Y | 0 | 0 | 0 |
| 16 | 8 | 0 | 0 | 0 |
| 16 | 13 | 0 | 0 | 0 |
| 16 | 14 | 0 | 0 | 0 |

| | | | | |
|---|---|---|---|---|
| 16 | 21 | 0 | 0 | 0 |
| 16 | 22 | 0 | 0 | 0 |
| 16 | Y | 0 | 0 | 0 |
| 17 | 14 | 0 | 0 | 0 |
| 17 | 22 | 0 | 0 | 0 |
| 17 | X | 0 | 0 | 0 |
| 17 | Y | 0 | 0 | 0 |
| 18 | 13 | 0 | 0 | 0 |
| 18 | 14 | 0 | 0 | 0 |
| 18 | 19 | 0 | 0 | 0 |
| 18 | 22 | 0 | 0 | 0 |
| 18 | Y | 0 | 0 | 0 |
| 19 | 13 | 0 | 0 | 0 |
| 19 | 14 | 0 | 0 | 0 |
| 19 | 18 | 0 | 0 | 0 |
| 19 | 20 | 0 | 0 | 0 |
| 19 | 21 | 0 | 0 | 0 |
| 19 | X | 0 | 0 | 0 |
| 19 | Y | 0 | 0 | 0 |
| 20 | 14 | 0 | 0 | 0 |
| 20 | 16 | 0 | 0 | 0 |
| 20 | 21 | 0 | 0 | 0 |
| 20 | X | 0 | 0 | 0 |
| 20 | Y | 0 | 0 | 0 |
| 21 | 13 | 0 | 0 | 0 |
| 21 | 14 | 0 | 0 | 0 |
| 21 | 16 | 0 | 0 | 0 |
| 21 | 20 | 0 | 0 | 0 |
| 21 | 22 | 0 | 0 | 0 |
| 21 | X | 0 | 0 | 0 |
| 21 | Y | 0 | 0 | 0 |
| 22 | 9 | 0 | 0 | 0 |
| 22 | 14 | 0 | 0 | 0 |
| 22 | 17 | 0 | 0 | 0 |
| 22 | 19 | 0 | 0 | 0 |
| 22 | 21 | 0 | 0 | 0 |
| 22 | Y | 0 | 0 | 0 |
| X | Y | 0 | 0 | 0 |
| Y | 1 | 0 | 0 | 0 |
| Y | 2 | 0 | 0 | 0 |
| Y | 3 | 0 | 0 | 0 |
| Y | 4 | 0 | 0 | 0 |
| Y | 5 | 0 | 0 | 0 |
| Y | 6 | 0 | 0 | 0 |
| Y | 7 | 0 | 0 | 0 |
| Y | 8 | 0 | 0 | 0 |
| Y | 9 | 0 | 0 | 0 |

| | | | | |
|---|---|---|---|---|
| Y | 10 | 0 | 0 | 0 |
| Y | 11 | 0 | 0 | 0 |
| Y | 12 | 0 | 0 | 0 |
| Y | 13 | 0 | 0 | 0 |
| Y | 14 | 0 | 0 | 0 |
| Y | 15 | 0 | 0 | 0 |
| Y | 16 | 0 | 0 | 0 |
| Y | 17 | 0 | 0 | 0 |
| Y | 18 | 0 | 0 | 0 |
| Y | 19 | 0 | 0 | 0 |
| Y | 20 | 0 | 0 | 0 |
| Y | 21 | 0 | 0 | 0 |
| Y | 22 | 0 | 0 | 0 |
| Y | X | 0 | 0 | 0 |
| Y | Y | 0 | 0 | 1 |
