## Supplemental Table 1B for "Retrotransposons facilitate the tissue-specific horizontal transfer of circulating tumor DNA between human cells"

**Supplemental Table 2B: Summary of the chromosomal location of origin and insertion and their frequency in Pacreatic Cancer co-culture experiment. CHR: chromosome**

| CHR.FROM | CHR.TO | _Of_Varia | % | Same_Chr |
|---|---|---|---|---|
| 2 | 2 | 5667 | 8.147978 | 1 |
| 1 | 1 | 5260 | 7.562796 | 1 |
| 3 | 3 | 4860 | 6.987678 | 1 |
| 7 | 7 | 4120 | 5.923711 | 1 |
| 6 | 6 | 4118 | 5.920835 | 1 |
| 5 | 5 | 4025 | 5.78712 | 1 |
| 10 | 10 | 3089 | 4.441345 | 1 |
| 11 | 11 | 3085 | 4.435594 | 1 |
| 12 | 12 | 2810 | 4.040201 | 1 |
| 8 | 8 | 2377 | 3.417636 | 1 |
| 16 | 16 | 2305 | 3.314115 | 1 |
| 9 | 9 | 2086 | 2.999238 | 1 |
| 17 | 17 | 2080 | 2.990611 | 1 |
| 14 | 14 | 2071 | 2.977671 | 1 |
| 18 | 18 | 1729 | 2.485946 | 1 |
| 20 | 20 | 1724 | 2.478757 | 1 |
| 4 | 4 | 1294 | 1.860505 | 1 |
| 21 | 21 | 1162 | 1.670716 | 1 |
| 19 | 19 | 1153 | 1.657776 | 1 |
| 15 | 15 | 763 | 1.097037 | 1 |
| 13 | 13 | 674 | 0.969073 | 1 |
| 22 | 22 | 544 | 0.78216 | 1 |
| X | X | 503 | 0.72321 | 1 |
| 2 | 1 | 135 | 0.194102 | 0 |
| 2 | 3 | 124 | 0.178286 | 0 |
| 5 | 2 | 107 | 0.153844 | 0 |
| 5 | 3 | 107 | 0.153844 | 0 |
| 3 | 2 | 102 | 0.146655 | 0 |
| 1 | 2 | 99 | 0.142342 | 0 |
| 2 | 5 | 99 | 0.142342 | 0 |
| 1 | 3 | 94 | 0.135153 | 0 |
| 3 | 6 | 90 | 0.129401 | 0 |
| 1 | 5 | 85 | 0.122212 | 0 |
| 1 | 6 | 84 | 0.120775 | 0 |
| 5 | 1 | 83 | 0.119337 | 0 |
| 3 | 5 | 81 | 0.116461 | 0 |
| 2 | 7 | 79 | 0.113586 | 0 |
| 6 | 2 | 79 | 0.113586 | 0 |
| 1 | 7 | 76 | 0.109272 | 0 |
| 2 | 6 | 76 | 0.109272 | 0 |
| 3 | 7 | 73 | 0.104959 | 0 |
| 7 | 2 | 72 | 0.103521 | 0 |
| 3 | 1 | 71 | 0.102083 | 0 |
| 1 | 10 | 70 | 0.100646 | 0 |

| | | | | |
|---|---|---|---|---|
| 6 | 3 | 68 | 0.09777 | 0 |
| 10 | 2 | 68 | 0.09777 | 0 |
| 5 | 7 | 67 | 0.096332 | 0 |
| 11 | 1 | 67 | 0.096332 | 0 |
| 4 | 2 | 66 | 0.094894 | 0 |
| 6 | 1 | 66 | 0.094894 | 0 |
| 6 | 7 | 66 | 0.094894 | 0 |
| 7 | 6 | 66 | 0.094894 | 0 |
| 8 | 2 | 64 | 0.092019 | 0 |
| 11 | 2 | 63 | 0.090581 | 0 |
| 11 | 6 | 63 | 0.090581 | 0 |
| 5 | 6 | 62 | 0.089143 | 0 |
| 2 | 11 | 61 | 0.087705 | 0 |
| 7 | 3 | 61 | 0.087705 | 0 |
| 1 | 11 | 60 | 0.086268 | 0 |
| 12 | 1 | 59 | 0.08483 | 0 |
| 6 | 5 | 58 | 0.083392 | 0 |
| 4 | 3 | 56 | 0.080516 | 0 |
| 8 | 1 | 56 | 0.080516 | 0 |
| 10 | 6 | 56 | 0.080516 | 0 |
| X | 3 | 56 | 0.080516 | 0 |
| 2 | 12 | 55 | 0.079079 | 0 |
| 10 | 5 | 55 | 0.079079 | 0 |
| 12 | 3 | 55 | 0.079079 | 0 |
| 5 | 10 | 54 | 0.077641 | 0 |
| 4 | 1 | 53 | 0.076203 | 0 |
| 11 | 3 | 53 | 0.076203 | 0 |
| 12 | 2 | 53 | 0.076203 | 0 |
| 3 | 11 | 52 | 0.074765 | 0 |
| 7 | 1 | 52 | 0.074765 | 0 |
| 7 | 5 | 52 | 0.074765 | 0 |
| X | 2 | 52 | 0.074765 | 0 |
| 8 | 7 | 51 | 0.073327 | 0 |
| 1 | 14 | 50 | 0.07189 | 0 |
| 10 | 1 | 50 | 0.07189 | 0 |
| 12 | 6 | 50 | 0.07189 | 0 |
| 2 | 16 | 49 | 0.070452 | 0 |
| 4 | 6 | 49 | 0.070452 | 0 |
| 8 | 6 | 48 | 0.069014 | 0 |
| X | 6 | 48 | 0.069014 | 0 |
| 1 | 12 | 47 | 0.067576 | 0 |
| 2 | 8 | 47 | 0.067576 | 0 |
| 2 | 10 | 47 | 0.067576 | 0 |
| 11 | 5 | 47 | 0.067576 | 0 |
| 2 | 14 | 46 | 0.066139 | 0 |
| 4 | 5 | 46 | 0.066139 | 0 |
| 6 | 11 | 46 | 0.066139 | 0 |

| | | | | |
|---|---|---|---|---|
| 8 | 3 | 46 | 0.066139 | 0 |
| 10 | 7 | 46 | 0.066139 | 0 |
| 14 | 1 | 46 | 0.066139 | 0 |
| 3 | 12 | 45 | 0.064701 | 0 |
| 5 | 11 | 45 | 0.064701 | 0 |
| 9 | 1 | 45 | 0.064701 | 0 |
| 2 | 9 | 44 | 0.063263 | 0 |
| 5 | 12 | 44 | 0.063263 | 0 |
| 6 | 16 | 44 | 0.063263 | 0 |
| 7 | 8 | 44 | 0.063263 | 0 |
| 11 | 7 | 44 | 0.063263 | 0 |
| 2 | 17 | 43 | 0.061825 | 0 |
| 3 | 10 | 43 | 0.061825 | 0 |
| 14 | 6 | 43 | 0.061825 | 0 |
| X | 1 | 43 | 0.061825 | 0 |
| 9 | 3 | 42 | 0.060387 | 0 |
| 11 | 9 | 42 | 0.060387 | 0 |
| 14 | 3 | 42 | 0.060387 | 0 |
| 1 | 9 | 41 | 0.05895 | 0 |
| 4 | 7 | 41 | 0.05895 | 0 |
| 6 | 10 | 41 | 0.05895 | 0 |
| 10 | 3 | 41 | 0.05895 | 0 |
| X | 5 | 41 | 0.05895 | 0 |
| 1 | 17 | 40 | 0.057512 | 0 |
| 3 | 8 | 40 | 0.057512 | 0 |
| 6 | 12 | 40 | 0.057512 | 0 |
| 12 | 7 | 40 | 0.057512 | 0 |
| 5 | 8 | 39 | 0.056074 | 0 |
| 8 | 5 | 39 | 0.056074 | 0 |
| 9 | 2 | 39 | 0.056074 | 0 |
| 20 | 2 | 39 | 0.056074 | 0 |
| 1 | 16 | 38 | 0.054636 | 0 |
| 1 | 8 | 37 | 0.053198 | 0 |
| 5 | 16 | 37 | 0.053198 | 0 |
| 10 | 12 | 37 | 0.053198 | 0 |
| 18 | 2 | 37 | 0.053198 | 0 |
| 2 | 18 | 36 | 0.051761 | 0 |
| 2 | 20 | 36 | 0.051761 | 0 |
| 8 | 10 | 36 | 0.051761 | 0 |
| 10 | 11 | 36 | 0.051761 | 0 |
| 18 | 1 | 36 | 0.051761 | 0 |
| 18 | 3 | 36 | 0.051761 | 0 |
| 6 | 14 | 35 | 0.050323 | 0 |
| 7 | 16 | 35 | 0.050323 | 0 |
| 13 | 1 | 35 | 0.050323 | 0 |
| X | 7 | 35 | 0.050323 | 0 |
| 1 | 18 | 34 | 0.048885 | 0 |

| | | | | |
|---|---|---|---|---|
| 1 | 20 | 34 | 0.048885 | 0 |
| 4 | 10 | 34 | 0.048885 | 0 |
| 4 | 12 | 34 | 0.048885 | 0 |
| 4 | 16 | 34 | 0.048885 | 0 |
| 9 | 6 | 34 | 0.048885 | 0 |
| 17 | 1 | 34 | 0.048885 | 0 |
| 17 | 3 | 34 | 0.048885 | 0 |
| 12 | 10 | 33 | 0.047447 | 0 |
| 18 | 6 | 33 | 0.047447 | 0 |
| 3 | 18 | 32 | 0.046009 | 0 |
| 4 | 11 | 32 | 0.046009 | 0 |
| 5 | 4 | 32 | 0.046009 | 0 |
| 5 | 9 | 32 | 0.046009 | 0 |
| 6 | 17 | 32 | 0.046009 | 0 |
| 7 | 11 | 32 | 0.046009 | 0 |
| 8 | 11 | 32 | 0.046009 | 0 |
| 13 | 3 | 32 | 0.046009 | 0 |
| 13 | 5 | 32 | 0.046009 | 0 |
| 16 | 6 | 32 | 0.046009 | 0 |
| 16 | 7 | 32 | 0.046009 | 0 |
| 7 | 12 | 31 | 0.044572 | 0 |
| 13 | 2 | 31 | 0.044572 | 0 |
| 14 | 2 | 31 | 0.044572 | 0 |
| 15 | 7 | 31 | 0.044572 | 0 |
| 18 | 5 | 31 | 0.044572 | 0 |
| 2 | 4 | 30 | 0.043134 | 0 |
| 3 | 14 | 30 | 0.043134 | 0 |
| 3 | 20 | 30 | 0.043134 | 0 |
| 6 | 8 | 30 | 0.043134 | 0 |
| 11 | 10 | 30 | 0.043134 | 0 |
| 12 | 5 | 30 | 0.043134 | 0 |
| 20 | 3 | 30 | 0.043134 | 0 |
| 11 | 12 | 29 | 0.041696 | 0 |
| 11 | 16 | 29 | 0.041696 | 0 |
| 12 | 16 | 29 | 0.041696 | 0 |
| 16 | 2 | 29 | 0.041696 | 0 |
| 3 | 9 | 28 | 0.040258 | 0 |
| 3 | 21 | 28 | 0.040258 | 0 |
| 7 | 10 | 28 | 0.040258 | 0 |
| 7 | 20 | 28 | 0.040258 | 0 |
| 9 | 5 | 28 | 0.040258 | 0 |
| 11 | 8 | 28 | 0.040258 | 0 |
| 11 | 20 | 28 | 0.040258 | 0 |
| 16 | 3 | 28 | 0.040258 | 0 |
| 16 | 12 | 28 | 0.040258 | 0 |
| 19 | 1 | 28 | 0.040258 | 0 |
| X | 12 | 28 | 0.040258 | 0 |

| | | | | |
|---|---|---|---|---|
| 7 | 9 | 27 | 0.03882 | 0 |
| 7 | 14 | 27 | 0.03882 | 0 |
| 15 | 3 | 27 | 0.03882 | 0 |
| 16 | 1 | 27 | 0.03882 | 0 |
| 18 | 11 | 27 | 0.03882 | 0 |
| 20 | 7 | 27 | 0.03882 | 0 |
| X | 8 | 27 | 0.03882 | 0 |
| 3 | 4 | 26 | 0.037383 | 0 |
| 3 | 16 | 26 | 0.037383 | 0 |
| 5 | 18 | 26 | 0.037383 | 0 |
| 6 | 18 | 26 | 0.037383 | 0 |
| 7 | 17 | 26 | 0.037383 | 0 |
| 14 | 11 | 26 | 0.037383 | 0 |
| 16 | 5 | 26 | 0.037383 | 0 |
| 17 | 7 | 26 | 0.037383 | 0 |
| 17 | 12 | 26 | 0.037383 | 0 |
| X | 16 | 26 | 0.037383 | 0 |
| 1 | 21 | 25 | 0.035945 | 0 |
| 4 | 18 | 25 | 0.035945 | 0 |
| 7 | 4 | 25 | 0.035945 | 0 |
| 7 | 18 | 25 | 0.035945 | 0 |
| 10 | 17 | 25 | 0.035945 | 0 |
| 13 | 6 | 25 | 0.035945 | 0 |
| 14 | 5 | 25 | 0.035945 | 0 |
| 15 | 2 | 25 | 0.035945 | 0 |
| 21 | 1 | 25 | 0.035945 | 0 |
| X | 11 | 25 | 0.035945 | 0 |
| 8 | 12 | 24 | 0.034507 | 0 |
| 9 | 7 | 24 | 0.034507 | 0 |
| 9 | 12 | 24 | 0.034507 | 0 |
| 10 | 9 | 24 | 0.034507 | 0 |
| 13 | 7 | 24 | 0.034507 | 0 |
| 14 | 10 | 24 | 0.034507 | 0 |
| 14 | 12 | 24 | 0.034507 | 0 |
| 17 | 2 | 24 | 0.034507 | 0 |
| X | 18 | 24 | 0.034507 | 0 |
| 3 | 17 | 23 | 0.033069 | 0 |
| 4 | 9 | 23 | 0.033069 | 0 |
| 9 | 16 | 23 | 0.033069 | 0 |
| 12 | 8 | 23 | 0.033069 | 0 |
| 4 | 8 | 22 | 0.031631 | 0 |
| 5 | 14 | 22 | 0.031631 | 0 |
| 5 | 17 | 22 | 0.031631 | 0 |
| 10 | 8 | 22 | 0.031631 | 0 |
| 10 | 18 | 22 | 0.031631 | 0 |
| 10 | 20 | 22 | 0.031631 | 0 |
| 15 | 1 | 22 | 0.031631 | 0 |

| | | | | |
|---|---|---|---|---|
| 17 | 5 | 22 | 0.031631 | 0 |
| 18 | 8 | 22 | 0.031631 | 0 |
| 19 | 2 | 22 | 0.031631 | 0 |
| 5 | 21 | 21 | 0.030194 | 0 |
| 6 | 9 | 21 | 0.030194 | 0 |
| 8 | 18 | 21 | 0.030194 | 0 |
| 11 | 17 | 21 | 0.030194 | 0 |
| 11 | 18 | 21 | 0.030194 | 0 |
| 12 | 14 | 21 | 0.030194 | 0 |
| 13 | 8 | 21 | 0.030194 | 0 |
| 15 | 6 | 21 | 0.030194 | 0 |
| 16 | 10 | 21 | 0.030194 | 0 |
| 18 | 12 | 21 | 0.030194 | 0 |
| 19 | 11 | 21 | 0.030194 | 0 |
| X | 10 | 21 | 0.030194 | 0 |
| 2 | 21 | 20 | 0.028756 | 0 |
| 8 | 14 | 20 | 0.028756 | 0 |
| 9 | 11 | 20 | 0.028756 | 0 |
| 10 | 4 | 20 | 0.028756 | 0 |
| 12 | 4 | 20 | 0.028756 | 0 |
| 13 | 11 | 20 | 0.028756 | 0 |
| 14 | 7 | 20 | 0.028756 | 0 |
| 14 | 16 | 20 | 0.028756 | 0 |
| 15 | 10 | 20 | 0.028756 | 0 |
| 18 | 7 | 20 | 0.028756 | 0 |
| 21 | 3 | 20 | 0.028756 | 0 |
| 1 | 15 | 19 | 0.027318 | 0 |
| 2 | 15 | 19 | 0.027318 | 0 |
| 4 | 14 | 19 | 0.027318 | 0 |
| 6 | 4 | 19 | 0.027318 | 0 |
| 8 | 17 | 19 | 0.027318 | 0 |
| 10 | 16 | 19 | 0.027318 | 0 |
| 12 | 11 | 19 | 0.027318 | 0 |
| 15 | 5 | 19 | 0.027318 | 0 |
| 17 | 8 | 19 | 0.027318 | 0 |
| 19 | 10 | 19 | 0.027318 | 0 |
| 1 | 4 | 18 | 0.02588 | 0 |
| 1 | 19 | 18 | 0.02588 | 0 |
| 4 | 17 | 18 | 0.02588 | 0 |
| 8 | 16 | 18 | 0.02588 | 0 |
| 8 | 20 | 18 | 0.02588 | 0 |
| 9 | 14 | 18 | 0.02588 | 0 |
| 10 | 14 | 18 | 0.02588 | 0 |
| 11 | 4 | 18 | 0.02588 | 0 |
| 12 | 9 | 18 | 0.02588 | 0 |
| 12 | 18 | 18 | 0.02588 | 0 |
| 16 | 8 | 18 | 0.02588 | 0 |

| | | | | |
|---|---|---|---|---|
| 16 | 11 | 18 | 0.02588 | 0 |
| 17 | 10 | 18 | 0.02588 | 0 |
| 18 | 14 | 18 | 0.02588 | 0 |
| X | 17 | 18 | 0.02588 | 0 |
| 6 | 20 | 17 | 0.024442 | 0 |
| 6 | 21 | 17 | 0.024442 | 0 |
| 16 | 9 | 17 | 0.024442 | 0 |
| 17 | 6 | 17 | 0.024442 | 0 |
| 19 | 7 | 17 | 0.024442 | 0 |
| 19 | 9 | 17 | 0.024442 | 0 |
| 20 | 1 | 17 | 0.024442 | 0 |
| 20 | 5 | 17 | 0.024442 | 0 |
| X | 14 | 17 | 0.024442 | 0 |
| 5 | 15 | 16 | 0.023005 | 0 |
| 5 | 20 | 16 | 0.023005 | 0 |
| 7 | 21 | 16 | 0.023005 | 0 |
| 10 | 19 | 16 | 0.023005 | 0 |
| 13 | 4 | 16 | 0.023005 | 0 |
| 14 | 9 | 16 | 0.023005 | 0 |
| 15 | 8 | 16 | 0.023005 | 0 |
| X | 20 | 16 | 0.023005 | 0 |
| 4 | 20 | 15 | 0.021567 | 0 |
| 6 | 15 | 15 | 0.021567 | 0 |
| 7 | 19 | 15 | 0.021567 | 0 |
| 11 | 14 | 15 | 0.021567 | 0 |
| 13 | 16 | 15 | 0.021567 | 0 |
| 16 | 14 | 15 | 0.021567 | 0 |
| 16 | 20 | 15 | 0.021567 | 0 |
| 20 | 6 | 15 | 0.021567 | 0 |
| 20 | 11 | 15 | 0.021567 | 0 |
| 2 | 19 | 14 | 0.020129 | 0 |
| 4 | 19 | 14 | 0.020129 | 0 |
| 7 | 15 | 14 | 0.020129 | 0 |
| 8 | 21 | 14 | 0.020129 | 0 |
| 9 | 18 | 14 | 0.020129 | 0 |
| 12 | 21 | 14 | 0.020129 | 0 |
| 14 | 17 | 14 | 0.020129 | 0 |
| 17 | 11 | 14 | 0.020129 | 0 |
| 17 | 20 | 14 | 0.020129 | 0 |
| 18 | 4 | 14 | 0.020129 | 0 |
| 19 | 3 | 14 | 0.020129 | 0 |
| 20 | 17 | 14 | 0.020129 | 0 |
| X | 9 | 14 | 0.020129 | 0 |
| 8 | 4 | 13 | 0.018691 | 0 |
| 9 | 8 | 13 | 0.018691 | 0 |
| 13 | 18 | 13 | 0.018691 | 0 |
| 14 | 8 | 13 | 0.018691 | 0 |

| | | | | |
|---|---|---|---|---|
| 15 | 9 | 13 | 0.018691 | 0 |
| 15 | 11 | 13 | 0.018691 | 0 |
| 15 | 16 | 13 | 0.018691 | 0 |
| 15 | 17 | 13 | 0.018691 | 0 |
| 17 | 9 | 13 | 0.018691 | 0 |
| 18 | 9 | 13 | 0.018691 | 0 |
| 19 | 5 | 13 | 0.018691 | 0 |
| 19 | 20 | 13 | 0.018691 | 0 |
| 20 | 10 | 13 | 0.018691 | 0 |
| 20 | 14 | 13 | 0.018691 | 0 |
| 21 | 7 | 13 | 0.018691 | 0 |
| 3 | 19 | 12 | 0.017254 | 0 |
| 10 | 21 | 12 | 0.017254 | 0 |
| 11 | 21 | 12 | 0.017254 | 0 |
| 12 | 17 | 12 | 0.017254 | 0 |
| 13 | 10 | 12 | 0.017254 | 0 |
| 13 | 12 | 12 | 0.017254 | 0 |
| 13 | 21 | 12 | 0.017254 | 0 |
| 14 | 20 | 12 | 0.017254 | 0 |
| 15 | 14 | 12 | 0.017254 | 0 |
| 17 | 4 | 12 | 0.017254 | 0 |
| 18 | 10 | 12 | 0.017254 | 0 |
| 20 | 16 | 12 | 0.017254 | 0 |
| 21 | 6 | 12 | 0.017254 | 0 |
| 21 | 11 | 12 | 0.017254 | 0 |
| 22 | 5 | 12 | 0.017254 | 0 |
| 4 | 15 | 11 | 0.015816 | 0 |
| 6 | 13 | 11 | 0.015816 | 0 |
| 9 | 17 | 11 | 0.015816 | 0 |
| 9 | 19 | 11 | 0.015816 | 0 |
| 12 | 20 | 11 | 0.015816 | 0 |
| 14 | 4 | 11 | 0.015816 | 0 |
| 15 | 18 | 11 | 0.015816 | 0 |
| 18 | 17 | 11 | 0.015816 | 0 |
| 18 | 20 | 11 | 0.015816 | 0 |
| 19 | 17 | 11 | 0.015816 | 0 |
| 21 | 5 | 11 | 0.015816 | 0 |
| 21 | 10 | 11 | 0.015816 | 0 |
| 22 | 3 | 11 | 0.015816 | 0 |
| 22 | 14 | 11 | 0.015816 | 0 |
| X | 4 | 11 | 0.015816 | 0 |
| 3 | 22 | 10 | 0.014378 | 0 |
| 5 | 19 | 10 | 0.014378 | 0 |
| 5 | 22 | 10 | 0.014378 | 0 |
| 9 | 21 | 10 | 0.014378 | 0 |
| 10 | 13 | 10 | 0.014378 | 0 |
| 11 | 19 | 10 | 0.014378 | 0 |

| | | | | |
|---|---|---|---|---|
| 13 | 14 | 10 | 0.014378 | 0 |
| 16 | 15 | 10 | 0.014378 | 0 |
| 16 | 17 | 10 | 0.014378 | 0 |
| 16 | 18 | 10 | 0.014378 | 0 |
| 17 | 14 | 10 | 0.014378 | 0 |
| 17 | 16 | 10 | 0.014378 | 0 |
| 19 | 6 | 10 | 0.014378 | 0 |
| 19 | 8 | 10 | 0.014378 | 0 |
| 19 | 16 | 10 | 0.014378 | 0 |
| 20 | 12 | 10 | 0.014378 | 0 |
| 20 | 18 | 10 | 0.014378 | 0 |
| 22 | 2 | 10 | 0.014378 | 0 |
| 1 | X | 9 | 0.01294 | 0 |
| 6 | 19 | 9 | 0.01294 | 0 |
| 6 | 22 | 9 | 0.01294 | 0 |
| 7 | 13 | 9 | 0.01294 | 0 |
| 8 | 9 | 9 | 0.01294 | 0 |
| 10 | 15 | 9 | 0.01294 | 0 |
| 13 | 20 | 9 | 0.01294 | 0 |
| 17 | 21 | 9 | 0.01294 | 0 |
| 18 | 16 | 9 | 0.01294 | 0 |
| 20 | 9 | 9 | 0.01294 | 0 |
| 20 | 19 | 9 | 0.01294 | 0 |
| 21 | 12 | 9 | 0.01294 | 0 |
| 22 | 9 | 9 | 0.01294 | 0 |
| 22 | 11 | 9 | 0.01294 | 0 |
| 3 | 13 | 8 | 0.011502 | 0 |
| 3 | 15 | 8 | 0.011502 | 0 |
| 5 | X | 8 | 0.011502 | 0 |
| 8 | 22 | 8 | 0.011502 | 0 |
| 11 | 22 | 8 | 0.011502 | 0 |
| 12 | 13 | 8 | 0.011502 | 0 |
| 12 | 15 | 8 | 0.011502 | 0 |
| 13 | 9 | 8 | 0.011502 | 0 |
| 19 | 12 | 8 | 0.011502 | 0 |
| 19 | 14 | 8 | 0.011502 | 0 |
| 20 | 8 | 8 | 0.011502 | 0 |
| 21 | 2 | 8 | 0.011502 | 0 |
| 22 | 1 | 8 | 0.011502 | 0 |
| X | 15 | 8 | 0.011502 | 0 |
| 1 | 13 | 7 | 0.010065 | 0 |
| 1 | 22 | 7 | 0.010065 | 0 |
| 2 | 13 | 7 | 0.010065 | 0 |
| 2 | 22 | 7 | 0.010065 | 0 |
| 4 | 21 | 7 | 0.010065 | 0 |
| 8 | X | 7 | 0.010065 | 0 |
| 9 | 4 | 7 | 0.010065 | 0 |

| | | | | |
|---|---|---|---|---|
| 9 | 22 | 7 | 0.010065 | 0 |
| 10 | 22 | 7 | 0.010065 | 0 |
| 11 | 15 | 7 | 0.010065 | 0 |
| 13 | 15 | 7 | 0.010065 | 0 |
| 15 | 12 | 7 | 0.010065 | 0 |
| 15 | 19 | 7 | 0.010065 | 0 |
| 16 | 4 | 7 | 0.010065 | 0 |
| 17 | 19 | 7 | 0.010065 | 0 |
| 19 | 4 | 7 | 0.010065 | 0 |
| 19 | 18 | 7 | 0.010065 | 0 |
| 20 | 4 | 7 | 0.010065 | 0 |
| 21 | 16 | 7 | 0.010065 | 0 |
| 21 | 20 | 7 | 0.010065 | 0 |
| 22 | 10 | 7 | 0.010065 | 0 |
| 22 | 16 | 7 | 0.010065 | 0 |
| X | 21 | 7 | 0.010065 | 0 |
| 5 | 13 | 6 | 0.008627 | 0 |
| 7 | X | 6 | 0.008627 | 0 |
| 8 | 13 | 6 | 0.008627 | 0 |
| 9 | 10 | 6 | 0.008627 | 0 |
| 12 | 19 | 6 | 0.008627 | 0 |
| 13 | 17 | 6 | 0.008627 | 0 |
| 15 | 20 | 6 | 0.008627 | 0 |
| 16 | 19 | 6 | 0.008627 | 0 |
| 17 | 13 | 6 | 0.008627 | 0 |
| 18 | 13 | 6 | 0.008627 | 0 |
| 18 | 15 | 6 | 0.008627 | 0 |
| 18 | 21 | 6 | 0.008627 | 0 |
| 19 | 21 | 6 | 0.008627 | 0 |
| 20 | 22 | 6 | 0.008627 | 0 |
| 22 | 4 | 6 | 0.008627 | 0 |
| 22 | 6 | 6 | 0.008627 | 0 |
| 22 | 17 | 6 | 0.008627 | 0 |
| 22 | 20 | 6 | 0.008627 | 0 |
| 3 | X | 5 | 0.007189 | 0 |
| 9 | 15 | 5 | 0.007189 | 0 |
| 10 | X | 5 | 0.007189 | 0 |
| 13 | 22 | 5 | 0.007189 | 0 |
| 14 | 21 | 5 | 0.007189 | 0 |
| 15 | 4 | 5 | 0.007189 | 0 |
| 16 | 21 | 5 | 0.007189 | 0 |
| 17 | 18 | 5 | 0.007189 | 0 |
| 17 | X | 5 | 0.007189 | 0 |
| 20 | 13 | 5 | 0.007189 | 0 |
| 20 | 21 | 5 | 0.007189 | 0 |
| 21 | 14 | 5 | 0.007189 | 0 |
| 21 | 17 | 5 | 0.007189 | 0 |

| | | | | |
|---|---|---|---|---|
| 22 | 12 | 5 | 0.007189 | 0 |
| X | 13 | 5 | 0.007189 | 0 |
| 2 | X | 4 | 0.005751 | 0 |
| 4 | 13 | 4 | 0.005751 | 0 |
| 4 | 22 | 4 | 0.005751 | 0 |
| 4 | X | 4 | 0.005751 | 0 |
| 8 | 15 | 4 | 0.005751 | 0 |
| 9 | 20 | 4 | 0.005751 | 0 |
| 11 | 13 | 4 | 0.005751 | 0 |
| 11 | X | 4 | 0.005751 | 0 |
| 14 | 13 | 4 | 0.005751 | 0 |
| 14 | 15 | 4 | 0.005751 | 0 |
| 14 | 18 | 4 | 0.005751 | 0 |
| 15 | 21 | 4 | 0.005751 | 0 |
| 16 | 13 | 4 | 0.005751 | 0 |
| 18 | 19 | 4 | 0.005751 | 0 |
| 19 | 13 | 4 | 0.005751 | 0 |
| 20 | 15 | 4 | 0.005751 | 0 |
| 22 | 8 | 4 | 0.005751 | 0 |
| 22 | 18 | 4 | 0.005751 | 0 |
| X | 19 | 4 | 0.005751 | 0 |
| 7 | 22 | 3 | 0.004313 | 0 |
| 9 | X | 3 | 0.004313 | 0 |
| 13 | 19 | 3 | 0.004313 | 0 |
| 14 | 19 | 3 | 0.004313 | 0 |
| 18 | 22 | 3 | 0.004313 | 0 |
| 20 | X | 3 | 0.004313 | 0 |
| 21 | 4 | 3 | 0.004313 | 0 |
| 21 | 8 | 3 | 0.004313 | 0 |
| 21 | 9 | 3 | 0.004313 | 0 |
| 21 | 13 | 3 | 0.004313 | 0 |
| 22 | 7 | 3 | 0.004313 | 0 |
| 22 | 13 | 3 | 0.004313 | 0 |
| X | 22 | 3 | 0.004313 | 0 |
| 6 | X | 2 | 0.002876 | 0 |
| 8 | 19 | 2 | 0.002876 | 0 |
| 9 | 13 | 2 | 0.002876 | 0 |
| 13 | Y | 2 | 0.002876 | 0 |
| 14 | 22 | 2 | 0.002876 | 0 |
| 15 | 13 | 2 | 0.002876 | 0 |
| 15 | 22 | 2 | 0.002876 | 0 |
| 15 | X | 2 | 0.002876 | 0 |
| 16 | 22 | 2 | 0.002876 | 0 |
| 16 | X | 2 | 0.002876 | 0 |
| 17 | 15 | 2 | 0.002876 | 0 |
| 19 | 15 | 2 | 0.002876 | 0 |
| 19 | X | 2 | 0.002876 | 0 |

| | | | | |
|---|---|---|---|---|
| 21 | 15 | 2 | 0.002876 | 0 |
| 21 | 18 | 2 | 0.002876 | 0 |
| 22 | 21 | 2 | 0.002876 | 0 |
| 12 | 22 | 1 | 0.001438 | 0 |
| 12 | X | 1 | 0.001438 | 0 |
| 14 | X | 1 | 0.001438 | 0 |
| 17 | 22 | 1 | 0.001438 | 0 |
| 18 | X | 1 | 0.001438 | 0 |
| 19 | 22 | 1 | 0.001438 | 0 |
| 21 | 19 | 1 | 0.001438 | 0 |
| 21 | 22 | 1 | 0.001438 | 0 |
| 22 | 15 | 1 | 0.001438 | 0 |
| 22 | 19 | 1 | 0.001438 | 0 |
| 22 | X | 1 | 0.001438 | 0 |
| 1 | Y | 0 | 0 | 0 |
| 2 | Y | 0 | 0 | 0 |
| 3 | Y | 0 | 0 | 0 |
| 4 | Y | 0 | 0 | 0 |
| 5 | Y | 0 | 0 | 0 |
| 6 | Y | 0 | 0 | 0 |
| 7 | Y | 0 | 0 | 0 |
| 8 | Y | 0 | 0 | 0 |
| 9 | Y | 0 | 0 | 0 |
| 10 | Y | 0 | 0 | 0 |
| 11 | Y | 0 | 0 | 0 |
| 12 | Y | 0 | 0 | 0 |
| 13 | X | 0 | 0 | 0 |
| 14 | Y | 0 | 0 | 0 |
| 15 | Y | 0 | 0 | 0 |
| 16 | Y | 0 | 0 | 0 |
| 17 | Y | 0 | 0 | 0 |
| 18 | Y | 0 | 0 | 0 |
| 19 | Y | 0 | 0 | 0 |
| 20 | Y | 0 | 0 | 0 |
| 21 | X | 0 | 0 | 0 |
| 21 | Y | 0 | 0 | 0 |
| 22 | Y | 0 | 0 | 0 |
| X | Y | 0 | 0 | 0 |
| Y | 1 | 0 | 0 | 0 |
| Y | 2 | 0 | 0 | 0 |
| Y | 3 | 0 | 0 | 0 |
| Y | 4 | 0 | 0 | 0 |
| Y | 5 | 0 | 0 | 0 |
| Y | 6 | 0 | 0 | 0 |
| Y | 7 | 0 | 0 | 0 |
| Y | 8 | 0 | 0 | 0 |
| Y | 9 | 0 | 0 | 0 |

| | | | | |
|---|---|---|---|---|
| Y | 10 | 0 | 0 | 0 |
| Y | 11 | 0 | 0 | 0 |
| Y | 12 | 0 | 0 | 0 |
| Y | 13 | 0 | 0 | 0 |
| Y | 14 | 0 | 0 | 0 |
| Y | 15 | 0 | 0 | 0 |
| Y | 16 | 0 | 0 | 0 |
| Y | 17 | 0 | 0 | 0 |
| Y | 18 | 0 | 0 | 0 |
| Y | 19 | 0 | 0 | 0 |
| Y | 20 | 0 | 0 | 0 |
| Y | 21 | 0 | 0 | 0 |
| Y | 22 | 0 | 0 | 0 |
| Y | X | 0 | 0 | 0 |
| Y | Y | 0 | 0 | 1 |
