## Supplemental Table 2 for "Retrotransposons facilitate the tissue-specific horizontal transfer of circulating tumor DNA between human cells"

Transposons per contig and the fraction of contigs with transposons in inserted and non-inserted ctDNA fragments

| | Multiple Myeloma | | Pancreatic Cancer | | P value |
|---|---|---|---|---|---|
| | Inserted | Non-inserted | Inserted | Non-inserted | |
| Trasnposon/Contig | 1.5 | 1.08 | 1.8 | 1.3 | <0.001 |
| % Contigs with Transposons | 77 | 68.5 | 81.5 | 74 | <0.001 |
