## Supplemental Table 3 for "Retrotransposons facilitate the tissue-specific horizontal transfer of circulating tumor DNA between human cells"

**Supplemental Table3A. Summary table of the TE family members for all inserted contigs from matching multiple myeloma co-culture experiments.**

| Transposon | Location relative to insertion | Contig position | Insertion position | Transposon position | Transposon orientation | Distance from insertion | blast chr | blast position | blast orientation |
|---|---|---|---|---|---|---|---|---|---|
| AluSx | intersect | 1-2229 | 1616-1669 | 1570-1871 | reverse complement | 0 | chr2 | 86528424 - 86530643 | Plus/Plus |
| L1MB3 | right | 1-2677 | 1616-1669 | 1737-2677 | reverse complement | -68 | chr2 | 86528424 - 86530643 | Plus/Plus |
| L2a | intersect | 1-2331 | 1372-2331 | 941-1454 | direct | 0 | chr14 | 32672352-32670273 | Plus/Minus |
| THE1C | intersect | 1-2519 | 1344-2990 | 2050-2519 | direct | 0 | chr14 | 32672352-32670273 | Plus/Minus |
| AluY | intersect | 1-2656 | 2355-2461 | 2346-2656 | reverse complement | 0 | chr13 | 22562455-22559956 | Plus/Minus |
| ]L1ME4b | intersect | 1-2686 | 2390-2477 | 1767-2686 | reverse complement | 0 | chr13 | 22562455-22559956 | Plus/Minus |
| AluSx | intersect | 1-3772 | 2547-2494 | 1573-1880 | direct | 0 | chr17 | | |
| AluY | intersect | 1-3123 | 419-446 | 153-469 | reverse complement | 0 | chr12 LIM domain only 3 | 58715-55601 | Plus/Minus |
| ]AluY | intersect | 1-1628 | 1327-1433 | 1327-1628 | reverse complement | 0 | chr8 | 15650931-15649499 | Plus/Minus |
| AluY | intersect | 1-2816 | 2445-2709 | 2505-2816 | direct | 0 | chr3 | 156308018-156310801 | Plus/Plus |
| AluJb | intersect | 1-1096 | 277-318 | 1-317 | direct | 0 | chr10 | 103114349-103115218 | Plus/Plus |
| AluSx | intersect | 1-1096 | 227-318 | 1-312 | direct | 0 | chr10 | 103114349-103115218 | Plus/Plus |
| MER11C | intersect | 1-3272 | 1584-3024 | 2188-3272 | reverse complement | 0 | chr14 | 46404852-46401883 | Plus/Plus |
| AluY | intersect | 1-1632 | 213-261 | 1-311 | reverse complement | 0 | chr2 | 178310609-178312037 | Plus/Plus |
| MER11B | left | 1-2915 | 2789-2911 | 1-1235 | reverse complement | 1554 | chr1 | 145617544-145619382 | Plus/Plus |
| AluSx | intersect | 1-1235 | 1031-1111 | 883-1190 | reverse complement | 0 | chr3 | 180870641-180871874 | Plus/Plus |

| | | | | | | | | | |
|---|---|---|---|---|---|---|---|---|---|
| AluY | intersect | 1-1786 | 133-284 | 1-312 | direct | 0 | chr6 | 143000057-143001731 | Plus/Plus |
| AluSx | intersect | 1-4743 | 381-552 | 261-574 | direct | 0 | chr5 | 71047678-71042940 | Plus/Minus |
| AluY | intersect | 1-4743 | 381-552 | 261-574 | direct | 0 | chr5 | 71047678-71042940 | Plus/Minus |
| Tigger3a | right | 1-4764 | 40-210 | 2599-2941 | direct | 2389 | chr5 | 71047678-71042940 | Plus/Minus |

**Supplemental Table3B. Summary table of the TE family members for all inserted contigs from matching pancreatic cancer co-culture experiments.**

| Transposon | Location relative to insertion | Contig position | Insertion position | Transposon position | Transposon orientation | Distance from insertion | blast chr | blast position | blast orientation |
|---|---|---|---|---|---|---|---|---|---|
| AluJb | right | 1-3705 | 51-220 | 3394-3705 | direct | +3174 | chr18 | 23191079-23187483 | Plus/Minus |
| AluSx | right | 1-3705 | 145-220 | 3394-3705 | direct | +3174 | chr18 | 23191079-23187483 | Plus/Minus |
| L1PA15 | right | 1-7716 | 84-279 | 6793-7704 | reverse complement | -6514 | chr4 | 74809282-74804853 | Plus/Minus |
| L1PB4 | right | 1-7716 | 84-279 | 6801-7704 | reverse complement | -6522 | chr4 | 74809282-74804853 | Plus/Minus |
| AluSx | intersect | 1-3368 | 198-336 | 169-483 | direct | 0 | chr20 | 60039432-60036075 | Plus/Minus |
| AluSx | intersect | 1-4730 | 4456-4591 | 4409-4720 | direct | 0 | chr16 | 30075661-30080375 | Plus/Plus |
| MIRc | left | 1-4739 | 4465-4600 | 2056-2259 | reverse complement | +2206 | chr16 | 30075661-30080375 | Plus/Plus |
| AluJb | intersect | 1-3676 | 2619-2651 | 2450-2747 | reverse complement | 0 | chr7 | 78202685-78206359 | Plus/Plus |
| AluSx | intersect | 1-3676 | 2619-2651 | 2450-2747 | reverse complement | 0 | chr7 | 78202685-78206359 | Plus/Plus |
| AluSx | intersect | 1-7197 | 1963-1980 | 1677-1989 | reverse complement | 0 | chr8 | 17791254-17789008 | Plus/Plus |
| AluY | intersect | 1-7197 | 6052-6112 | 5828-6138 | reverse complement | 0 | chr8 | 17791254-17789008 | Plus/Plus |

| | | | | | | | | | |
|---|---|---|---|---|---|---|---|---|---|
| MIRb | left | 1-7240 | 5476-5518 | 4109-4377 | direct | -1099 | chr8 | 17791254-17789008 | Plus/Plus |
| MIRc | left | 1-7205 | 5441-5483 | 4106-4324 | direct | -1117 | chr8 | 17791254-17789008 | Plus/Plus |
| AluSx | intersect | 1-3669 | 3257-3375 | 3270-3567 | direct | 0 | chr3 | 128761943-128759611 | Plus/Plus |
| MLT2B1 | intersect | 1-3763 | 3352-3470 | 2852-3356 | direct | 0 | chr3 | 128761943-128759611 | Plus/Plus |
| AluSx | intersect | 1-7277 | 1421-3369 | 2841-3166 | reverse complement | 0 | chr21 | 39057278-39054368 | Plus/Plus |
| L2a | intersect | 1-7340 | 1421-3654 | 2474-2881 | direct | 0 | chr21 | 39057278-39054368 | Plus/Plus |
| MLT1J2 | intersect | 1-7311 | 1473-3625 | 1167-1529 | direct | 0 | chr21 | 39057278-39054368 | Plus/Plus |
| AluSx | intersect | 1-4351 | 989-1127 | 960-1264 | direct | 0 | chr7 | 22688622-22692178 | Plus/Plus |
| AluY | intersect | 1-4251 | 989-1127 | 960-1264 | direct | 0 | chr7 | 22688622-22692178 | Plus/Plus |
| L1ME4b | right | 1-4422 | 989-1127 | 3057-3720 | direct | +1930 | chr7 | 22688622-22692178 | Plus/Plus |
| MIRb | intersect | 1-9468 | 3869-3929 | 3790-3904 | direct | 0 | chr14 | 101625831-101623094 | Plus/Minus |
| MER41E | right | 1-5966 | 2493-2559 | 5071-5660 | direct | +2512 | chr2 | 240694078-240699987 | Plus/Plus |
| AluSx | intersect | 1-7125 | 2220-2359 | 2191-2502 | direct | 0 | chr9 | 77999115-78001422 | Plus/Plus |
| Mam_R4 | intersect | 1-7466 | 2269-2410 | 1015-4872 | direct | 0 | chr9 | 77999115-78001422 | Plus/Plus |
| MIRb | left | 1-7141 | 2236-2375 | 281-550 | reverse complement | +1686 | chr9 | 77999115-78001422 | Plus/Plus |
| MIRb | right | 1-10415 | 2751-2783 | 3949-4170 | reverse complement | -1166 | chr4 | 155243887-155240822 | Plus/Minus |
