## Supplemental Methods for "Retrotransposons facilitate the tissue-specific horizontal transfer of circulating tumor DNA between human cells"

### Identification of tissue specific inserted transposons- Analysis pipeline

#### a. Approach 1

- **Sequencing raw data QC and preprocessing and de novo assembly**

Sequencing data quality check was performed using FastQC [1] and multiQC tools [2]. Illumina adapters were trimmed using trimmomatic v0.39 [3].

For de novo assembly of ct-DNA reads ABySS [4] de novo assembler was used. Before assembly, ct-DNA samples were 10x depth normalized with bbnorm [5]. The best k-mer size for the assembly was predicted with the KmerGenie tool [5]. In addition, ct-DNA samples used in cell line cultured experiments (772 and P201812-2) were assembled without any read depth normalization.

- **Cluster analysis of ct-DNA assembled contigs**

Contig-level assembly sequences were used for cluster analysis. cd-hit-est-2d was used to select contigs specific for MM and PC [7] with 95% of identity as a threshold. cd-hit-est-2d compared to sequence datasets (db1 and db2) and reports sequences that are not similar in db2 as well as sequences that are similar between db1 and db2. Since we were interested in MM- and PC-specific contigs we performed cd-hit-est-2d twice: first, MM contig assemblies were assigned as db2 (for MM-specific contigs), and then PC contig assemblies were assigned as db2 (for PC-specific contigs).

- **Detection of de novo insertions in cell line samples**

All cell line samples were aligned to the human reference genome(hg38) with bwa-mem software (v 0.7.17) [8]. After the alignment step *de novo* insertions detection was performed with the Pamir tool [9]. Pamir uses "one-end anchor" reads (i.e. one-end is mapped while the other is unmapped around breakpoint location) and orphan reads (read pairs where none of the ends can be mapped to the reference) to characterize the novel sequence contents and their insertion breakpoints. Algorithm steps include de novo assembly, re-alignment, and clustering on mentioned reads to generate contigs for putative novel insertions. Aligned BAM files for cell culture sequences along with the hg38 genome reference were supplied to Pamir. The tool outputs a VCF file with the sequence, location, and length of identified novel insertions.
Selection of cancer-specific insertions

To select cancer type-specific inserts, full-length contigs were converted to BLAST databases. Next insertions identified in the match-, mismatch- and no-culture samples for corresponding cell line samples (MM1S or MIA) were blasted against the corresponding sample full-length assembly contig database (772 for MM1S cells and P201812-2 for MIA cells). An insert was considered cancer-specific if: 1) it was present in the matching co-culture sample, but not in mismatch co-culture and no-culture samples; 2) it was aligned to the corresponding ct-DNA sample database with identity at least 70% (to maximize hits for further processing).

- **Multiple sequence alignment and alignment-based contig reassembly and extension**

To define if the selected unique contigs are present in all samples, we have created a BLAST database for all samples and blasted the unique contigs against each database. The myeloma database consisted of contigs from 5 MM patients, for pancreatic cancer the database consisted of 10 PC patients. Unique contigs from each cancer type were blasted against the corresponding database. Then, we have selected the contigs, with alignment length >= 650bp and BLAST identity ≥90%. As a result, we have defined the unique contig sets, which were present in all samples belonging to one cancer type (MM1S - 14 contigs and MIA - 13 contigs). In the next step, we have aligned contigs in all samples with the corresponding insertion, using MAFFT [10], and constructed the consensus sequences.

- **Short-read alignment-based contig re-assembly**

As the aligned contigs did not fully cover the whole full assembly contigs (772 and P201812-2), we have used consensus sequences, obtained in the previous step, as a reference to align fastq reads on it. The alignment was performed with the bwa-mem algorithm [8] and a final consensus was obtained for each unique contig. Alignment visualization and alignment consensus retrieval were performed with UGENE v41 [11].

- **Transposon identification**

We have used the final consensus obtained in the previous step, to identify repeats/transposons with RepeatMasker (RM) rmblast [12]. Then we aligned hit repeats/TE sequences, unique contigs for all samples, and insertion sequence with MAFFT. Mutations in TE sequences were identified by comparison of nucleotides at each position in the multiple sequence alignment with the “Biostrings” R package (version 4.1). We considered a position to contain mutation if the substitution was present in contigs of all samples. In the case of ambiguous nucleotides in contigs introduced by short read alignment (putative heterozygosity in the sample), non-matched nucleotide was considered as a mutation if it was present in all samples. Finally, based on the identified mutations, we have constructed the mutated transposon sequences for each unique contigs (for MM1S and MIA datasets).

- **Transposon expression variability analysis**

Single-end RNA-seq data for MM (n=60) and PC (n=22) samples were processed with the STAR aligner and TEtranscripts pipeline [13]. hg38 and Gencode and TE-specific curated annotations were used as reference genome and gene/TE annotations, respectively. Raw counts were processed using the DESeq2 R package [14]. Low count genes and TEs (less than 1000 read in total for all samples) were removed. Counts across samples were normalized for library sizes and log-transformed using 'regularized log' transformation. Expression variability (EV) of TEs and genes (probes) was estimated using the previously described method [15, 16]. EV estimation allows for estimating the expression-independent variability. We have used two parameter sets: 1) median/median absolute deviation (MAD), and 2) mean/standard deviation(SD). First, a bootstrapped estimate of the MAD/SD of each probe was calculated using 1000 bootstrap replicates. Next, the expected MAD/SD as a function of median/mean was calculated using local polynomial regression (*loess* R function). Expression variability (EV) was calculated as the difference between the bootstrapped MAD/SD and the expected MAD/SD for each median/mean expression level. EV, in this case, shows the level of variability for a particular probe compared to other probes with the same expression values. The empirical distribution of EV (ecdf R function) was used to estimate the significance of variability ($P_{EV}$).

First, we calculated magnitude-independent expression-variability for all probes (genes and TEs). Next, we regressed expression values - variance using local polynomial regression to estimate expected variability and estimated the deviance of observed variance for each probe (expression variability, EV) (figure 4). Negative EV values indicate lower variance compared to other probes with a similar expression, and positive EV values show a higher variance. Empirical distribution of EV was also used to calculate the lower-tail p values for each probe. Following thresholds have been defined for highly expressed invariant Tes (HEI Tes): log-transformed Mean/Median expression $\geq 10$ (corresponds to 1024 reads), $P_{EV} < 0.1$.

#### Identification of highly expressed and invariant TEs

##### 1. Data preprocessing

Rowcounts were processed using **DESeq2** R package. Low count genes and TEs (less than 1000 read in total for all samples) were removed. After pre-filtering 1021 TEs and 20251 genes remained. Counts across samples were normalized for library sizes and log-transformed using 'regularized log' transformation. Batch normalization was performed on log-transformed data with *ComBat* function from **sva** R package (figure 1-3).

Before adjustment

After adjustment

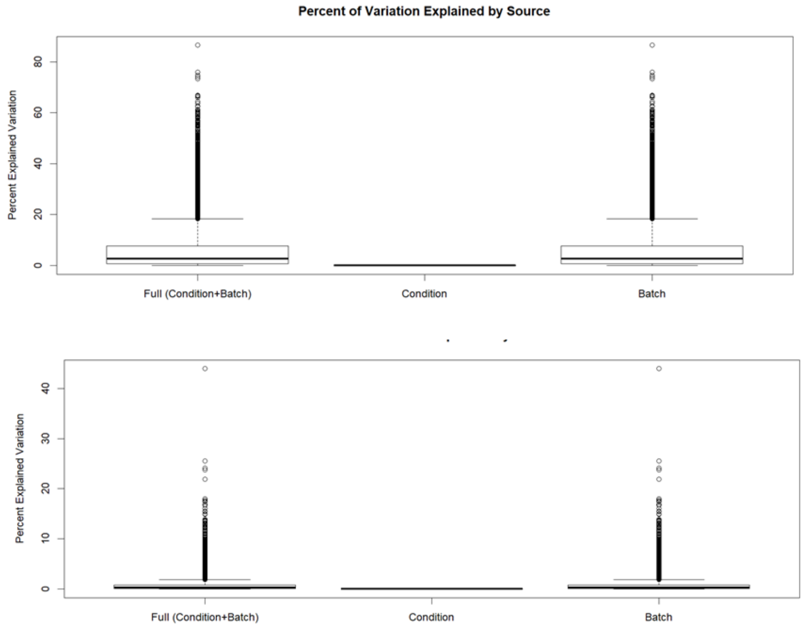


Figure 1. Batch-related variance before and after ComBat transformation (21272 genes + TEs).

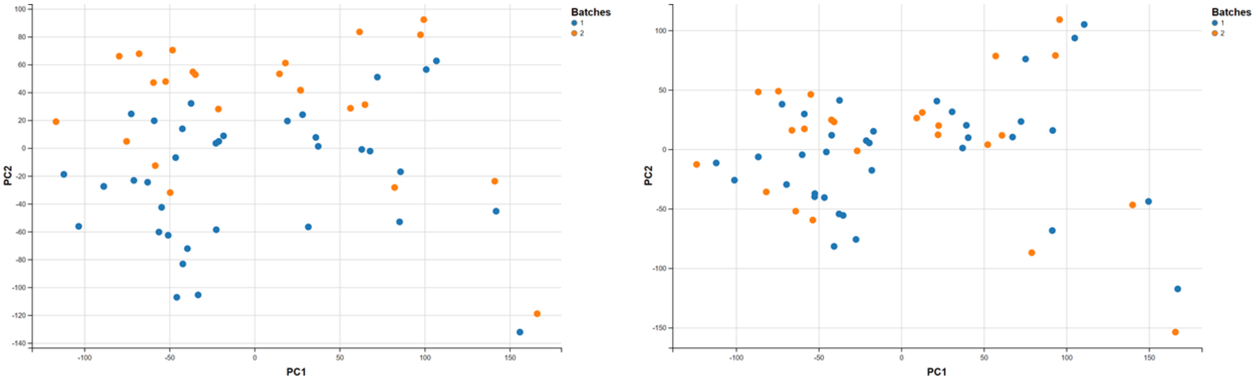


Before adjustment

After adjustment

Figure 2. PCA plots before and after ComBat transformation (21272 genes + TEs)

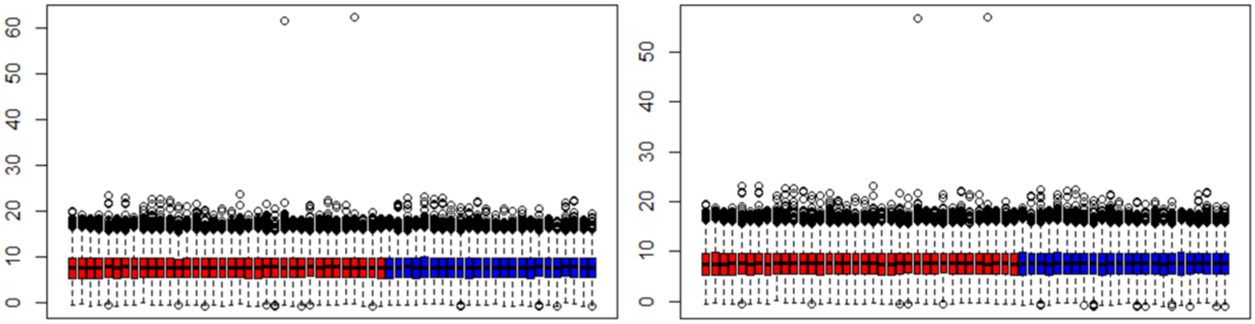

Before adjustment

After adjustment

Figure 3. Expression boxplot before and after ComBat transformation (21272 genes + TEs)

##### 2. Expression variability estimation

Expression variability (EV) of TEs and genes (probes) was estimated using the previously described method [1, 2]. EV estimation allows for estimating the expression-independent variability. We have used two parameter sets: 1) median/median absolute deviation (MAD), and 2) mean/standard deviation(SD). First, a bootstrapped estimate of the MAD/SD of each probe was calculated using 1000 bootstrap replicates. Next, expected MAD/SD as a function of median/mean was calculated using local polynomial regression (*loess* R function) (figure 4).

| A | B |
|---|---|
| 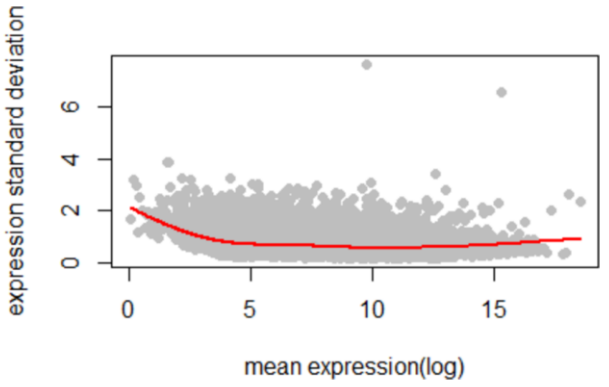  | 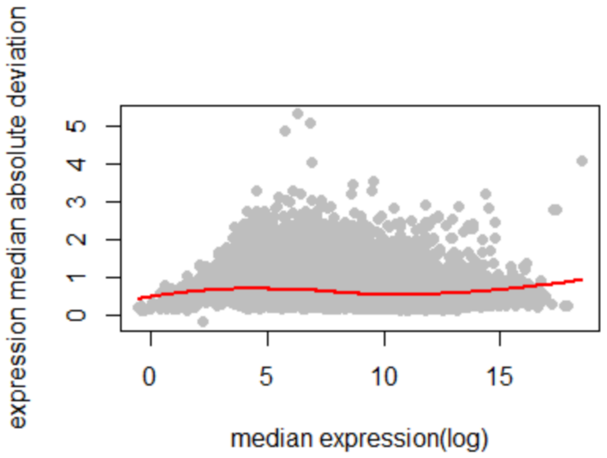  |

Figure 4. Bootstrapped (grey dots) and regressed variation (red line) for mean/SD (A) and median/MAD (B).

Expression variability (EV) was calculated as the difference between the bootstrapped MAD/SD and the expected MAD/SD for each median/mean expression level. EV in this case shown the level of variability for a particular probe compared to other probes with the same expression values. The empirical distribution of EV (*ecdf* R function) was used to estimate the significance of variability.

| A | B |
|---|---|

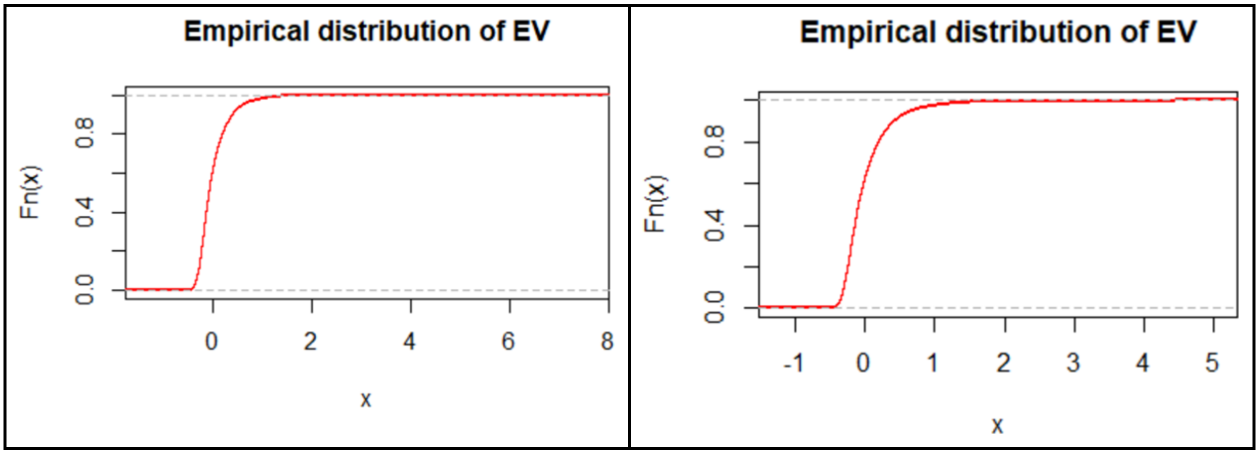


Figure 5. Empirical distribution of expression variability for mean/SD (A) and median/MAD (B).

##### 3. Identification of highly expressed invariant (HEI) TEs (with Mean/SD)

First, we calculated magnitude-independent expression-variability for all probes (genes and TEs). Next, we regressed expression values - variance using local polynomial regression to estimate expected variability and estimated the deviance of observed variance for each probe (expression variability, EV) (figure 4). Negative EV values indicate lower variance compared to other probes with a similar expression, and positive EV values show the higher variance. Empirical distribution of EV was also used to calculate the lower-tail p values for each probe.

Following thresholds have been defined for HEI TEs: log-transformed Mean/Median expression ≥ 10 (corresponds to 1024 reads), $P_{EV} < 0.05$. Only one TE satisfied these criteria. With less stringent criteria $P_{EV} \leq 0.1$, 6 additional TEs were identified (Table 1, Figure 6).

Table 1. HEI TEs (Mean-SD EV assessment).

| Rank (mean expression) | Rank (EV p value) | Probes | Mean | Obs.SD | Exp.SD | EV | EV.pvalue |
|---|---|---|---|---|---|---|---|
| 6 | 19 | MIRb:MIR:SINE | 16.364 | 0.390 | 0.793 | -0.403 | 0.020 |
| 7 | 55 | L2a:L2:LINE | 16.224 | 0.444 | 0.783 | -0.339 | 0.061 |
| 12 | 52 | MIR:MIR:SINE | 15.846 | 0.417 | 0.759 | -0.342 | 0.059 |
| 13 | 94 | L2c:L2:LINE | 15.778 | 0.450 | 0.755 | -0.305 | 0.099 |
| 14 | 59 | L2b:L2:LINE | 15.609 | 0.414 | 0.745 | -0.332 | 0.068 |
| 17 | 71 | MIRc:MIR:SINE | 15.157 | 0.400 | 0.720 | -0.320 | 0.080 |
| 18 | 50 | MIR3:MIR:SINE | 15.151 | 0.375 | 0.720 | -0.344 | 0.057 |

* Ranks are calculated across all genes & TEs. Full data is available in ‘0.rlogcpm.sd.xls file’.

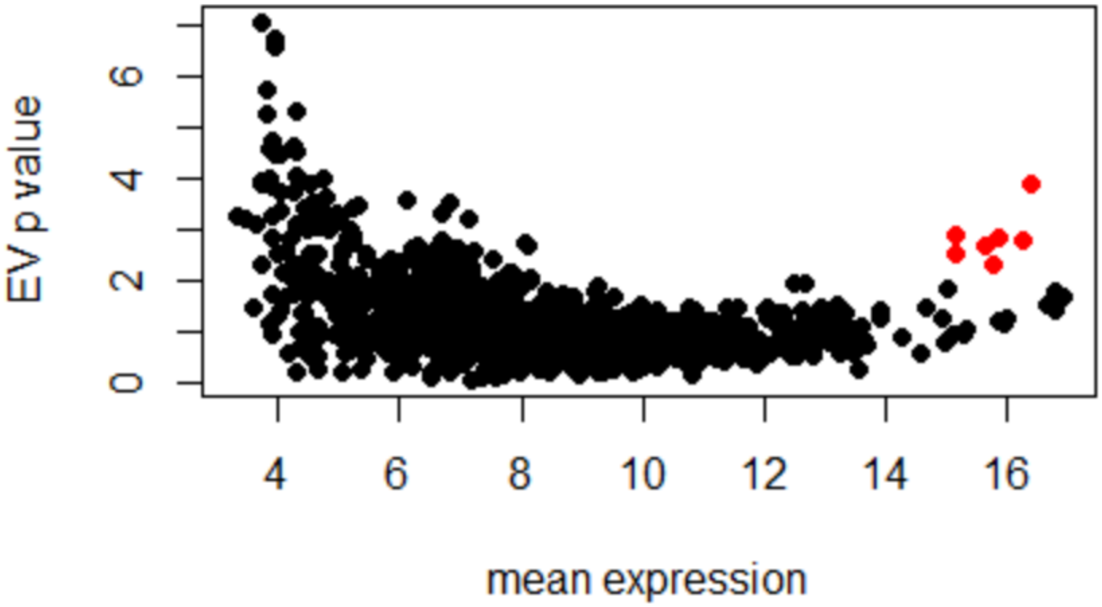


Figure 6. Expression variability - mean expression plot. Red dots - highly expressed invariant TEs (mean log-transformed expression ≥ 10, $P_{EV}$ ≤ 0.1).

##### 4. Identification of highly expressed invariant (HEI) TEs (with Median/MAD)

Median-MAD EV estimation produced a larger set of highly expressed and invariant TEs compared to Mean-SD estimation. In addition to 7 HEI TEs (table 1), three additional TEs were found (table 2, figure 7).

Table 2. HEI TEs (Median-MAD EV assessment).

| Rank (median expression) | Rank (EV p value) | Probes | Median | Obs.MAD | Exp.MAD | EV | EV.pvalue |
|---|---|---|---|---|---|---|---|
| 3 | 53 | AluY:Alu:SINE | 16.68 | 0.41 | 0.78 | -0.38 | 0.030 |
| 6 | 1 | MIRb:MIR:SINE | 16.42 | 0.29 | 0.76 | -0.47 | 0.001 |
| 7 | 101 | L2a:L2:LINE | 16.26 | 0.41 | 0.75 | -0.34 | 0.057 |
| 9 | 47 | MIR:MIR:SINE | 15.92 | 0.35 | 0.73 | -0.38 | 0.028 |
| 12 | 85 | L2c:L2:LINE | 15.81 | 0.37 | 0.72 | -0.35 | 0.050 |
| 14 | 15 | L2b:L2:LINE | 15.66 | 0.29 | 0.71 | -0.42 | 0.011 |
| 15 | 21 | MIRc:MIR:SINE | 15.25 | 0.28 | 0.68 | -0.41 | 0.014 |
| 16 | 22 | MIR3:MIR:SINE | 15.24 | 0.28 | 0.68 | -0.41 | 0.015 |
| 27 | 156 | L1M5:L1:LINE | 13.85 | 0.30 | 0.61 | -0.31 | 0.093 |
| 75 | 102 | L1MC3:L1:LINE | 12.21 | 0.22 | 0.56 | -0.34 | 0.057 |

* Ranks are calculated across all genes & TEs. Full data is available in ‘0.rlogcpm.mad.xls file’.

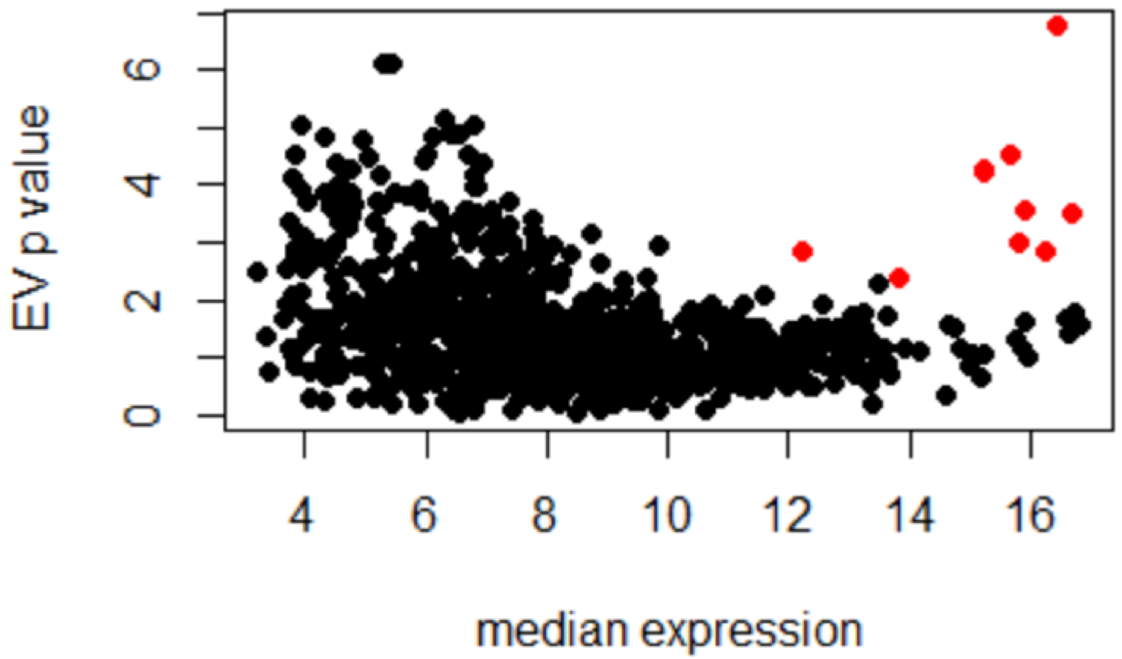


Figure 7. Expression variability - median expression plot. Red dots - highly expressed invariant TEs (median log-transformed expression ≥ 10, $P_{EV}$ ≤ 0.1).

In order to access the uniformity of count and log-transformed data we generated barplots of count and log-transformed expression values for each identified TE (file: 0.HEI_TE_barplot.pdf). It could be seen that these TEs demonstrate little variability of expression across samples.

5. **Identification of highly expressed invariant (HEI) TEs (with Median/MAD)**

Heatmap clustering confirmed that selected TEs have high expression values (figure 8).

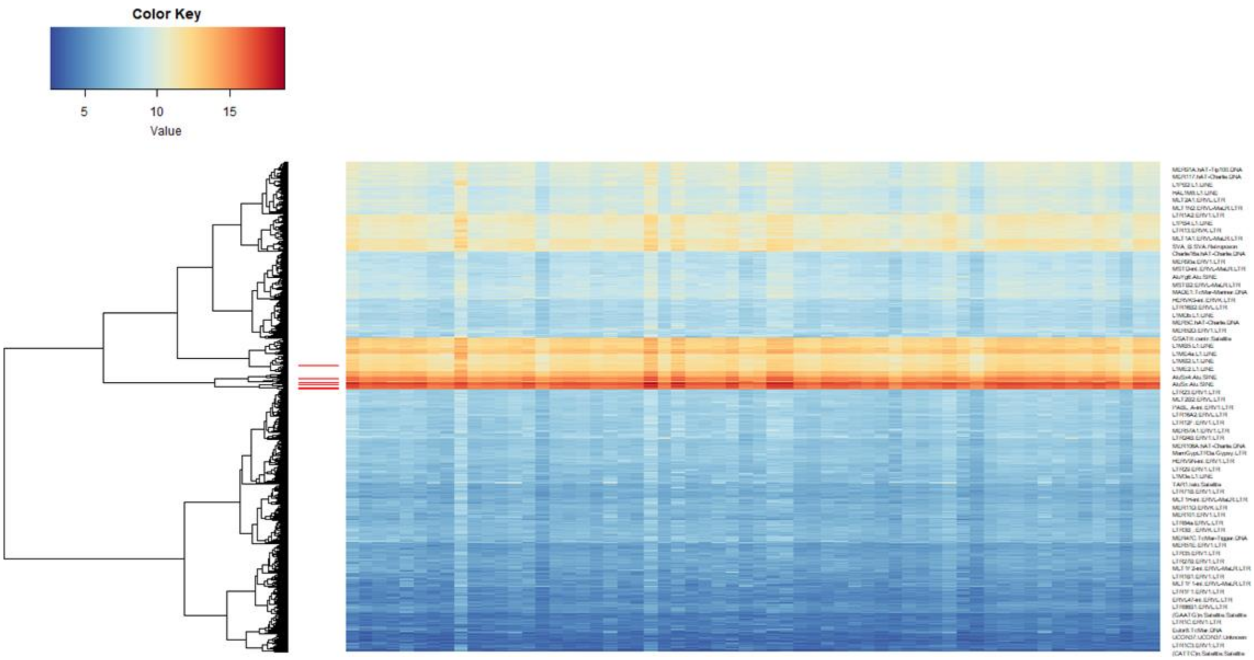

Figure 8. Heatmap clustering of TEs expression. Red stripes indicate identified HEI TEs.

Next, we generated random datasets of TEs by 100 bootstrap replicates for each gene and performed clustering for each dataset (file: 0.clustering.pdf). In the majority of datasets the HEI TEs cluster together.

**Accompanying files:**

Figures.pptx - Separate slides for figures 1-8
0.rlogcpm.mad.xls - meadian/MAD EV results
0.rlogcpm.sd.xls - mean/SD EV results
0.HEI_TE_barplot.pdf - count and log expression barplots across samples for HEI TEs
0.cluster.pdf - Clustering in bootstrapped TE datasets

#### Synthetic Transposon Design

A polynucleotide comprising sequences corresponding to the transposon that contained mutations shared by all the MM samples was generated by Integrated DNA Technologies, Inc (IDT). The majority of the transposons used were selected because of their ability to be generate as gBlock except for AluSq that required to be produced in 2 blocks to overcome interference from the poly T segment. Additionally, in the AluSq an EcoRI complementary site was added at the end to facilitate ligation to Cytomegalovirus-green fluorescent protein (CMV-mCherry) or -herpes simplex virus thymidine kinase (HSV-TK) linearized vectors.

AlusSq
5’ACCCGGCCTTGGACACGCCATTTTCAACTCCGTGGTGCGTTTTTTTTTTTTTTTTTTT
TTTTTGTAATGGAGTTTTGCTCTTGTTGCCCAGGATGGAGTGCAAGGGATCTTGGCTC
ACCACAGCCTCTGCCTCCTGGGTTCAAGTGATTCTTCTGCCTCAGCCTCCCAAGTAG
CTGGGATTATAAGCACCCACCACCACGCCCAGCTAATTTTGTATTTTTTAGAAGAGA
TGGAGTTTCTCCAGTTGGCCAGGATGGTCTGTATATCCTGACCTCATGATCTGCCCAC
CA 3’.
